# Demographic history and genetic structure in pre-Hispanic Central Mexico

**DOI:** 10.1101/2022.06.19.496730

**Authors:** Viridiana Villa-Islas, Alan Izarraras-Gomez, Maximilian Larena, Elizabeth Mejía Perez Campos, Marcela Sandoval-Velasco, Juan Esteban Rodríguez-Rodríguez, Miriam Bravo-Lopez, Barbara Moguel, Rosa Fregel, Jazeps Medina Tretmanis, David Alberto Velázquez-Ramírez, Alberto Herrera-Muñóz, Karla Sandoval, Maria A. Nieves-Colón, Gabriela Zepeda, Fernando A Villanea, Eugenia Fernández Villanueva Medina, Ramiro Aguayo-Haro, Cristina Valdiosera, Alexander Ioannidis, Andrés Moreno-Estrada, Flora Jay, Emilia Huerta-Sanchez, Federico Sánchez-Quinto, María C. Ávila-Arcos

## Abstract

Aridoamerica and Mesoamerica are two distinct cultural areas that hosted numerous pre-Hispanic civilizations between 2,500 BCE and 1,521 CE. The division between these regions shifted southward due to severe droughts ca. 1,100 years ago, allegedly driving demographic changes and population replacement in some sites in central Mexico. Here, we present shotgun genome-wide data from 12 individuals and 26 mitochondrial genomes from eight pre-Hispanic archaeological sites across Mexico, including two at the shifting border of Aridoamerica and Mesoamerica. We find population continuity spanning the climate change episode and a broad preservation of the genetic structure across present-day Mexico for the last 2,300 years. Lastly, we identify a contribution to pre-Hispanic populations of northern and central Mexico from an ancient unsampled ‘ghost’ population.

## Introduction

Before European colonization, the territory comprised by present-day Mexico was home to numerous civilizations that occupied two main cultural areas: Aridoamerica in northern Mexico, inhabited mainly by hunter-gatherers, and Mesoamerica in central and southern Mexico (Fig. 1), where some of the largest agriculture-based civilizations flourished between 2,500 BCE and 1,521 CE (*1, 2*). The distinction between Aridoamerica and Mesoamerica is based on the cultural characteristics and subsistence strategies of the peoples that inhabited them, as well as the ecological features of each region (*3, 4*). Archaeological evidence indicates that the border between these two areas shifted southward between 900 and 1,300 CE following multi-decadal droughts (*5*). This period is also known as the Medieval Warm Period in other regions of the world (*6*). Moreover, this climate change period allegedly led to population replacements in the Northern Frontier of Mesoamerica (NFM) by semi-nomadic hunter-gatherers (“Chichimecas”) from Aridoamerica (*7*), and the fall of some pre-Hispanic societies and the abandonment of Mesoamerican cities in central (*5, 8*) and southeast (*8–10*) Mexico.

**Fig.1.**
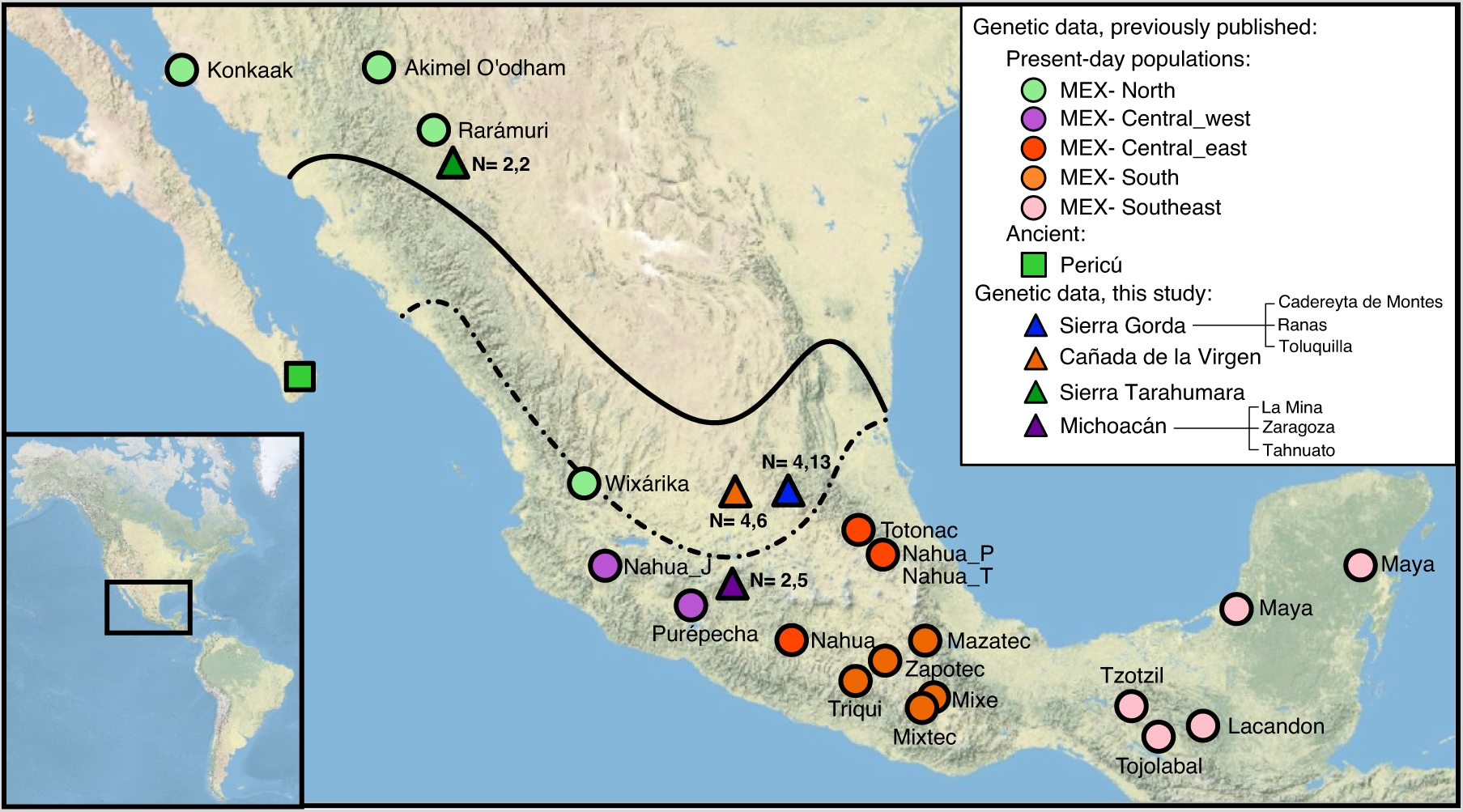
Map of Mexico indicating the location of the pre-Hispanic sites analyzed here and in previous studies (*14, 15*) and, the approximate locations of present-day Indigenous populations sampled in previous studies (*14, 50, 54, 110*). The continuous line indicates the border between Aridoamerica (northern) and Mesoamerica (central and southern) during the ninth century, while dashed line indicates this border in the sixteenth century based on (*2*). Numbers next to archaeological sites from this study indicate the number of whole-genome data and mitochondrial genomes recovered per site (separated by a comma).

Evidence for the population replacement at the NFM comes solely from the archaeological record, and whether this change was the product of migration or acculturation has been debated by archaeologists for years (*8, 11–13*). Studying the genetic variation of ancient populations that span this period of climate change across these two regions is thus necessary to reveal with more precision the regional population dynamics in response to this drastic environmental change. However, ancient genomic data for pre-Hispanic populations from Mexico is very limited, with only two studies reporting eleven low-depth genomes (<0.3x) for a handful of individuals restricted to northern Mexico (*14, 15*), and no available genomes from central and southern Mexico.

Here, we generate the most extensive set of complete mitogenomes and shotgun genome-wide data to date from pre-Hispanic individuals from Mexico spanning a time transect of 830 years: 7 genomes with coverage ranging from 0.01x to 0.14x, and five genomes with coverage ranging from 0.5x to 4.7x depth. Moreover, we sequenced 26 mitogenomes that span a time transect of 1,600 years. We include three previously published ancient genomes from northern Mexico as well as nine ancient mitogenomes yielding a total of 15 ancient genomes and 35 ancient mitogenomes from the Mexican territory. We analyze our data jointly with publicly available datasets of other ancient Native Americans outside of Mexico and present-day Indigenous populations from Mexico. This large compendium of ancient genomic data allows us to study the pre-Hispanic genetic structure in the territory occupied by Mexico, answering longstanding questions regarding population dynamics at the NFM, disentangling the complex structure of central Mexico populations, and revealing a previously unknown ancient contribution of an unsampled genetic linage to Mexican populations.

## Results

### Ancient genomic data from pre-Hispanic individuals from Northern and Central Mexico

To gain insights into the population dynamics of Mexico in the last 2,300 years, we investigated the genetic diversity, population structure, and demographic history of the populations inhabiting the Mexican territory prior to European colonization. We screened archeological samples from 39 pre-Hispanic individuals. Thirty-seven out of the 39 pre-Hispanic individuals were sampled from sites in Mesoamerica. These samples were taken from individuals located in seven archaeological sites from central Mexico (n=37). The central Mexico sites include three in the Sierra Gorda (SG) in Querétaro state, namely Toluquilla (TOL, n=9), Ranas (R, n=3) and a cave in Cadereyta de Montes (CCM, n=1); Cañada de la Virgen (CdV) in Guanajuato state (n=16), and the Zaragoza (n=4), Tanhuato (n=1) and La Mina (n=3) sites in Michoacán state (Mich). Furthermore, we produced additional genome-wide data for two mummies from the Sierra Tarahumara, northern Mexico (Aridoamerica) (Table S1), which were sequenced at lower depths in a previous study (*14*) (Fig. 1).

For each sample, we evaluated the extent of endogenous ancient DNA (aDNA) via shotgun sequencing. Out of the 39 samples, only three had endogenous content above 10% (13 – 39%) (Table S2). We then generated additional sequence data for these three samples using shotgun sequencing, while for the remaining libraries, we performed whole-genome or mitochondrial DNA (mtDNA) capture (*16, 17*) (Table S2). For all of these, we verified the presence of the characteristic misincorporation patterns of aDNA using mapDamage (*18*) (Fig. S1-S4). We obtained informative amounts of genome-wide data for 12 individuals (0.01 – 4.7 x). Ten were from Mesoamerica; SG (n=4), CdV (n=4), and Michoacán (n=2), and two from the Sierra Tarahumara in Aridoamerica (Table S1). We integrated these data with previously reported low-depth genomes (0.09 – 0.3 x) from three ancient Pericúes from Mexico (Aridoamerica) reported in (*14, 15*), resulting in a total of 15 pre-Hispanic individuals from Mexico for which we had genome-wide data. Our analysis also include ancient genomic data from previous studies for 21 individuals from across the Americas (Table S3)(*14, 15, 19–24*). Additionally, we reconstructed the mitochondrial genomes for 26 individuals across Mexico (5.7 – 1284.8x), spanning 1,600 years (320 BCE to 1,351 CE). Twenty-four from Mesoamerica; SG (n=13), CdV (n=6), and Michoacán (n=5), and 2 individuals from Aridoamerica; Sierra Tarahumara (n=2) (*14*) (Table S1 and S2). These mitochondrial genomes were analyzed along with previously published mitochondrial genomes from Mexico (*14, 15, 25*) (Table S4).

Individuals’ names were stated according to burial and individual numbers provided by archaeologists. Additionally, we added two labels, one consisting of 1-3 letters that refer to the archaeological site and the other corresponding to one letter that represents the period of the individuals: a letter ‘b’ for before the longstanding droughts and ‘a’ for those from the period during or after the longstanding droughts. An underscore character ‘_’ separates these three pieces of information. Throughout the text we refer to all individuals using these IDs.

### Genetic structure and diversity in uniparental markers

All individuals carried one of the mtDNA haplogroups found in Indigenous populations of the Americas: A (n=10), B (n=8), C (n=4), and D (n=4) (*26*) (Table S1) (see supplementary text). The pre-Hispanic mtDNA haplogroup distribution resembles the one observed in present-day Mexico (*27–29*), showing an overall continuity of matrilineal genetic structure for at least 2,300 years. We observed haplogroup C to be predominant in Aridoamerica in the Sierra Tarahumara, with decreasing frequency toward Mesoamerica. This gradient has been previously observed in ancient and contemporary populations from Mexico (*25, 30–36*). Specifically for central Mexico, we found the presence of all four autochthonous haplogroups, similar to previous observations on ancient and present-day populations inhabiting this region (*27–31, 37*), which highlights the higher genetic diversity and heterogeneity in this particular area (*38, 39*), and which is in agreement with larger civilizations and greater population mobility than in Aridoamerica. Furthermore, we detected unreported variants in the mtDNA PhyloTree (*40*) for seven individuals assigned to the sub-haplogroups A2d (n=4), B2c (n=2), and B2l (n=1) .The absence of these variants could reflect a poor sampling of Native American mitochondrial genomes in the PhyloTree database or a loss of these haplotypes as a consequence of drift that includes the effect of a population decline following European colonization as reported in other Native American populations (*41, 42*). Accordingly, we propose adding these new unreported variants to the PhyloTree database (Table S5).

Notably, in SG we found the sub-haplogroup A2d in six out of 13 pre-Hispanic individuals, including the oldest in our dataset, P_CCM_b (320 BCE) (Table S1). Out of this six A2d individuals, four belong to the pre-drought period according to dating based either on C14 or the archaeological context, and two to the drought periods. A haplotype median-joining network for the haplogroup A2 shows a clustering of the SG individuals carrying the sub-haplogroup A2d despite a time transect of 1,480 years (Fig. 2A). This observation supports a population continuity, at least of the female population despite the severe droughts in 900 – 1,300 CE in this region of the NFM. For the other haplogroups B, C, and D, when individuals from the same site share the same haplogroup their sub-haplogroups tend to cluster together (Fig. S5).

**Fig. 2.**
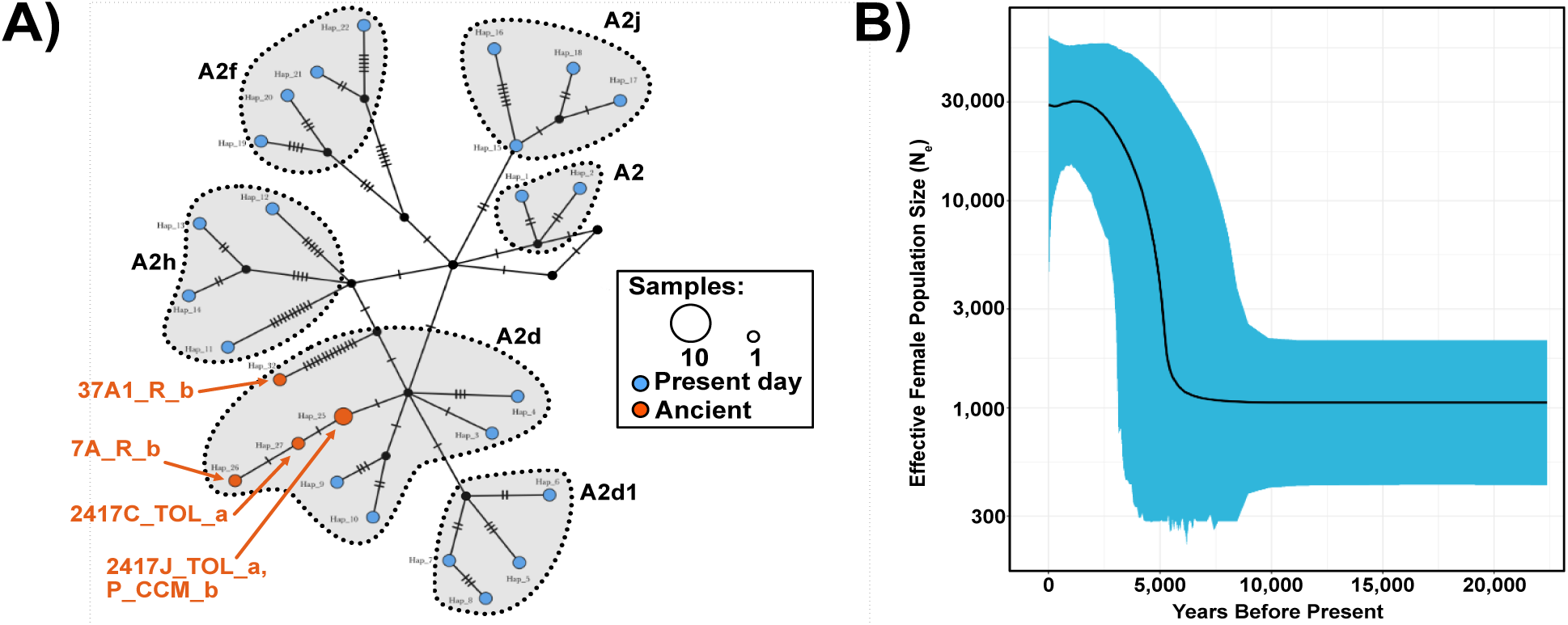
Haplotype network analysis for haplogroup A including ancient and modern mtDNA, and Extended Bayesian Skyline Plot (EBSP) for all present-day mtDNA haplogroups from Mexico. A) Haplotype network of mitochondrial haplogroup A including present-day and ancient mitochondrial haplotypes from Mexico. Individuals from SG within haplogroup A2d cluster together independently of their period. B) EBSP for all present-day mtDNA haplogroups from Mexico. EBSP shows a constant population size after the peopling of the Americas, followed by a population expansion 15,000 years ago. After that, it shows a population decline starting slowly at 2,000 years ago and accelerating at 0.5 years ago when contact with Europeans occurred. The Median is shown as a black line, and the credibility interval of 95% is shown in blue.

To estimate the past female effective population size, we merged our dataset of pre-Hispanic mtDNA sequences (n=16, selected after alignments, see methods) with 164 publicly available ancient (n=9) and modern (n=155) mtDNA sequences from Mexico (Table S4 and S6). We generated an extended Bayesian Skyline Plot (EBSP) for all present-day haplogroups, for all pre-Hispanic haplogroups, for each haplogroup (present-day + pre-Hispanic), and for all haplogroups together merging present-day and pre-Hispanic (Fig. 2B and Fig. S6A-F). We ran the EBSP twice to observe the basic statistics of the runs, and both showed similar results (we report here the plots of the first replicate) (Table S7), we plotted the median value (black line) for all the EBS. The EBSP of all the present-day haplogroups together reconstructs a constant population size after the peopling of the Americas until 5,000 years ago, when a population expansion is observed. This expansion is followed by a population decline starting 2,000 years ago and becoming more pronounced 500 years ago (Fig. 2B). A similar trend is observed in the median value of the EBSP for haplogroups B and C (from modern + pre-Hispanic) but with higher variance in the 95% CI (Figs. S6C-S6D). While the EBSP from all the pre-Hispanic haplogroups together, haplogroup A (modern + pre-Hispanic), haplogroup D (modern + pre-Hispanic) and all haplogroups together (modern + pre-Hispanic) do not show any population decline in the median values observed when present-day mtDNA is used for the analysis (Fig. S6A, S6B, S6E and S6F). Similar demographic scenarios of population decline as in Fig.2B have been observed in other regions of the Americas colonized by Europeans in the 16th century, such as Puerto Rico and Peru (*41, 43*). However, we found a larger population size and a less drastic decline in the effective female population size than in these two regions (based on the 95% CI) (*41, 43*). This finding is consistent with a previous study that reconstructed the Ne based on exome and whole-genome data from present-day populations and found that in Mexico the Ne was higher than in Puerto Rico and Peru (*44*).

Determination of biological sex was made using the tool reported in (*45*). This approach computes the proportion of reads mapped to the Y chromosome with respect to the reads mapping to the X chromosome and Y chromosomes (Ry). According to the method, Ry > 0.075 corresponds to XY, while Ry < 0.016 corresponds to XX. We could assign a sex for 16/39 individuals (females=3, males= 13). For another four individuals, although the Ry values were above the required threshold for XX assignment, they were very close (E2_Mich_b: 0.0218, E4_Mich_b: 0.022, E7_CdV_b: 0.0339 and E19_CdV_b: 0.0312). Sex assignment for these individuals was possible after applying a filter with pmdtools (*46*) with thresholds of 0 – 3 (see Table S1), even though contamination estimates based on the mitochondrial genome were <3%. After pmdtools, all these four individuals were assigned as XX (Table S1). Interestingly, we performed whole-genome in-solution capture for these female individuals. According to a previous study, captured libraries tend to be enriched in pseudo-autosomal and repetitive regions, due to an artifact caused by the use of DNA of male individuals for preparing the RNA baits, which could lead to a bias in the estimate of the Ry value (*47*).

Likewise, we analyzed pre-Hispanic male sub-lineages on the Y chromosome (Y-DNA). The identification of Y-DNA haplogroups was only possible in five out of the thirteen individuals genetically assigned as males, due to the low coverage (Table S1). All individuals presented the Native American Q lineage as expected and in agreement with previous studies on ancient and present-day Indigenous Mexican individuals (*29, 34, 37, 48–50*). Specifically, we identified the sub-haplogroups Q1a2a1-L54 in the Sierra Tarahumara (n=1) and Ranas (n=1) and Q1a2a1a1a1-M3 in the Sierra Tarahumara (n=1), Toluquilla (n=1) and CdV (n=1), with no apparent differences between Aridoamerica and Mesoamerica (Table S1).

### Autosomal genetic structure in pre-Hispanic Mexico

We performed ADMIXTURE(*51*) and Principal Components Analysis (PCA) (*52, 53*) to visualize the genetic relationship and genetic structure at the autosomal level between the pre-Hispanic individuals and other ancient and present-day Native American and continental populations (Table S8) (*14, 34, 50, 54, 55*).

All pre-Hispanic individuals from Mexico, as well as all ancient individuals from California, Belize and Patagonia clustered together in PCA with present-day Indigenous populations from Mexico (Fig. S7) and share similar genetic composition between themselves as evidenced in ADMIXTURE analysis (Fig. S8). The same analyses were repeated including only the pre-Hispanic individuals and the present-day Indigenous populations from Mexico (*14, 34, 50, 54*) as a reference. When projecting the pre-Hispanic individuals onto the principal component space, these individuals clustered closely with present-day populations from the same geographical region, respectively. In particular, the individual F9_ST_a from northern Mexico clusters closely with present-day Rarámuri and Wixárika Indigenous populations (from north Mexico). Pericúes from Baja California (*14, 15*) cluster closely with Wixárika, central Mexico Nahuas from Jalisco and Purépechas. Regarding the individuals from central Mexico (SG, CdV, and Michoacán), these cluster closely with present-day Indigenous populations from this same region (Nahua from Jalisco, Purépecha, Totonac, Nahua from Puebla, and Nahua) (Fig. 3A). Notably, pre-, and post-mega-drought SG individuals, cluster together with central Mexico present-day populations, contrary to what would be expected if a population replacement from northern Mexico in the Sierra Gorda would have taken place due to the climate change (Fig. 3A). For the ancient individual MOM6_ST_a from northern Mexico, he clustered closely with present-day Indigenous populations from central-west Mexico (Purépecha and Nahuas from Jalisco).

**Fig.3.**
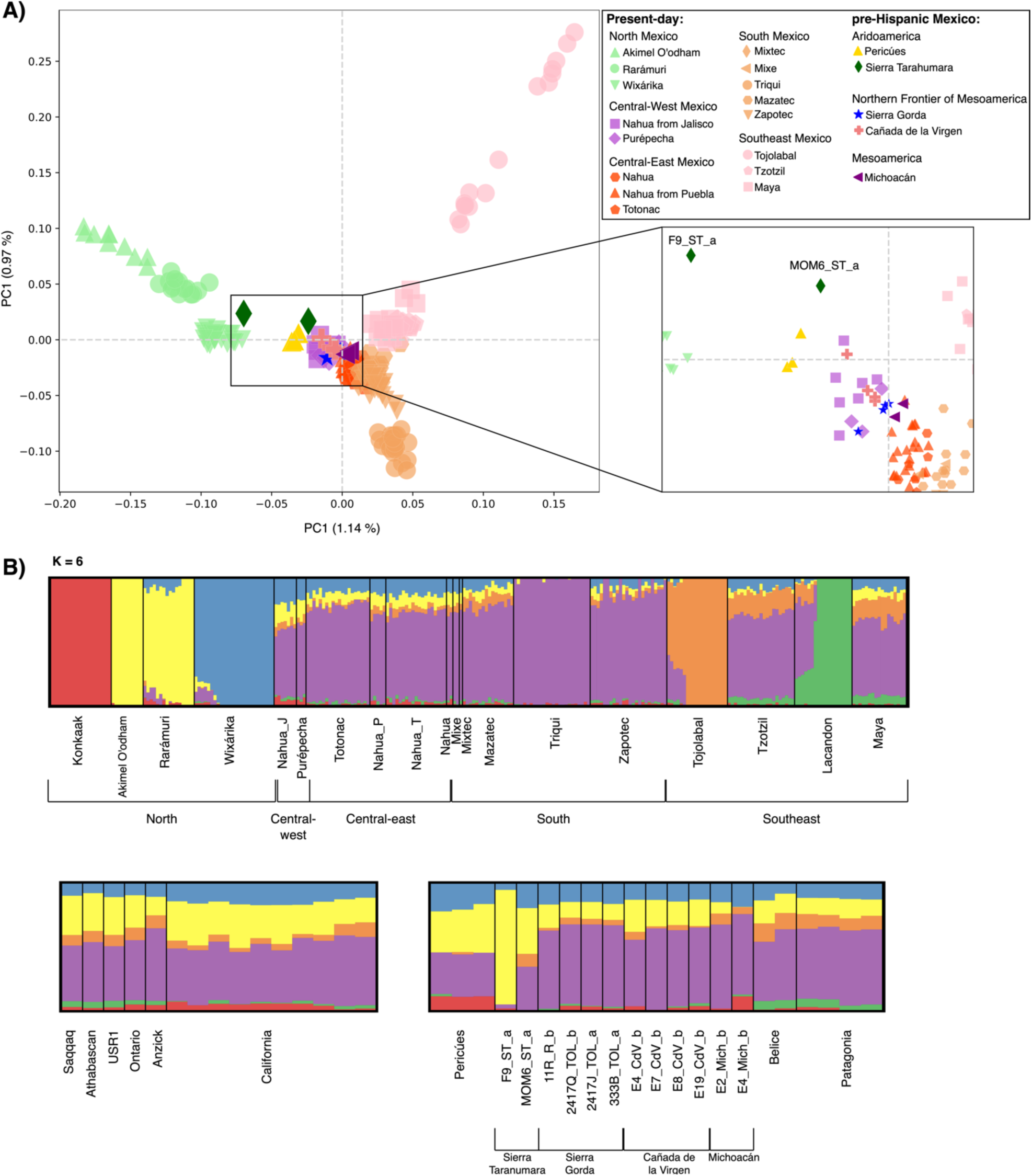
PCA and Admixture plots. a) Principal Component Analysis of the reference panel of present-day Indigenous Populations (see methods), the genomic data of ancient individuals from the Americas previously published and the pre-Hispanic individuals from Mexico were projected onto the panel with lsqproject from the eigensoft package version 6.0.1.1. The northern individual F9_ST_a clusters near current northern populations and the individual MOM6_ST_a clusters with central-west populations as the pre-Hispanic individuals from central Mexico. Pre-Hispanic individuals SG and CdV cluster close to present-day central-west indigenous populations, and Michoacán individuals cluster closely to central-east indigenous populations. b) Unsupervised clustering analysis using ADMIXTURE for k=6, showing component red, yellow and blue in northern populations, component green and orange in southeast populations and component purple in south and central Mexico populations. Individuals from the Sierra Tarahumara and Pericúes, from northern Mexico have a bigger proportion of the yellow component shared with present-day Akimel O’odham and Rarámuri from northern Mexico. Prehispanic individuals from central Mexico have similar ancestral components as present-day central Mexico populations. Prehispanic individuals from the Sierra Gorda show similar genetic composition between them.

We contrasted the placement of individuals in PC space with that obtained when applying a Temporal Factor Analysis (TFA) (*56*) to correct the ancestral relationships using the dates of the pre-Hispanic individuals. For this, we included only four pre-Hispanic individuals with >1x genome-wide coverage (F9_ST_a, 2417Q_TOL_b, 2417J_TOL_a, and 333B_TOL_a). This analysis confirmed the closer genetic affinity of individual F9_ST_b with populations from northern Mexico. Toluquilla (SG) individuals slightly shifted towards the central-east populations in contrast with their position in the PCA plot that places them between central-west and central-east populations. Furthermore, despite the different periods of the individuals from Toluquilla, they keep clustering together as in the PCA (Fig.S9). To further support this structure, we applied a novel approach named “missing data” PCA (mdPCA, see methods) on the 15 pre-Hispanic individuals from Mexico. We observed a similar clustering of the ancient individuals as in the standard PC projection, with the subtle difference that F9_ST_a clustered within present-day Rarámuri and Wixárika populations, instead of in the vicinity as with the other two methods (Fig. S10).

In agreement, ADMIXTURE analysis (K=6) reveals that pre-Hispanic individuals from northern Mexico (Sierra Tarahumara and Pericúes) have a bigger proportion of ancestry shared with present-day northern populations Akimel O’odham and Rarámuri (yellow component in Fig. 3B). Pre-Hispanic individuals from central Mexico also display a similar genetic composition to present-day central-west and central-east Mexico populations (Nahua from Jalisco, Purépecha, Totonac and, Nahua from Puebla). Interestingly, individuals from SG show a homogeneous genetic composition independently of the period in which they lived. The genetic composition of individuals from CdV is similar to that in SG, but CdV individuals have a slightly higher proportion of the yellow component shared with northern populations. In addition, we found the two individuals from Michoacán have a higher proportion of a different genetic component shared with northern populations (red component observed in Konkaak), E2_Mich_b presents the yellow component while E4_Mich_b presents the red one.

We further explored the structure by computing outgroup-f3 and D-statistics. Outgroup-f3 statistics were of the form f3(Test, Source1; YRI), where the Test refers to the pre-Hispanic individual under analysis, and Source1 being a present-day Indigenous population. D-statistics were of the form D(Pop1, Pop2; Test, YRI), where Test refers to the pre-Hispanic individual under analysis, while Pop1 and Pop2 were all possible combinations of present-day Indigenous populations. For both analysis we considered only the combinations that have >1,000 overlapping Single Nucleotide Variants (SNVs).

We found F9_ST_a to share higher genetic drift with present-day northern Mexico populations: Rarámuri, Akimel O’odham and Konkaak (0.291,0.282,0.277), than with any other present-day population (Fig. S11) (Table S9). Consistently, D-statistics showed F9_ST_a to be significantly more related to the present-day Rarámuri than to any other population (Pop1) when tested in the form D(Pop1, Rarámuri; F9_ST_a, YRI) (D= -0.0763 to -0.0210, |Z| = 6.913 to 20.281) (Fig. S12) (Table S10).

In contrast, the genetic ancestries of pre-Hispanic individuals from central Mexico are more complex. They do not seem to have a higher shared genetic drift with any present-day Indigenous population according to the f3 outgroup values, where almost all standard errors overlap. Furthermore, in D-statistics analysis, we find mostly D=0 in all possible combinations of Pop1 and Pop2 (Fig. S13 – S18) (Table S9-S10). These results are consistent with populations from central Mexico having extensive gene flow between them and not completely diverged from each other.

### Genetic diversity in pre-Hispanic Mexico

The ancient genomic data we generated also enabled us to make inferences about past genetic diversity and effective population sizes at the sites of study. This is relevant because it allows us to get insights into the differences between the pre-Hispanic populations, contrast past and present-day genetic diversity, and investigate changes in patterns of genetic variation that may have arisen due to the drought period. To this end, we estimated the conditional nucleotide diversity (CND) (*57*) and runs of homozygosity (ROH) (*58*), and contrasted these between the different pre-Hispanic individuals and those observed in present-day Indigenous populations from Mexico (*34, 50*). CND, a relative measure of genetic diversity, was estimated in pairs of individuals at genomic positions that are variable in African individuals (*57*) and grouping individuals per archeological site. This analysis included 16 pre-Hispanic individuals (Pericúes, Sierra Tarahuamara, SG, CdV and Patagonia) with at least 10,000 SNVs in common with the other individuals from the same archaeological site to perform the analysis by pair of individuals (Table S11 and S12). We found similar values of CND in pairs of pre-Hispanic individuals to those observed in present-day Indigenous populations from Mexico, except for Pericúes from Baja California, who had the lowest CND values (Fig. S19) (Table S12). This is in line with Pericúes being hunter gatherers, likely with small population sizes and living in isolation in Baja California (*59*).

On the other hand, the ROHs reflect the proportion of the genome in a homozygous state, which is inversely proportional to the heterozygosity, and can be the result of demographic scenarios that lead to low population sizes and endogamy, thus it can be used to infer such processes (*60–64*). We performed this analysis for six pre-Hispanic individuals with the highest coverage; namely two from northern Mexico (the Pericú B03 and F9_ST_a from Sierra Tarahumara) and four from central Mexico (three from Toluquilla and one from CdV) (Table S13). Of the three Toluquilla individuals included in this analysis, one was dated pre-drought (2417Q_TOL_b), and two post-drought (2417J_TOL_a and 333B_TOL_a). Pre-Hispanic individuals display fewer segments in ROH than some present-day Indigenous individuals (especially present-day individuals from northern Mexico). The segment size distribution of ROHs suggests that the pre-Hispanic individuals studied here belonged to populations with small effective population (Ne) sizes (2Ne= 1,600-6,400) (Fig. S20), which is in agreement with Ne previously calculated for present-day northern (Rarámuri, Ne= 2,419; Wixárika, Ne= 2,193) and southern (Triqui, Ne= 2,407; Maya, Ne= 2,750) populations (*38*). We contrasted the CND values and ROH estimations of Pericúes from Mexico with another isolated pre-Hispanic population from South America, Patagonia (1,275 – 895 BP) (*24*), expecting low levels of genetic diversity in the latter too. We found that two pairs of Pericúes had similar CND levels as Patagonia individuals (∼0.16-0.175), and the third pair of Pericúes presented a lower CND value (∼0.14). However, in ROH analysis, we found the four individuals from Patagonia have a higher sum of fragments in ROH than the B03 Pericú individual, suggesting that the Pericú population had a bigger population size than that of South American Patagonian individuals (Fig. S20). In summary, the CND and ROH information showed that pre-Hispanic and present-day individuals from northern Mexico (Pericúes and Akimel O’odham, respectively) present the lowest CND values and the highest sum of inferred ROH >4 cM compared to other populations of their respective period. Individuals from the Sierra Tarahumara, Toluquilla and Cañada de la Virgen show similar values of genetic diversity and sum of inferred ROH segments. Furthermore, in the case of Toluquilla, where we have three individuals from different dates, we found that the CND value increase with the date difference between pairs of individuals, probably reflective of the accumulation of new mutations or genetic drift through time during the 489-680 years of difference between 2417Q_TOL_b and the individuals of a later period (2417J_TOL_a and 333B_TOL_a).

### Genetic continuity before and after the 900 – 1,300 CE droughts in the Sierra Gorda

To formally test the hypothesis of population replacement in the SG from the NFM during the 900 – 1,300 CE droughts (*7*), we applied different combinations of outgoup-f3 and D-statistics using individuals with more than 1000 SNVs overlapped. For the former, we used the individuals with the highest depth from SG as the test populations in outgroup-f3 of the form f3(Test, Source1; YRI), with individuals either representing before (TOL_b and R_b) or after (TOL_a) the climate change episode. Source1 represents another pre-Hispanic individual, while YRI represents the outgroup population. This allowed us to test if individuals from the SG shared higher genetic drift between them than with any other pre-Hispanic individual from other archaeological sites, and if this difference has a temporal significance.

We observed that the genetic affinities of the pre-Hispanic individuals from the SG before and after the climate change episode share higher genetic drift between them than with any other pre-Hispanic individual (Fig. 4A), except for the individual 11R_R_b, who shows higher genetic drift with the individual E8_CdV_b from CdV (Fig. S21) (Table S14). However, standard errors for 11R_R_b are high, probably due to a low coverage genome-wide for this individual (Fig. S21). This is in contrast with the outgroup-f3 values obtained when pre-drought CdV individuals were used as the Test population f3(CdV, Source1; YRI) (Fig. S22). Here, the highest outgroup-f3 value systematically corresponds to another CdV individual but corresponds to the second-grade relative according to the ‘READ’ analysis (see methods). The following closest individuals do not belong to CdV. In the case of the two individuals from Sierra Tarahumara, the higher value of genetic drift is between both (Fig. S23).

**Fig.4.**
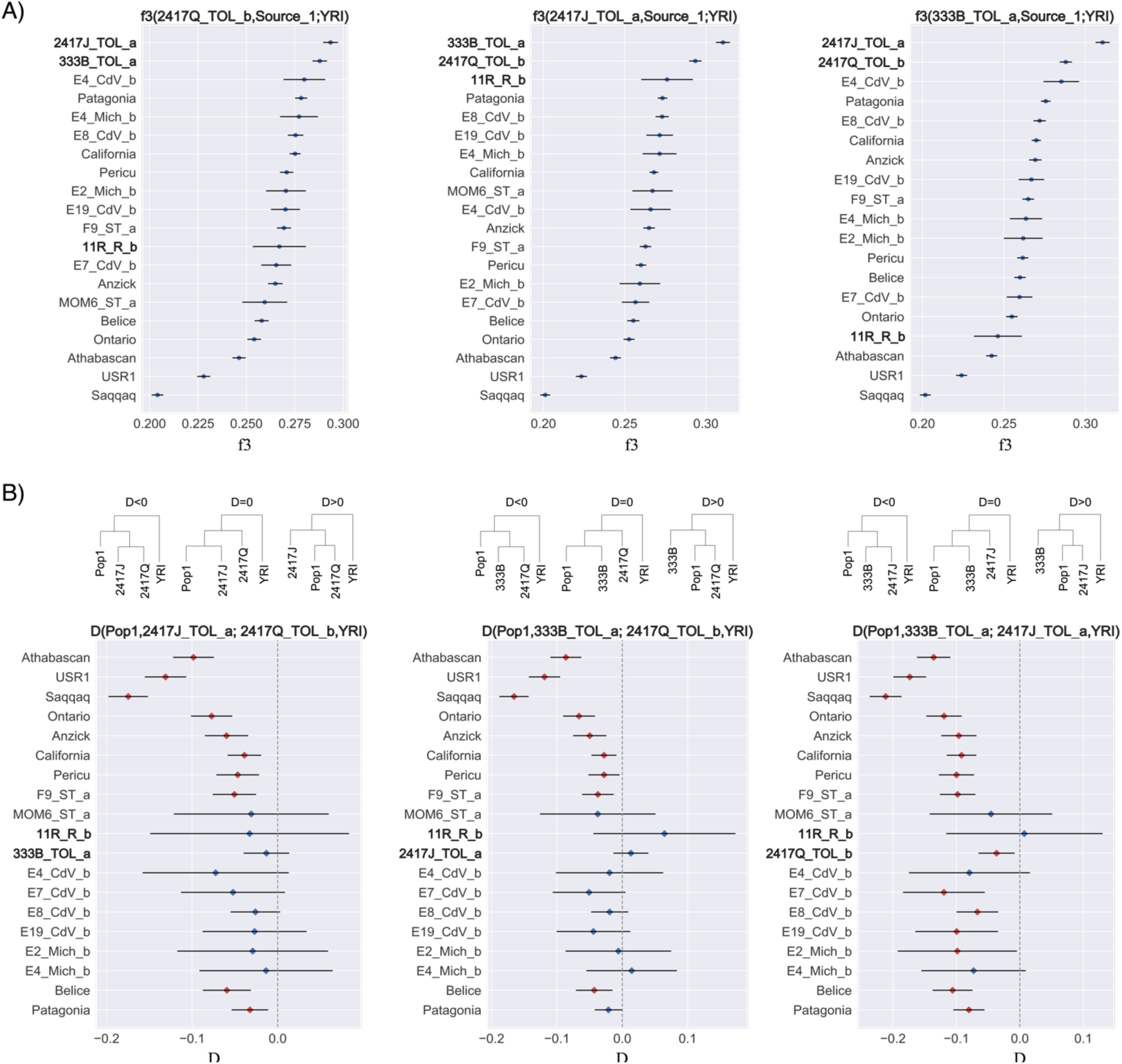
Population continuity in Toluquilla. A) Outgroup f3 statistics for pre-Hispanic individuals from the Sierra Gorda, 2417Q_TOL_b, 2417J_TOL_a, and 333B_TOL_a compared with pre-Hispanic individuals from Mexico and the Americas. Higher values of f3 indicate higher shared genetic drift. Point estimates and one standard error are shown. Individuals 2417Q_TOL_b, 2417J_TOL_a, and 333B_TOL_a share higher genetic drift between them than with any other ancient Native American. B) D statistics analysis in the form D(Pop1,Toluquilla1;Toluquilla2;YRI). Pop1 is any of the other pre-Hispanic individuals from Mexico or America and shown in the Y axis. YRI corresponds to individuals from Africa from the 1000 Genomes Project (*55, 109*). Expected tree topologies according to different D values are drawn on the top of the plot. Individual IDs in the trees are indicated with no suffixes. Red dots indicate significant deviations from D=0 (|Z|>3). When Toluquilla1 is the pre longstanding drought individual 2417Q_TOL_b, and Toluquilla2 is one of the post drought individuals (2417J_TOL_a or 333B_TOL_a), all Toluquilla individuals are equally related to the rest of pre-Hispanic individuals from central Mexico (D=0), but show a higher genetic relation when Pop1 is a population from northern Mexico. When Toluquilla1 and Toluquilla2 are both post drought individual, they tend to be more related to each other than to any other pre-Hispanic individual (D<0). Individuals from the same archaeological site (Sierra Gorda) are highlighted in bold.

D-statistics were calculated in the form (Pop1, Pop2; Test, YRI), where Test refers to the pre-Hispanic individual under analysis, Pop1 representing any pre-Hispanic individual, and Pop2 representing a pre-Hispanic individual from the same region (Sierra Tarahumara, SG or CdV) as Test (Fig. S24 – S26) (Table S15).

Individuals from SG are generally more closely related between them (D<0) in D tests of the form D(Pop1,SG1; SG2, YRI) (Fig. 4B), except when analyzing 11R_R_b (Fig. S25). We do not find a higher relationship between the individual 11R_R_b from Ranas and the other individuals from the Sierra Gorda (Toluquilla). We caution that error bars in D(Pop1, SG; 11R_R_b, YRI) are wide (Fig. S25). In most of the cases, D statistics in form D(Pop1, SG; 2417Q_TOL_b, YRI) show a significantly higher relationship (|Z|> 3) between individuals from Toluquilla than with any other North America or South America individual. Indeed, the two individuals who lived after the megadrought (2417J_TOL_a and 333B_TOL_a) are significantly more closely related than to any other pre-Hispanic individual, except when compared with the individuals MOM6_ST_a (from the north), 11R_R_b (from SG), E4_CdV_b (from CdV) and E4_Mich_b (from Michoacán) (Fig. 4B), for which the D-statistic is not significant probably due to the low-coverage in these genomes which increases the standard error. In a scenario of a population replacement, we would expect the Toluquillan individuals who lived after the drought to be more closely related to pre-Hispanic individuals with higher proportions of northern ancestry (Pericúes and F9_ST_a) than to Toluquillan individuals who lived before the drought (2417Q_TOL_b), since the alleged population replacement occurred from the north. Therefore, our results suggest genetic continuity in Toluquilla before and after the reported climate change event. In some cases, D-statistic values were not significant when another pre-Hispanic central Mexico individual was used as Pop1. These results could be related to the low coverage mentioned for some pre-Hispanic individuals, or they might speak about the previously reported heterogeneity in central Mexico (*38, 39*).

We then used f-statistic-based admixture graphs (qpGraph) (*65*) to characterize the ancestry proportions of the pre-Hispanic individuals of SG (Fig. 5). For the model, we used the 11,500-kyo USR1 (*21*) as the surrogate for ancient Beringian, 10,700-kyo Anzick (*22*) individual as the surrogate for ancestral Southern Native American ‘SNA’(*14*) and the 4,200-kyo Ancient Southwestern Ontario or ASO individual (*15*) as the surrogate for Northern Native American ‘NNA’(*14*). In the analysis, we included the pre-Hispanic individuals 2417Q_TOL_b, 2417J_TOL_a, and 333B_TOL_a and present-day Indigenous populations in Mexico (*54*) representing the northern (Konkaak), central (Purépecha or Nahua from Puebla), and southern (Triqui) regions. The results show that the oldest individual 2417Q_TOL_b seems to have a higher contribution of the ancestral B branch than the individuals after the megadrought (2417J_TOL_a and 333B_TOL_a) (Fig. 5). However, overall results (PCA, TFA, mdPCA, Admixture, f3-outgroup and D-statistics) point to a genetic continuity for the Toluquillan individuals.

**Fig.5.**
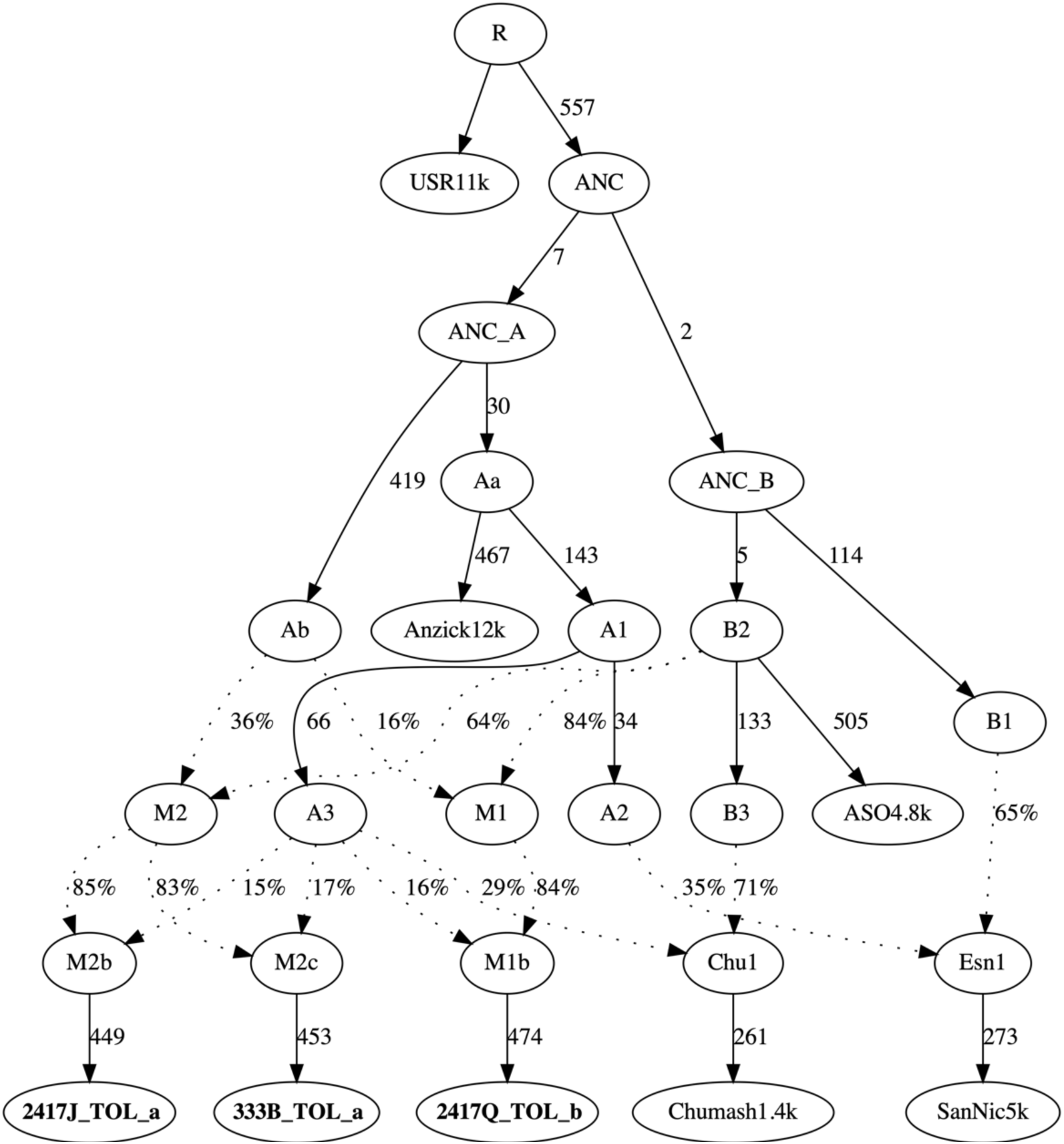
Ancestry proportions for the individuals from Toluquilla (Sierra Gorda) before and after the megadrought analyzed through qpGraph. All individuals show a similar percentage of ancestry from A3. The individual 2417Q_TOL_b seems to have a higher contribution of the ancestral B branch from the ancestor M1, than the individuals after the megadrought who had the ancestor M2. All f statistics are within 1.75 standard error. The tree has no inner 0 branches and obtained a |Z-score| of 1.757.

### Admixture model for Central Mexico populations

Given the complex relationships observed for ancient individuals in central Mexico, we tested additional demographic models using qpGraph. The base model that fitted the pre-Hispanic populations includes Anzick (*22*) individual as the ancestral SNA (12), present-day Athabascan (*14*) as the representative for NNA (*14*), and present-day Indigenous populations in Mexico (44) representing the northern (Konkaak), central (Nahua from Puebla), and southern (Triqui) regions. Furthermore, this model involves a split of the Southern Native American into S2A and S2B sources. The pre-Hispanic populations from SG and CdV show different levels of ancestry from Southern Native American and Northern Native American. We found that the population from SG had a higher percentage of the Southern Native American branch from S2B (64%) than the population from CdV (37%) who had 62% ancestry shared with the Northern Native American branch, indicating a different demographic history for each of these populations despite its relative geographic proximity (Fig. S27). This model supports previous studies that report several admixture events between the two branches that have given rise to Central and South American populations (*15*). For the Michoacán population, we obtained terminal branches with no support (zeros) and, because of that, we did not make conclusions with this model for the Michoacán pre-Hispanic population (Fig. S27).

### Genetic continuity from 1,100 to the present in Sierra Tarahumara, northern Mexico

We decided to evaluate the possible scenario of population continuity in the Sierra Tarahumara from northern Mexico, considering that the individual F9_ST_a was significantly more related to present-day Rarámuri than to any other indigenous population from Mexico in f3 and D statistics. Thus, we used the population continuity test described in (*66*), in such a way that the null hypothesis of population continuity is tested using read counts in the ancient individual and allele frequencies in the reference population (Rarámuri). Using this method, we found no population continuity between F9_ST_a and present-day Rarámuri and rejected the null hypothesis with a p-value of 10^-499.3.

### Ghost population contribution to pre-Hispanic Mexico

Earlier reports have demonstrated the detection of ancestry from an unsampled group, designated UpopA, among the present-day Mixe from Mexico and Lagoa Santa from Brazil populations (*67*). We tested the presence of UpopA in the pre-Hispanic individuals, using an admixture graph model with the same topological framework but some modifications compared to (*67*). We used Mbuti as an outgroup, Ami as the East Asian, MA1 as the ancient north Eurasian, USR1 as the ancient Beringian, Anzick-1 (*22*) and Spirit Cave (*67*) as the NNA, and Athabascan as the SNA. Interestingly, the Mummy F9_ST_a from the north and the individuals from CdV from central Mexico have ancestry from a ‘ghost’ population, at 28% and 17%, respectively (|Z-score|: 2.142 and 1.655, respectively) (Fig. 6), consistent with the “ghost” UpopA ancestry reported in Mixe. We repeated this test with the Mixe population to verify the contribution of the ghost population in our tested admixture graph structures, and found Mixe to have an ancestry coming from an unsampled UpopA population at 13% (|Z-score|: 2.34) (Fig. S28), which is similar to previous estimates of 11% (*67*). Models including individuals from other archaeological sites had |Z-score| values over 3 and branches with inner zeros, thus we rejected them.

**Fig.6.**
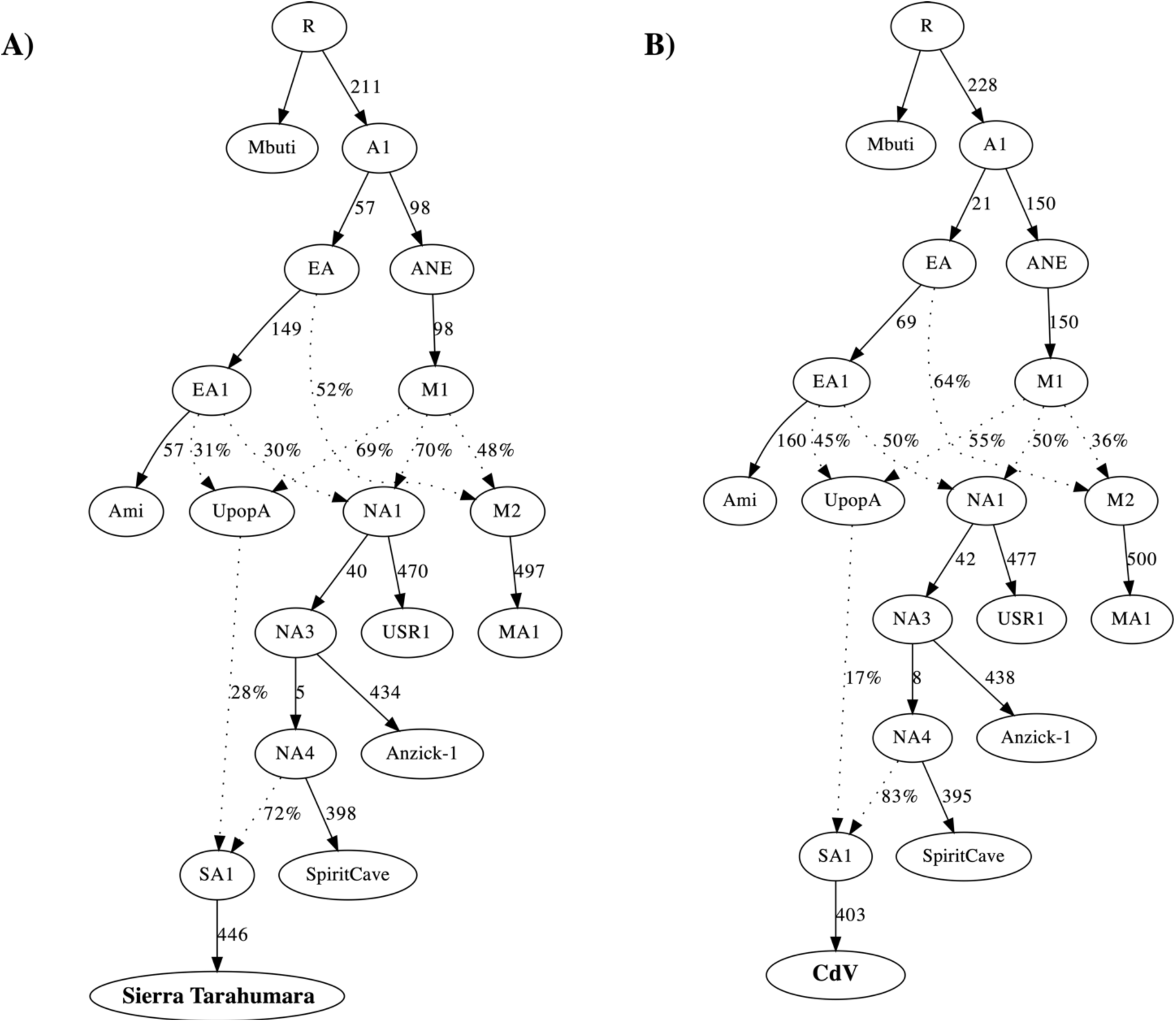
Ghost population in pre-Hispanic individuals from Mexico. A) qpGraph model for F9_ST_b. The tree has no inner 0 branches and obtained a |Z-score| of 2.142. B) qpGraph model for individuals from Cañada de la Virgen. The tree has no inner 0 branches and obtained a |Z-score| of 1.655. Models show different proportions of the ghost population UpopA in the pre-Hispanic individuals from Mexico, being 28% of UpopA for the Sierra Tarahumara individual and 17% of UpopA for CdV individuals. All f statistics are within 1.75 standard error.

## Discussion

We have generated the largest available ancient genome data set from Mexico to address long-standing questions about population dynamics between Aridoamerica and Mesoamerica. The data allowed us to i) describe levels of population structure and genetic diversity in Mexico prior to European colonization, ii) test a previous hypothesis of population replacement in central Mexico following a drastic climate change 1,100-700 years ago, iii) demonstrate a complex admixture model for central Mexico populations, and iv) detect a contribution from unsampled population A (UpopA) to some northern and central Mexico populations.

We find that present-day Indigenous populations from Mexico exhibit substantial geographical structure following a northwest-southeast cline (*54*). The pre-Hispanic individuals from this study also follow this structure. They cluster in proximity to present-day Indigenous populations from the same geographical area based on PCA, TFA, and mdPCA, and exhibit similar ancestral components based on ADMIXTURE analysis, except for the individual MOM6_ST_a (Fig. 3 and S9-S10). This reflects an overall conservation of the genetic structure of the populations inhabiting the Mexican territory (*54*) for at least 1,400 years (considering that the most ancient individual used for autosomal genomic analysis is 1,400 years old), which is consistent with demographic models inferred from exome (*38*) and genome-wide genotype (*68*) data from present-day Indigenous populations from Mexico. These models propose a northern/southern population split between 4,000 and 10,000 years ago followed by multiple waves of admixture events between them (*68*). The exome data further supports subsequent splits within each of these regions 6,523 and 5,715 years ago for north and south, respectively (*38*).

This geographical structure is also reflected in the maternal lineages. Even though we identified previously unreported variants in the ancient mitogenomes (Table S5), the spatial distribution of the haplogroups found, namely A, B, C and D, closely resembles that of present-day Mexico (*27, 36*). The information on the paternal lineage was limited, since, despite confirming the presence of the Native American chromosomal haplogroup Q in the pre-Hispanic males, there was no clear differential spatial distribution of the two identified sub-haplogroups, Q1a2a1-L54 and Q1a2a1a1a1-M3, in both Aridoamerica and Mesoamerica. Few studies have recovered information of ancient Y chromosome haplogroups across the Americas and, given the sex-biased admixture events after European colonization, the frequency of haplogroup Q in the present-day population is low. Thus, further studies on more ancient individuals from the Americas at higher coverage will likely shed light on the demography of the male lineages.

We delved further into the past demography of the Indigenous population by performing a Bayesian coalescent analysis on the mitogenome data. The resulting Extended Bayesian skyline plot (EBSP) showed a larger female population size and a less drastic population decline upon European colonization in Mexico compared to other regions in the Americas (based on the 95% CI) (*41, 69*). This is intriguing considering previous reports of massive population decline during European conquest (*70, 71*). However, these estimates were based primarily on historical accounts of the early colonial Mexican population through taxation lists of most settlements across Mexico but not including all of them (*71*). The mitogenomes used for the EBSP belong to individuals across the Mexican territory ranging from the north to central Mexico. Thus, we suggest that the demographic impact of European colonization on the female population was variable across Mexico, which is generally less drastic than in insular and isolated regions of the Americas. EBSP, including only modern mitochondrial genomes, shows the female population bottleneck experienced after European contact (Fig. 2B). In contrast the EBSP that includes only pre-Hispanic mitochondrial genomes does not show the female population decline (Fig. S6A).

The characterization of past genetic diversity revealed that ancient Aridoamerica (Sierra Tarahumara) and NFM populations (Toluquilla and CdV) have similar levels of conditional nucleotide diversity (CND) between them and to those observed in some present-day Indigenous populations from northern (Akimel O’odham) and southern Mexico (Mixe and some Mayan) (Fig. S19). However, we caution that we lack genome-wide genetic information of present-day individuals from the same site as the pre-Hispanic individuals analyzed here to make a direct temporal comparison. We observed, however, that the lowest CND and the highest total number of segments in ROH were found in the ancient hunter gatherer Pericúes, and the present-day Akimel O’odham population, both from northern Mexico. In contrast, pre-Hispanic individuals from the NFM and present-day populations from central, southern, and southeastern Mexico have the lowest total number of segments in ROH (Fig. S20). Altogether, these results are reflective of the lower population sizes maintained in isolated Aridoamerica populations and larger population sizes in pre-Hispanic and present-day populations in Mesoamerican (central to southeastern Mexico). Nonetheless, it is important to note that the pre-Hispanic individuals from this study come from small-sized villages (*72, 73*) and do not necessarily reflect the demography of other pre-Hispanic populations from larger and multiethnic pre-Hispanic metropoles (e.g., Teotihuacán, Tenochtitlán, or Palenque) (*74, 75*) where higher levels of diversity and lower ROH would be expected. Ancient DNA studies on individuals from these sites could reveal more accurately the extent of genetic diversity in these megalopoleis prior to the population collapse resulting from European colonization.

To directly test the hypothesis of a population replacement of Mesoamerican populations at the NFM by Aridoamerican hunter gatherers because of a drastic climate change in the years 900-1,300 CE, we studied individuals from the Sierra Gorda (Toluquilla and Ranas sites), for which we had access to remains from the time before (671-867 CE) and after (1,160 ± 60 and 1,351± 50 CE) the resulting megadrought. Using D statistics, we observed that the pre-drought individual 2417Q_TOL_b and the post-drought individuals 2417J_TOL_a and 333B_TOL_a, all have the same genetic relationships to present-day Indigenous populations. In other words, 2417J_TOL_a and 333B_TOL_a are not significantly more related to northern (Aridoamerica) populations than 2417Q_TOL_b, as it would be expected under a scenario of population replacement in the NFM. This population continuity is further supported by the longstanding presence of the mitochondrial lineage A2d in the region for at least 1,480 years (Fig. 2A). Notably, we found shorter ROH segments in the pre-drought pre-Hispanic individual 2417Q_TOL_b than in the post-drought individuals 2417J_TOL_a and 333B_TOL_a, which might indicate a reduction in the population size post-climate change period. Altogether, these results suggest population continuity in the Sierra Gorda of the NFM, before and after the climate change episode. Additional assessment of this hypothesis at other NFM sites will shed light into the population migrations in other regions of the NFM.

The observed population continuity in the Sierra Gorda might have been supported by the favorable climatic conditions of the northern Sierra Gorda Mountain range, where both Toluquilla and Ranas sites are located. The landscape could have helped maintain higher humidity conditions than in other regions of the NFM. In this part of the Sierra, the humid winds from the Gulf of Mexico course trough. The area is characterized by coniferous and mixed forests, where the average maximum temperature is 25°C, and the minimum temperature is 6-7°C (*76*). This landscape is very different from the conditions found in Cañada de la Virgen, Guanajuato, where the climate is considerably more arid. CdV is located north of the Lerma River, where rainfall is known to be scarce. The inhabitants of that region had to develop systems for the capture and maintenance of water needed for cultivation, which probably became impossible after the prolonged droughts. This forced them to abandon the site during 1,000-1,100 CE and migrate to areas with better climatic conditions suitable for agriculture, which was the core of their primary subsistence activity (*73*). In contrast, the main subsistence strategy in Toluquilla and Ranas was mining. Residents at these sites exploited cinnabar for trade purposes with other villages and metropolises (*72, 77*). Cinnabar was of great sacred value to pre-Hispanic cultures. Some ancient communities used it to anoint their dead, especially in cases of highly regarded individuals, and as part of ritual offerings (*77, 78*). We hypothesize that the cinnabar trade and the landscape of the Sierra Gorda allowed the peoples of Toluquilla and Ranas to subsist despite low rainfall conditions during the drought.

On the other hand, we were also interested in exploring putative scenarios of population continuity between pre-Hispanic and present-day populations. The only pre-Hispanic individual found to be significantly more related to a present-day population was the individual F9_ST_a, who was significantly more related to present-day Rarámuri than to any other present-day population from Mexico. This relationship was expected since F9_ST_a was found in the Sierra Tarahumara, where present-day Rarámuri currently reside. The northern region of Mexico was one of the most difficult to access and the last areas to be invaded by the Europeans in the seventeenth century, due to its geography and the resistance movements by the local people. Nowadays, Rarámuri still preserve their pre-Hispanic cultural elements in language, music, dresses, and handcraft (*79*). However, the genetic continuity between F9_ST_a and the Rarámuri was rejected despite the high genetic affinities between them. We hypothesize that this genetic population discontinuity observed from this analysis can result from the changes in allele frequencies derived from the admixture events, population bottleneck, and isolation experienced by Rarámuri since pre-Hispanic times and after the European colonization (*68*).

We were also interested in obtaining insights into the demography of central Mexico populations since previous attempts of demographic modeling have been hampered by the high genetic heterogeneity observed in these populations (32, 33). Our outgroup-f3 and D statistics results show that, although pre-Hispanic individuals from central Mexico share higher genetic drift with present-day populations from that region, none of these relationships were statistically significant. We thus leveraged the ancient genome data to propose an admixture model for central Mexico populations. Using qpGraph we found that the pre-Hispanic populations from central Mexico, all have different ancestry sharing with the NNA and SNA branches. This was expected given previous studies on ancient genomes from Central and South America reporting multiple admixture events between these two branches after their split ca. 15,000 years ago (*15, 68*). Together, these observations point to a scenario in which populations from central Mexico have not completely diverged from one another, possibly due to extensive gene flow as expected based on the active commercial exchange between different Mesoamerican populations for centuries (*80*). This interaction was mainly through trade routes and alliances between different nations (55), as has been evidenced in present-day Indigenous populations who share identity-by-descent (IBD) segments (43, 54) and through the analysis of STR loci (56). In addition, there was also gene flow between Mesoamerican and Aridoamerican populations in pre-Hispanic times, as evidenced by the study of IBD segments in present-day Indigenous populations from Mexico (*68*). However, this gene flow occurred less frequently than between Mesoamerican populations. The individual MOM6_ST_a from northern Mexico may have been the result of such admixture between Mesoamerican and Aridoamerican ancestors. This individual has a higher shared genetic drift with F9_ST_a (also from Aridoamerica), as evidenced by outgroup-f3 tests. However, PCA and ADMIXTURE analyses show that MOM6_ST_a has ancestry components found in central west present-day populations. Also, using D statistics, we found MOM6_ST_a to be equally related to all individuals from Aridoamerica and Mesoamerica, supporting the idea of gene flow between these two areas.

Furthermore, the qpGraph admixture models explored for the pre-Hispanic populations showed F9_ST_a and the ancient individuals from CdV have ancestry from an unsampled population that we hypothesize is the ghost population previously found named UpopA (*67*). The contribution from UpopA was identified in present-day Mixe from Mexico and Lagoa Santa, Brazil, through qpGraph models and estimated to have diverged ∼24,700 years ago from Native Americans (*67*). Also, a more recent study through D statistics analysis has suggested that UpopA contributed to present-day populations from north Mexico as Akimel O’odham, Guarijo, Rarámuri, Cora, Mexicanero, and Tepehuano, as well as in present-day populations from central and south Mexico as Mixe, Totonac, Nahuas from Puebla, Otomi from Hidalgo, Chocholteco and, Mocho (*68*). Our study is the first to identify this UpopA ancestry in admixture models involving pre-Hispanic individuals from Mexico. Further aDNA studies from the Americas could help better characterize the source of this UpopA ghost population contributing to many present-day Indigenous populations from Mexico.

In summary, we find that the pre-Hispanic population structure from over a thousand years ago, can still be observed today. Our work, together with previous studies (*15, 38, 39, 54, 68*), show that the demographic events that gave rise to Aridoamerican and Mesoamerican populations are more complex than previously thought. Commercial trade routes may have contributed to increased mobility, facilitating gene flow between different populations within and between various cultural areas. Furthermore, we found genetic continuity in the Sierra Gorda region at the NFM which suggest the local population stayed in their homeland despite the longstanding droughts that forced other NFM populations to abandon their cities. This study opens the door for further research to address the questions of the unknown genetic past and population dynamics of Mexican pre-Hispanic populations, whose genetic legacy is retained today among Indigenous and admixed populations.

## Materials and Methods

### Laboratory procedures

DNA extraction and library preparation (before amplification) were performed in the Human Paleogenomics Laboratory, a clean lab facility at the International Laboratory for Human Genome Research from the Universidad Nacional Autónoma de México (LIIGH-UNAM). Bones and teeth surfaces were cleaned with a 1% sodium hypochlorite solution, followed by a solution of 75% ethanol. Then, the surface was UV irradiated (256 nm) for 1.5 minutes on each side using a UVP CL-1000 crosslinker. A Dremel tool was used to remove the outer surfaces of bones and teeth. Bones were cut to get an inner sample of around 100-200 mg for DNA extraction. Teeth were cut at the cementoenamel junction, the roots were wrapped in aluminum foil and pulverized using a hammer. Around 100-200 mg of the root was used for DNA extraction. DNA extraction was performed using the methods described in (*81, 82*), as described in Table S2.

DNA extraction from individual P_CCM_b was performed taking 150 mg of mummified skin. The sample was washed in deionized sterile water and dried. Then, the sample was UV irradiated (256 nm) for 1.5 minutes using a UVP CL-1000 crosslinker. Epithelium was cut in small pieces and incubated at 50°C for 24 hours in lysis buffer (10mM Tris-HCl, 10mM NaCl, 5mM CaCl2, 2.5 mM EDTA, 1% SDS, 10 mg/ml proteinase K, 10 mg/mL DTT). Followed by centrifugation at 16,100 x g for 5 minutes and recovery of the supernatant. Then, DNA purification was performed according to the protocol in (*82*).

Double-stranded libraries for Illumina sequencing were prepared according to (*83*), using single-indexed adapters with 6-bp barcodes or double-indexed with 7-bp barcodes, depending on the sequencing platform, NextSeq550 or NovaSeq, respectively. Libraries were analyzed with qPCR to determine the optimum number of cycles during the indexing PCR step. Barcoded libraries were sequenced for a first screening on the NextSeq550 equipment from Illumina using a 2x75 run at either LANGEBIO’s genomics core facility (National Laboratory of Genomics for Biodiversity, Irapuato, Guanajuato.) or INMEGEN genomics facilities (National Institute of Genomic Medicine, Mexico City). Sequence data were processed as described in the next section. Depending on the quality of the libraries (% endogenous and % clonality), as revealed by the screening sequence data, some were chosen to be subjected to mitochondrial genome capture or whole-genome capture using Daicel Arbor Biosciences (Ann Arbor, MI, USA) commercial kits (Table S2). Captured libraries were sequenced to assess complexity and yield with the tool preseq (*84*) and sequenced to higher depth using the NextSeq 550 (2x75 cycles) at LANGEBIO’s core facility or in the NovaSeq at the Centro Internacional de Mejoramiento de Maíz y Trigo (CIMMYT) with an S1, 2x100 cycles run (Table S2). Sequencing runs not reported previously for Sierra Tarahumara individuals were carried out at the Sci LifeLab Uppsala Sequencing Center and at the Bustamante Lab in the Department of Genetics at Stanford University (Table S2).

### Sequence data processing

Raw reads were processed using Adapterremoval (*85*) for trimming Illumina adapter sequences and collapsing of pairs of reads with an overlap of at least 11 bp (--trimns --trimqualities --qualitybase 33 --minlength 30 --collapse). Collapsed reads with >30bp and quality above 33 were retained for downstream processing.

The retained reads were mapped to the human reference genome b37 (hg19), with the mitochondrial sequence replaced by the revised Cambridge reference sequence (rCRS, NC_012920). Mapping was done with bwa 0.7.13, aln algorithm, seed disabled (-l 500), and keeping reads with mapping quality >25. Clonal duplicates and reads mapping to more than one place in the genome were removed with SAMtools rmdup (*86*) and eliminating reads with labels ‘XT:A:R’ and ‘XA:Z’, respectively. Then, reads were realigned using GATK with RealignerTargetCreator and IndelRealigner (*87*). To authenticate ancient DNA sequence data, mapDamage2 (*18*) was used to assess the damage patterns and distribution of reads length, using default parameters (Fig S1-S4). Alignments to rCRS were separated into new bam files using SAMtools (*86*) to analyze the mitochondrial genome for haplogroup assignment, haplotype network analysis and Extended Bayesian Skyline Plot (EBSP) (see respective sections below for details). For the analysis of autosomal variants, base qualities were rescaled with mapDamage2 (*18*), this step reduces the base quality of the bases in the reads that have a misincorporation from C to T and G to A in the alignment with the reference genome, since this changes might be present due to ancient DNA damage. Depth of coverage was estimated using ATLAS (*88*).

### mtDNA analysis

Reads aligned to the Cambridge reference sequence (rCRS, NC_012920) were analyzed with Schmutzi (*89*) to estimate contamination and obtain a consensus mitochondrial sequence for each individual. Script contDeam.pl from Schmutzi was run with the parameters --library double and the default --uselength, and schmutzi.pl was run with 8 threads. Haplogrep 2 (*90*) was used to assign a mitochondrial haplogroup and quality of assignment for each library using the fasta file with the consensus mitochondrial genome generated by Schmutzi (Table S1).

### mtDNA alignments

Consensus fasta sequences of the whole mtDNA of ancient individuals from this and previous studies were aligned together with previously published modern sequences (*14, 15, 32, 34, 40, 41, 91–100*) (Table S4 and S6) using MEGA X (*101*). Multiple sequence alignments of mitochondrial genomes of the same haplogroup (A, B, C or D) were further visually inspected (i.e. removing common indels, as well as to confirm variants and making sure sequences remain at the same length), to assure an accurate alignment. The same process was also done for a sequence alignment with the combination of all four haplogroups. Common hypervariable sites that are phylogenetically uninformative were excluded from the analysis (309, 315, 515-522 AC indels, 3107, 16182-16183, 16193, and 16519) (*40*). Both sequence alignments for haplotype networks and Extended Bayesian Skyline plots were done following this description, although they contain different samples.

### Haplotype networks

Haplotype median-joining networks were constructed in Popart (*102*) after collapsing aligned full mtDNA sequences into haplotypes with DnaSP . Haplotype networks were constructed with both ancient (from this study and previously published) (n=25) and present-day (n=78) mtDNA of individuals in the Mexican territory (Table S4 and S6). All present-day samples were retrieved from the PhyloTree (version 17) database (*40, 91*). Table S4 reports the ancient individuals for which the mtDNA sequences were used. Only sequences that were reported as being of the same sub haplogroup or neighboring sub-haplogroup as our ancient samples were chosen for sequence comparison.

### Extended Bayesian skyline

Past female effective population sizes were reconstructed using a Bayesian skyline approach (*104*) in BEAST 2 (*105*). The ancient mtDNA sequences, mentioned above were merged with present-day mtDNA sequences collected from published public databases (Table S4 and S6). Alignments were partitioned into five concatenated regions in the following order (Dloop, Coding, rRNA, and tRNA) as proposed by (*41*). HKY+G was found to be the best substitution model for this arrangement of the data using PartitionFinder 2 software (*106*). Substitution rates were estimated following a strict molecular clock model starting from point estimations as in (*107*). Additionally, we used tip calibrations for ancient samples using dating estimates (Supplementary Table S1). We ran Markov Chain Monte Carlo (MCMC) chains of 100 million steps for each haplogroup alignment with a sampling of parameters every 10,000 generations, discarding the first 10 million steps as burn-in. Extended Bayesian skyline analysis was plotted using SkyViz reported in (*42*). Two independent runs of EBSP were run, showing a similar behavior (see Table S7 for statistics of each run). The first run of each EBSP was used to be reported.

### Sex assignment

Determination of biological sex was made using the tool reported in (*45*). This approach computes the proportion of reads mapped to the Y chromosome with respect to the reads mapping to the X chromosome (Ry) and Y chromosomes. According to the method, Ry > 0.075 corresponds to XY, while Ry < 0.016 corresponds to XX (Table S1).

### Y-chromosome haplogroup inference

Y chromosome genotype calling and haplogroup assignment were made as reported in (*108*). Genotype calling was performed sampling one random base at each site of the Y chromosome covered at least once, with ANGSD (-dohaplocall 1 -doCounts 1 -r chrY: -minMinor 0 -maxMis 4). Then, haplogroup assignments were made using the phylogenetic tree of Y chromosome single nucleotide polymorphisms constructed from the 1000 Genomes Project Phase 3, as in (*108*). The most derived haplogroup was assigned.

### Autosomal reference panels

For analysis at a continental level, we constructed a reference panel (Supplementary Table S5) that includes genetic information of: Yoruba (YRI), European Ancestry (CEU), and Chinese (CHB) from the 1000 Genomes Project Phase 3 (*55, 109*), and available genome-wide information for Indigenous populations from Mexico previously published by the Human Genome Diversity Project (HGDP) (*50, 110*) . These genome-wide data were intersected with previously published reference panels of genotype information of Indigenous populations from Mexico (*54*), after masking non-Native American sites as in (*111*) and keeping individuals with > 85% genotype information (mind 0.15) and SNVs with a minor allele frequency of 0.01 (maf 0.01), and with a 90% genotyping rate (geno 0.1) using the software plink (*112*). The final reference panel includes genotype information of 576,409 SNVs from 596 individuals from 4 continental populations (YRI, CEU, CHB, and Indigenous from Mexico) (Table S8).

For the analysis within Mexico, we used the reference panel including only the Indigenous populations from Mexico (without the other three continental populations: YRI, CEU, and CHB) and keeping the individuals with >90% of Native American ancestry. In total, this reference panel included genetic information of 561,327 SNVs from 268 individuals (Table S5).

Both reference panels (continental and within Mexico) where pseudohaplodized as described in the next section.

### Autosomal genotype calling and population genomic analysis

Pseudo-haploid calls of the ancient genomic data were made for the positions included in the reference panel. This was done by randomly sampling one read (when more than one read covered the site) at each site present in the reference panel and keeping the observed base at that site if it had a minimum base quality of 30. If only one read covered the site, the observed base was only kept if it had a base quality above 30. Then, the calls were turned into homozygous genotypes. Similarly, for the genotype data in the reference panel, one allele was randomly sampled at heterozygous sites and turned into a homozygous genotype. For downstream analyses, we kept only ancient individuals with >1,000 SNVs intersected with the panel.

We carried out the analysis of autosomal genomes of 13 pre-Hispanic individuals with the highest depth of coverage per archaeological site: Sierra Tarahumara (n=2, 0.01-1.8x), Sierra Gorda (n=5, 0.005-4.7x), Cañada de la Virgen (n=4, 0.03-0.5x), and Michoacán (n=2, 0.04-0.05x) (Table S1). As a first approach, ancient genomes were compared with present-day Indigenous populations from Mexico and three continental populations (YRI, CEU, and CHB), using a set of 561,3279 SNVs.

#### Relatedness

Relatedness between ancient individuals from the same site was assessed using READ (*113*). A normalization step with a panel of non-related individuals is required previous to estimating the relationship in the ancient individuals, this step was performed with all Nahuas from the reference panel as they are the biggest group in the reference panel. Then, the normalization value was used to run READ with the ancient individuals. Two individuals from Toluquilla (333C_TOL_a and 333B_TOL_a) were identified as first-degree relatives. Thus, the one with the highest coverage (333B_TOL_a) was used for downstream analyses. Furthermore, individuals E4_CdV_b and E8_CdV_b were identified as second-degree relatives, as well as E7_CdV_b and E19_CdV_b. All of these individuals from CdV were kept in the downstream analyses.

#### ADMIXTURE analysis

A first ADMIXTURE (*51*) analysis was carried out on the genotype reference panel including Indigenous individuals from Mexico and the three continental populations (YRI, CEU, and CHB) to identify present-day Indigenous individuals with <90% Native American ancestry. One hundred replicates were run for each k, from k=2 to k=9 with the cross-validation error (-cv) parameter. We found k=4 to be the k with the lowest mean cv error, separating YRI, CEU, CHB, and Native American ancestry. Then the run of k=4 with the best likelihood was chosen and the Indigenous individuals with <90% Native American ancestry were removed from the panel. A second ADMIXTURE analysis was carried out with only the present-day Indigenous populations with >90% Native American ancestry, from k=2 to k=8 using cv error parameter to identify the best value of k, being k=2 the one with the lowest cv error, and separating populations with northern and southern ancestries. Then, a third ADMIXTURE run was carried out including the present-day Indigenous populations and pre-Hispanic individuals. In all cases, one hundred replicates were run with randomly generated seed values; the run with the best likelihood for each k was plotted using pong software (*114*).

#### Principal Components Analysis

We performed principal component analyses (PCA) using smartpca from the software eigensoft v6.0.1 (*52*) and projecting the individuals (lsqproject: YES) on the PCs estimated for the modern populations. The PCA (Fig. 3A) with only the ancient individuals as well as the Indigenous populations from Mexico, only included present-day individuals masked and with >90% Native American ancestry.

#### Temporal Factor Analysis

Temporal Factor Analysis (TFA) (*56*) were run with the reference panel that includes only the Indigenous populations from Mexico. The ancient samples included in the TFA consisted exclusively of individuals from Mexican territory from this study with >1x genome-wide coverage: one individual from Sierra Tarahumara (F9_ST_b), and three from Toluquilla (2417Q_TOL_b, 2417J_TOL_a, and 333B_TOL_a). Genotyping data was manipulated with Plink version 1.90 beta (*112*). The dataset was converted from Plink format to Eigenstrat’s geno format with the command convertf from Eigensoft version 6.0.1.1 (*52*). Missing genotypes were imputed with the snmf function from the LEA package, using the parameters K = 2, entropy = TRUE, and repetitions = 5. The runs with the lowest cross-entropy value were considered for posterior steps. Genotypes were corrected according to the autosomal coverage depth of each sample. We assigned the average depth from the HGDP dataset mentioned in the publication: 35X (*50*) . We assigned the respective coverage read depth for all 11 individuals from SGDP. Modern samples genotyped with microarrays (*54*) did not have a coverage depth value as they represent a different technology without sequencing reads, therefore, we assigned the maximum value from the dataset to all microarray samples, i.e. 56.19X. TFA was applied to these imputed and corrected genotypes with the following parameters: lambda = 5e-1, and K = 2. Imputation, coverage correction, TFA, and the manipulation of the genotyping matrix in geno format were performed with R version 4.0.2. Variance explained by lambda did not change considerably across logarithmic values, thus we chose the default lambda value 5e-1. We plotted factor 1 and 2 as negative when required to match geography and facilitate the comparison between TFA plots.

#### mdPCA

Missing DNA PCA (mdPCA) is a principal component analysis that corrects for genotypes missing due to ancestry masking or degradation in ancient DNA samples. The correction is performed by comparing genetic distances between all samples. The dataset included 15 ancient samples and 231 modern samples. Modern indigenous populations included Northern Mexico (Pima, Rarámuri, and Wixárika), Central-West Mexico (Purépecha, and Nahua from Jalisco), Central-East Mexico (Nahua, and Totonac), South Mexico (Mazatec, Zapotec, and Mixe), and Southeast Mexico (Maya, Tzotzil, and Tojolabal). The ancient samples included in the mdPCA consisted exclusively of individuals from Mexican territory: two individuals from Sierra Tarahumara (La Ventana: F9_ST_a, and MOM6_ST_a), three Pericú individuals (Pericu_B03, Pericu_BC25, and Pericu_BC30), four individuals from Sierra Gorda (Ranas: 11R_R_b; Toluquilla: 333B_TOL_a, 2417J_TOL_a, and 2417Q_TOL_b), four individuals from Cañada de la Virgen (E4_CdV_b, E7_CdV_b, E8_CdV_b, and E19_CdV_b), and two individuals from Michoacán (La Mina: E2_Mich_b, and E4_Mich_b). Genotyping data in plink format was converted to an unphased VCF format with Plink version 1.90 beta (*112*). The method used this VCF file as an input with the following parameters: each individual was plotted as an average of both parental haplotypes (AVERAGE_PARENTS=True), weights for each individual were used inversely proportional to the number of samples from the corresponding population (IS_WEIGHTED=True), the simple weighted of covariance PCA without any optimization was used (METHOD=1), no genotype was masked based on local ancestry calls (IS_MASKED=False), and only the three first PC’s were calculated (NUM_DIMS=3). Individuals with <0.15x whole genome coverage (19,851 SNP markers or less) were further down-weighted to 0.1 to avoid having the principal components be selected based on noise variance stemming from these samples. These included BC30, MOM6_ST_a, 11R_R_b, E4_CdV_b, E7_CdV_b, E19_CdV_b, E2_Mich_b, and E4_Mich_b.

#### Outgroup f3

To estimate each pre-Hispanic individual’s genetic relatedness with a particular pre-Hispanic or present-day Indigenous population, we performed outgroup f3 statistics using ADMIXTOOLS v5.0 (*65*). Outgroup f3 was calculated in the form (Test, Source1; YRI), where the “Test” is the pre-Hispanic individual under analysis, and Source1 being another pre-Hispanic individual or present-day Indigenous population. We used all YRI from the 1000 Genomes Project (*55, 109*) as an outgroup population. Assuming there was no admixture in the tree, the f3 value is proportional to the shared genetic drift between Test and Source1.

#### D statistics

We performed D statistics using ADMIXTOOLS v5.0 (*65*) to identify whether a pre-Hispanic individual had higher genetic affinities with: i) other pre-Hispanic individuals from the same archaeological site than to pre-Hispanic individuals from other sites; or ii) a specific present-day indigenous population than to other present-day population. D values were computed in the form (Pop1, Pop2; Test, YRI). The “Test” refers to the pre-Hispanic individual under analysis, while Pop1 and Pop2 were all possible combinations between pre-Hispanic individuals and present-day Indigenous populations. Under the null hypothesis, Test individual would be equally related to Pop1 and Pop2, and we expect no significant deviations from D=0. Significant D<0 indicates closer relation between Test individual and Pop2. Moreover, a significant D>0 indicates closer relation between Test individual and Pop1. Only absolute values of Z-score > 3 were considered as significant (which corresponds to a p-value of ∼ 0.0027).

#### qpGraph

We utilized the qpGraph tool of the ADMIXTOOLS v5.0 software package (*65*) to construct admixture graphs and test which various models of demographic history best fit the data. With the reference panel including Indigenous populations from Mexico previously published (*50, 54, 110*), we included in the baseline model the following individuals/populations: Mbuti as an African outgroup, USR-1 (*21*) as the reference Beringian, Anzick-1 (*22*) as the reference ancestral Southern Native American, and Athabascan (*14*) or ASO (*15*) as the reference ancestral Northern Native American branch. Present-day Indigenous populations from Mexico and pre-Hispanic individuals were subsequently added to the model in various combinations to test the fit. The following parameters were applied: outpop (NULL), diag (0.0001), hires (YES), blgsize (0.05), and lsqmode (YES). A worst fitting f statistical test of |Z| score >3 is set as a threshold for model rejection.

#### Population continuity test

According to the PCA, TFA, ADMIXTURE, outgroup-f3 and D statistics, individual F9 from north Mexico is significantly closer to present-day Rarámuri. Thus, we analyzed whether individual F9 belonged to the present-day Rarámuri ancestral population using the method published in (*66*). Derived alleles in individual F9 and present-day Rarámuri were identified using ancestral alleles’ information from (ftp://ftp.1000genomes.ebi.ac.uk/vol1/ftp/pilot_data/technical/reference/ancestral_alignments).

Then, allele counts were performed for each site in the present-day Rarámuri (reference population), excluding alleles with frequency 0 or 1. We performed allele counts for the pre-Hispanic individual F9. Parameters alpha and beta necessary for the analysis were estimated using min_samples = 14, which refers to the minimum number of individuals required in the reference panel that have information for each SNV. Under the null hypothesis, it is assumed that reference population and ancient individual belong to a continuous population, rejection of the null hypothesis would mean no population continuity.

#### Conditional nucleotide diversity

We estimated relative levels of genetic diversity for pre-Hispanic and present-day indigenous individuals from Mexico as previously reported in (*50, 110*). The analysis was made according to the approach reported in (*57*). This consists of counting the differences between the ascertained alleles present in one pair of individuals from the same population. For this analysis, we used a panel of 1,938,919 transversions SNVs with a minor allele frequency of 0.01 in the YRI population. In all cases, pseudohaploid genotype calls were performed before the comparisons. CND values were calculated for all pairs of individuals for each present-day Indigenous population and for all pair of pre-Hispanic individuals for each archaeological region. For present-day populations we masked the sites with no Native American ancestry to avoid counting levels of diversity from a different ancestry. Mean and standard error values were estimated 100 times using a window size of 50 Kb even for pairs of ancient individuals. Results are reported in a violin plot, where each dot inside the violin corresponds to the CND value for a unique pair of individuals (Fig. S19).

#### Runs of homozygosity

We used the tool hapROH (*58*) to detect runs of homozygosity in pre-Hispanic individuals. The distribution of ROHs was used as an indirect way of measuring genetic diversity and effective population size. This method detects the ROH’s lengths longer than four centimorgans(cM) and up to 300 cM. The sum of the ROH detected gives insights into the relatedness of the parents of each individual and informs about population sizes. Length of ROH and relatedness between parents are positively correlated. While the higher the number of ROH segments, the smaller the population size and the smaller the number of heterozygote sites, which means less genetic diversity. hapROH uses a built-in set of 1240K SNVs from (*115*) to perform pseudohaploid genotype calls and a reference panel of 5,008 present-day human haplotypes from (*54*). We performed a pseudohaploid variant calling for the 1240K SNVs for each pre-Hispanic genome. hapROH’s manual recommends using samples with a minimum of 400K SNVs intersected with the 1240K SNVs. However, we also included the individual E8_CdV_b, which intersected 369,504 SNVs, and the Pericú individual B03, which intersected 370,026 SNVs, to have at least one individual of these sites in the analyses and considering the number of SNVs is not too far from 400K (Table S13). Detection of runs of homozygosity was made using the set of 5,008 haplotypes mentioned above.

## Supporting information

Supplementary_Table_S1-S3_Sample_infomation

Supplementary_Table_S4-S7_mtDNA_analysis

Supplementary_Table_S8_Reference_panel_present-day

Supplementary_Table_S9_F3outgroup_prehispanic-present-day

Supplementary_Table_S10_Dstats_prehispanic-present-day

Supplementary_Table_S11_CND

Supplementary_Table_S12_CND_pairwise_values

Supplementary_Table_S13_hapROH

Supplementary_Table_S14_F3outgroup_prehispanic-prehispanic

Supplementary_Table_S15_Dstats_prehispanic-prehispanic

## Acknowledgments

**Acknowledgments:** We greatly acknowledge the IT support of Luis Alberto Aguilar Bautista, Alejandro de León Cuevas, Carlos Sair Flores Bautista and Jair Garcia Sotelo of the Laboratorio Nacional de Visualización Científica Avanzada (LAVIS UNAM). We also thank Alejandra Castillo Carbajal and Carina Uribe Díaz from LIIGH-UNAM for wetlab technical support during the project. We thank Alexandra Sockell for technical support at the Bustamante Lab. We thank Diego Ortega-Del Vecchyo and Angélica González Oliver for enriching comments throughout the project on earlier versions of the manuscripts.

## Author contributions

M.C.A.-A. and V.V.-I. conceived and designed the study with input from E.M.P.C. V.V.-I., M.S.-V., M.B.-L., B.M., R.F., K.S., and M.A.N.-C. performed the wetlab work. V.V.-I. processed all sequence data. V.V.-I. carried out most analyses with help from, A.I.-G., M.L., J.M.T., J.E.R.-R., D.A.V.-R., and F.S.-Q. E.M.P.C., A.H.-M., K.S., G.Z., M.V.-M, R.A.-H., A.G.-O., C.V. contributed samples, provided the archaeological context and discussed results. F.A.V., A.M.-E., F.J., A.I., E.H.-S., and F.S.-Q. suggested analyses, discussed results and provided feedback throughout the project. V.V.-I. and M.C.A.-A. wrote and edited the manuscript with input from all authors.

## Data and materials availability

Data will be available from the European Nucleotide Archive under the project accession PRJEB51440, upon publication.

## Ethics statement

See supplementary material.

