## Supplementary material for "Demographic history and genetic structure in pre-Hispanic Central Mexico": Suplementary Materials pdf

**This PDF file includes:**

Supplementary Text
Figs. S1 to S28
References (1 to 22)

**Other Supplementary Materials for this manuscript include the following:**

Supplementary_Table_S1-S3_Sample_infomation.xlsx
Supplementary_Table_S4-S7_mtDNA_analysis.xlsx
Supplementary_Table_S8_Reference_panel_present-day.xlsx
Supplementary_Table_S9_F3outgroup_prehispanic-present-day.xlsx
Supplementary_Table_S10_Dstats_prehispanic-present-day.xlsx
Supplementary_Table_S11_CND.xlsx
Supplementary_Table_S12_CND_pairwise_values.xlsx
Supplementary_Table_S13_hapROH.xlsx
Supplementary_Table_S14_F3outgroup_prehispanic-prehispanic.xlsx
Supplementary_Table_S15_Dstats_prehispanic-prehispanic.xlsx

**Supplementary Text**

Sampling and permits

Processing of samples and data generation for DNA analysis were performed with the approval of the Archaeology Council of the Instituto Nacional de Antropología e Historia (INAH) of Mexico (Permit number 401.1S.3-2022/1152).

Samples include skeletal remains from Northern Mesoamerica: Sierra Gorda - Querétaro (n=13), Cañada de la Virgen - Guanajuato (n=16) and Central Mesoamerica: Michoacán (n=8); along with updated genomic data from Aridoamerica: Sierra Tarahumara – Chihuahua (n=2), previously reported at less resolution (*1*). The most extended period covered here belongs to Sierra Gorda samples from 320 BCE to 1,351 CE (Classic-Postclassic). While the rest of the archaeological sites correspond to Classic (Cañada de la Virgen and Michoacán), Postclassic (Sierra Tarahumara) (see Table S1 for dates).

Ethics statement.

The sampling of the archeological human remains was made upon approval by the Consejo de Arqueología (Archaeology Council) of the Instituto Nacional de Antropología e Historia (National Institute of Anthropology and History) of each collection. The permit numbers are: 401.3S.16-2017/990, 401.3S.16-2019/222

We sampled the minimum possible amount of tissue to avoid unnecessary destruction of these precious materials. For teeth, we tried to separate the root from the crown without damaging the crown. We returned all leftover material to the archeologists responsible of each collection. In addition, we took photographs of each sample during several steps of the sample processing to have a visual archive. Because of historical and socio-political reasons, together with the prominent admixture of different continental ancestries in the territory and the imposed institutionalized narrative of "mestizaje"(*2*, *3*) to minimize the contribution of present-day Indigenous peoples to our national identity (something utterly condemnable), it becomes quite complex to assign a present-day population in Mexico as the single direct descendant of the remains of human ancestors recovered from archaeological contexts. This complexity raises several questions regarding consultation of descendant populations for this kind of research (e.g. should admixed "mestizo" populations who share the geographical area be consulted? who should be considered a representative for them? under what capacity? In part because of these uncertainties, consultation with Indigenous populations for destructive analysis of archeological remains is not a standard procedure in the country. This is on its own an emerging focus of discussion that warrants further investigation and debate beyond the scope of this research (but see Ávila-Arcos et. al 2022) (*4*). However, a series of efforts to communicate genetic findings resulting from this work to local communities from the sites of Toluquilla and Ranas have taken place over the course of the project.

Naming of individuals

Names of individuals are stated according to burial and individual numbers provided by archaeologists. Additionally, we added two labels, one consisting of 1-3 letters that refers to the archaeological site and the other corresponding to one letter that represents the period of the individuals: with a letter 'b' for before the longstanding droughts and 'a' for those from the period after the longstanding droughts. A '_' character separates these three pieces of information. Throughout the text, we refer to all individuals with their complete ID. Those individuals who do not have the last label do not have an estimated date.

### Archaeological context

#### Sierra Tarahumara (Mummy-F9 and Mummy-MOM6)

These individuals and their archaeological context were previously published in (*1*)

#### Sierra Gorda, Querétaro (Mummy-P)

##### Cadereyta cave, Sierra Gorda, Querétaro (P_CCM_b)

In 2002, the discovery of the mummified remains of individual P_CCM_b was reported in the vicinity of the village of Altamira, in the municipality of Cadereyta, Querétaro. The remains were removed and taken to the public prosecutor's offices in the municipal capital. Once the remains were handed over to INAH and archaeologist Elizabeth Mejia Perez Campos, the site was visited and cleaned. Fragments of textiles, organic matter such as bird feathers, and wood formed a bracelet, and human hair braids were found. The piece was taken to the Templo Mayor Museum in Mexico City, where computerized axial tomography, necropsy with an endoscope to see internal organs and samples were taken. Likewise, mtDNA studies were performed by PCR amplification of the HVR at the Centro de Investigación y Estudios Avanzados of the Instituto Politécnico Nacional (CINVESTAV - IPN) in Mexico City and the textile and skin were dated by C13 and C14 at Beta Analytical in Florida, USA. The results showed an infant of one year eight months old who lived in 340 BC, shrouded with a cotton blanket with human hair interwoven and with an offering of maguey quills, stork feathers, pine, oak, and sotol leaves (*5*).

##### Toluquilla, Sierra Gorda, Querétaro

Toluquilla archaeological site is located in the Sierra Gorda, in the State of Querétaro. The occupation of Toluquilla spans ca. 2,000 years, from 400 BCE to 1,550 CE.

Eight individuals, 333A_TOL_a, 333B_TOL_a, 333C_TOL_a, 333O_TOL_b, 333Q_TOL_b, 2417C_TOL_a, 2417Q_TOL_b and, 2417J_TOL_a, from the archaeological site of Toluquilla were sampled for this study. We analyzed samples from individuals from two ritual burials at the site. The burials were analyzed by Dr. Elizabeth Mejia Perez Campos as part of her doctoral thesis (*6*) and were named based on the relevance of the offerings. These burials correspond to the 'Manatee context', and the 'Mirror context'.

Manatee Context (Individuals 333x)

The manatee context has a great quantity, quality, and variety of offerings. It includes shells and snails from both coasts (Gulf of Mexico and the Pacific Ocean), obsidian from the Sierra de las Navajas (Hidalgo, Mexico), local animals such as river fish lynx, armadillo plates, and foreign aquatic species from the deep sea.

Individuals 333A_TOL_a, 333B_TOL_a and 333C_TOL_a 333O_TOL_b, 333Q_TOL_b from this context were sampled for this study. This context is located in building 33 for which dates are available, either by C14 dating, obsidian hydration dating, or the analysis of ceramic materials.

In this context, the more recent burial is dated between 1,300 and 1,400 CE. In this burial we found the individuals 333A_TOL_a, 333B_TOL_a, and 333C_TOL_a. Charcoal was deposited among the sediment that covered the bodies and has been C14 dated to 1,350 ± 50 CE.

Woman 333A_TOL_a was between 25 and 30 years old. She had a pendant made from a lynx tusk placed as a trousseau on her chest, possibly hung with knots since it has no orifice, in addition to a necklace of Pacific olive shells and a human jaw with a perforation as a pendant. As an offering, a piece in the shape of a star or flower made with a spondilus snail, a stingray caudal spine, a natural shell, a fragment of river fish, an obsidian flake, a yew, and a bean-shaped piece with a mixture of cinnabar and almagre were deposited under the body (*7*, *8*).

333B_TOL_a was an infant individual; he was found carefully arranged, leaving the arm in almost total order, first the humerus, radius, and ulna. However, the latter was placed upside down, and in the place of the wrist, a necklace was placed as a trousseau. The necklace comprises four pieces that form a snake, the first of a cylindrical clay bead, another similar one of bone, a flat and square bead of the mother-of-pearl shell to finish with an elongated piece also of mother-of-pearl shell, which represents the rattlesnake's rattle. It is deduced that this burial is of ritual type, since the coxal bones of the child were placed around the circle, and as an offering, there is a complete shell, an earring made with the incisor of a six or seven-year-old child, and a necklace of shells.

Individual 333C_TOL_a was a woman between 20 and 21 years old; she was located to the North. A number of pieces were found with her. Her trousseau was a necklace of 1,150 beads and three snail earrings. Also, as offerings on the body there was a button-like circular piece, an armadillo plaque, an obsidian flake, shuffleboard or ceramic circle, and, next to her to the south, a pectoral made on a manatee rib, which gives its name to this archaeological context.

Other older burials were found in this building. Individual 333O_TOL_b was found next to the platform wall, C14 dated to 700 ± 50 CE. Individual 333Q_TOL_b was found in an older burial C14 dated to 520 ± 40 CE . The platform was to the west of the altar, and the burial was 50 cm deep, it corresponds to the first phase of occupation (*6*).

Mirror context (Individuals 2417x)

Individuals 2417Q_TOL_b, 2417C_TOL_a and 2417J_TOL_a from this context were sampled for this study. This archaeological context is composed of sixteen individuals; it is located in front of the altar of building 24. Among the fill, scattered bones of additional thirty-five people were found. The bones in the fill and the closeness of the primary individuals suggest that the deposition of the primary bodies was at a single time, occupying the bones of previous individuals as fill.

A total of four levels of human body deposition were observed. There is at least one primary body surrounded by secondary bodies, remnants of previous burials, and filler bones in each level. In the first and most superficial level, in front of the altar, a group formed by a woman's primary body (2417C_TOL_a) and the secondary remains of three children (not analyzed in this study) were placed. The remains form a U shape, with the open portion pointing west. The bones of the woman 2417C_TOL_a seem to have received scalding liquids or some other substance, which produces an effect similar to boiling the bones, as they appear crystallized, although additional studies are required to define what was the precise taphonomic process. In addition, she presented pathologies such as spondylitis (the fusion of the last vertebra and the sacrum) and poor dental health, orbital sieve and marks at the insertions of the arms. The woman had as trousseau a necklace formed by sixteen river mollusks of the genus *Unio*, and as an offerings a snail button and obsidian flake and razor (*9–12*). The offering of individual 2417C_TOL_a is significant for its quality, variety, abundance, and treatment given to the corpse.

Five individuals were found in the third level of the northeast of the assemblage. The most relevant is individual 2417J_TOL_a, who presents pathologies such as spondylitis, evident in the sacrum, iliac and lumbar vertebrae. In addition to poor dental health, he also performed exercises that developed his muscles and left a mark on his arms and legs.

In the fourth and last level, the remains of four more individuals were located, all adults, two secondary, and two remnant primary portions. Individual 2417Q_TOL_b corresponds to a remnant primary portion that was placed in the ventral decubitus position (face down) of a male between forty-five and fifty-five years old. Under individual 2417Q_TOL_b, there were two isolated bones, a coxal, and a vertebrae. Also under this individual, a pyrite mosaic mounted on a ceramic base was found, thought to serve as a mirror. The mirror is a significant offering; it is related to the priests and the god Tezcatlipoca, the deity of the underworld and the night, characterized because it carries a mirror, which is also used in the arts of divination, to obtain omens and to provoke evils (*6*).

Individual 6428A_TOL_b

The body of individual 6428A_TOL_b was found in building 64, being part of burial 28. It corresponds to an adult male in a flexed dorsal decubitus position with a significant offering since it has a flint bifacial, two jade beads. All these are allochthonous materials and ten amulets carved in bone (*6*).

Ranas, Sierra Gorda, Querétaro

Ranas archaeological site is located in the Sierra Gorda, in the State of Querétaro, and the occupation of the site is recorded from 400 BCE to 1,300 CE.

Individual 11R_R_b

Individual 11R_R_b was found in a burial that contained three individuals, one primary (11R_R_b) and two secondary arranged on the sides of the primary. 11R_R_b corresponds to an adult male over 35 years old. He was found deposited in dorsal decubitus, with the trunk slightly flexed upwards. The arms were placed in front of the body and the wrists were touching. His hands were on the pelvic region and in the right hand he was holding a ceramic pipe. Two bone circles forming a truncated cone were found in the area of the pubic symphysis. He presented osteological characteristics of repetitive work and high stress to the body. The enteropathies in the long bones of the legs, arm, vertebrae and calcaneus may be directly related to mining. Thus, individual 11R_R_b is inferred to have worked in mining for several years. The measurement of high values of heavy metals in a combination of Pb, Hg, As and Sb indicate chronic exposure to a wide variety of heavy metals present in geological contexts, and which were systematically deposited in the bones of the individual (*13*).
The relative chronology of the excavated materials and those directly associated with a more superficial burial context suggested a chronology between 450 and 700 CE. In terms of Mesoamerican periodicity, it corresponds to the transition between the Classic and Epiclassic periods (*13*).

Individuals 7A_R_b and 3A7I_R_b were found in structure 3A, in burial seven at the archaeological site of Ranas.

Burial 7 is a complex burial, where initially, five areas of intrusion are defined, and inside each of them, there is evidence of skeletal remains of one or more individuals.

Individual 7A_R_b

This individual is the most important since it is located just below the altar. It is a direct primary burial with a very flexed seated disposition. The individual was deposited as a mortuary bundle strongly tied (at the height of the shoulders and waist), so when the body decomposed in its taphonomic process, the head fell between the legs. The care taken in the deposition and the fact that limestone slabs covered the individual from its lowest level helped to keep most of the skeleton in its primary position, where the only element of post-depositional alteration was a rodent burrow that altered the metatarsal bones and phalanges of the left foot and altered the distal phalanges of the right foot. The burial had two obsidian microliths associated (a portion of a prismatic blade manufactured in green obsidian from the Sierra de las Navajas in Hidalgo, Mexico). One of them had abrupt lateral retouch, very worn because it is an instrument for cutting hard plant materials (due to the striations and scratches on the ventral face of the piece and the extension of the micro polishes that exceed 0.5 mm in extension). In comparison, the second piece of carved lithic is manufactured on a gray veined obsidian flake with physical characteristics very similar to the deposits of Zacualtipan, Hidalgo. From the morphological and functional point of view, this piece was part of a scraper on a percussion blade, although it does not show as much wear due to use as the green obsidian piece.

Individual 3A7I_R_b

The case of individual 3A7I_R_b is another burial of an important personage since it is the only one deposited in an extended form with the head to the south, its face looking to the east, with the right arm flexed on its chest. The hand was deposited on its left shoulder, while its left arm rested on the belly, and its hand is extended between the right ribs and the right humerus, with the torso slightly leaning forward. The pelvic region was horizontal on the ground while the legs were separated with the right knee to the east, the left knee to the west, and the feet under the coxae. Therefore, this is a primary burial, in dorsal decubitus slightly raised from the back and head, with legs flexed on themselves. As it is a primary individual, the different body portions could be recognized and recovered, although in many cases, they were found in regular and poor preservation.

It is essential to mention that two individuals were removed for the deposition of the individual 3A7I. As part of the trousseau, a fragment of fine-grained light gray vesicular basaltic rock (fragment of a metate), remains of the body of a smoothed orange clay pot (a domestic Soyatal vessel) and portions of the flat bottom of a smoothed brown clay vessel (of the Trejo pyrite smoothed type) were deposited. A relevant element is a delimitation with red ferruginous rocks to the cist of the individual on its east, west, and north sides. It is important to note that this type of rock is found only in the region of "El Doctor" (west of Ranas and Toluquilla) or more than 7 km north in the area of San Cristobal and "El Apartadero" in the Sierra Gorda.

This individual had an offering in his chest and neck: 1) A set of 4 bone punches (4.02 and 5.1 cm long by 0.4 cm of maximum thickness) finely worked in bone, well-polished, and evidence of being hardened by fire. Due to their structure, they may be of animal origin. 2) A large necklace of 6 pieces of Oliva spicata snails. 3) Two unfired clay pipes, manufactured in very dark gray granular clay without firing, corresponding to 2 different types. One angled bowl and the other plate. The cupped one measures 6.5 cm long and 3.7 cm high in the cup. While plate one is 4.6 cm long and 3.1 cm high, they are not functional. They are votive elements. The bowl pipe corresponds to the tradition in the Huasteca region, while the plate pipe is better represented in the lowlands and western Mexico and is present in archaeological contexts from 650 to 1,000 CE.

Cañada de la Virgen, Guanajuato

Individual E2_CdV_b

Direct secondary burial, located in layer III of Table V-4 (although it partially intrudes up to layer IV). It presents only proximal fragments of incomplete femurs corresponding to an adult individual of undetermined age. Although it is possible that its partial intrusion into floor 1 and layer IV is accidental, the only difference with respect to the rest of the skeletons from Room 3 is the fact that it does not have contact with floor 2, but is found right at the level of floor 1. It seems more likely that this burial, due to its position and secondary character, functioned as a ritual seal for floor 2, in a termination ritual aimed at "killing the funerary space between the two floors (*14*).

Individual E4_CdV_b

Located in layer IV of Table X-4. It is an indirect primary burial deposited in a cist in extended dorsal decubitus facing east. The estimated age is 5 years +/- 24 months and sex indeterminate. It presents 60% of the skeleton with absence of bones of the feet, patella, sacrum and coccyx (*14*).

Individual E6_CdV_b

Located in layer IV of Table V-3. Indirect secondary burial in cist, corresponding to a male individual of 40-45 years of age. It presents only fragmented iliacs, complete right radius, proximal part of both femurs at 50% and part of the proximal portion of the right tibia. Position and orientation not discernible (*14*).

Individual E7_CdV_b

Located in layer IV of Table X-3. Indirect primary burial deposited in cist in dorsal decubitus flexed oriented to the east. Corresponds to a female subject of 25 to 30 years of age. Feet and hands are missing (*14*).

Individual E8_CdV_b

Located in layer IV of table U-3. Direct primary burial in flexed dorsal decubitus oriented to the east. It corresponds to a male individual between 35 and 40 years old. He presents 90% of the skeleton with absence of feet (*14*).

Individual E9_CdV_b

Located in layer IV of Table T-4. It is an indirect primary burial deposited in flexed dorsal decubitus with orientation to the east. It corresponds to a male subject of 20 to 25 years of age. It presents 80% of the skeleton with absence of feet (*14*).

Individual E10_CdV_b

Located in layer IV of box V-3. This is a direct secondary burial with no discernible anatomical position. It consists only of a left femur diaphysis and a mandible (*14*).

Individual E11_CdV_b

Located in layer IV of squares U/V-3 and 4. It is a direct secondary burial formed by three fragments of long bones, fragments of ribs, a clavicle, some phalanges, fragments of pelvis and skull, as well as some molars (*14*).

Individual E14_CdV_b

Located in complex C, which is a circular structure, with an association to the wind and related to the North Star and Ursa Minor, which is linked to Tezcatlipoca. This burial was found at the foot of the access ramp to the second structure (*14*).

Individual E15_CdV_b

This individual was called the "girl of the rain". This subadult individual, through osteological method was estimated to be approximately 7 year old and possible a female, but we take this with caution since the individual is a subadult. She was found in complex B, it was found linked to the pluvial drainage, which is large and well-constructed with superimposed slabs. She was 1.10 m tall and is thought to be part of the ancestral veneration practiced in the place, since she was accompanied by a coyote and placed in such a way that it was necessary to decompose her body in order to accommodate her seated and compressed. Possible post mortem treatment in the temazcal that is at the top of the pyramidal base.

The skeleton is female and corresponds to a seven-year-old girl, deposited in the center of a circle of stones and accompanied by a ceramic offering of cajetes and plates. The girl of the rain, carried in her neck a small necklace of beads that includes a bead in the form of butterflies in the central part of the necklace.

The offering is completed with a ceramic spindle and a fragment of marine shell. Next to the girl and as a companion and vestige of the practices of ancestral veneration, a small coyote was placed -a complete specimen of Canis latrans-, an animal of totemic importance in the region.

Forensic studies performed on the girl of the rain provided her height, estimated at 1.10 centimeters, and some pathologies associated with infectious processes such as periostitis or infection of the periosteum, as well as cranial lesions probably diagnostic of tuberculosis (*14*).

Individual E18_CdV_b

"The decapitated one". Part of his burial offering consisted of an elongated handle smoker.

It was found associated with the enclosed patio located on the elevated platform on one side of the pyramidal base of Complex B.

The individual is male, aged between 20 and 30 years and with an unusual height of 1.73 centimeters - the left femur itself measured 45 centimeters. Forensic analysis indicates evidence of disease, specifically ankylosing arthritis, as well as apparent postmortem decapitation (*14*).

Individual E19_CdV_b

According to the skeleton, the burial of E19_CdV_b corresponds to a female individual, 20-25 years old. It was identified as an indirect primary burial in a dorsal decubitus position (slightly to the right) flexed. This burial presented a series of alterations regarding the original position, and the hands were absent. Because there are no marks of violence or dismemberment, archaeologists suggest that the most likely hypothesis is that these elements were removed during some looting and were subsequently lost by agricultural work or by the action of rodents. The remains of this burial were found under a layer of killed pottery, and associated with it were the remains of a canid (Canis familiaris) (*15*).

Michoacán

Eight individuals were retrieved from the state of Michoacán. Individuals E1A_Mich_b, E2_Mich_b and E4_Mich_b from the site La Mina; individual M1_Mich from the site Tanhuato and Individuals M2_Mich_b, M3_Mich_b, M4_Mich_b and M5_Mich_b from the site Zaragoza.

La Mina, Michoacán

At the end of December 2014, the INAH Michoacán Center was notified of the discovery of a series of skeletons located in the construction of a perimeter wall in the Telesencundaria #133 school, in the town of La Mina, municipality of Álvaro Obregón, Michoacán.

Upon arriving at Telesecundaria School #133, the archaeologists detected that the funerary context had been seriously altered due to the construction of a trench for the foundation of the perimeter wall. From the initial reports obtained from the masons, it was possible to determine that the bodies were concentrated in the same sector.

During the first visit, towards the end of December 2014, a collection of those osteological elements extracted due to the construction works was carried out.

So far, the extension of the site of La Mina is unknown due to the lack of monumental architecture and the growth of the current settlement, which occupies much of the area with vestiges. Excavations have shown that La Mina appears to be on what was once the shores of a village founded on the southwestern shore of Lake Cuitzeo.

The absence of architectural elements on the surface, in the intervened area, and the surroundings, suggests an architecture based on perishable materials. The bioarchaeological remains were located in the northern sector of an embankment covering an area of approximately 1200 m2. The ceramic materials on the surface indicate that the settlement was occupied between 200 B.C. and 900 A.D. Although this data indicates an extensive temporal extension, the stratigraphic unit indicates a single moment of occupation.

The excavations carried out at this site were only aimed at recovering the bioarchaeological materials affected by the construction work, so the archaeologists do not rule out the existence of more burials at the site.

E1A_Mich_b

Individual E1A_Mich_b was in a poor state of preservation. Only part of the lower limbs, two-thirds of the mandible, and the upper dental arcade were located. The bad preservation limited the identification of the sex of the individual based on its skeleton. This individual corresponded to a young adult placed in a right lateral decubitus position and was oriented in a west-east direction.

The grave goods consisted of a vessel of the Paso Ancho/graffito type placed at the level of the lumbar vertebrae and showed clear traces of use. Two reused sherds deposited in front of the face were other elements that completed the offering. The intentional placement of this ceramic fragment leads us to suppose that it is an object used in the development of the funerary ritual and later placed as part of the trousseau. These two ceramic fragments were dated together with another sample from another burial using the archaeomagnetic technique. The results indicated that the burial corresponded to 647-825 CE (*16*) during the so-called Epiclassic period.

A third element that made up the offering consisted of a dog in its juvenile development phase. Despite its poor state of preservation, it was possible to identify that the animal was not more than eight months old, considering the process of ossification of the epiphyses. From this specimen, it was possible to appreciate a series of taphonomic alterations that indicated the animal's treatments prior to being placed in the funerary context, specifically areas with exposure to heat in the tibia and the area of the maxilla (*17*).

E2_Mich_b

Individual E2_Mich_b corresponds to an early infant who was approximately six months old. This infant did not present any artifact or other element that could be identified as part of an offering. The infant was buried at a greater depth than the rest of the individuals, approximately 1m, even surpassing the tezontle layer that formed the leveling embankment.

E4_Mich_b

The individual E4_Mich_b corresponds to a skull collected at the site without context because it was removed by personnel building the school's perimeter fence. This skull was placed next to the burial of individual E1A_Mich_b.

Tanhuato, Michoacán

M1_Mich corresponds to the burial excavated in a rescue excavation in the town of Tanhuato, Michoacán, during the construction of a canal in a street inside the town. Tanhuato is located in the municipality of the same name in the state of Michoacán. The burial was found at 1,543 meters above sea level without a greater context, since it is located in an urbanized area. The surrounding area is almost flat and there is a large amount of cultivated land. The burial was mixed, and the inhumations were made in an enclosure dug in the soil matrix, whose bottom is one meter deep with respect to the level of the street; it covers an area of about 5 $m^2$. Primary and secondary burials were found in the pit and two incomplete individuals, two ossuaries and a funerary pot could be identified. Unfortunately, both bones and materials had already been removed by construction workers, so the context was very altered. Absolute dates for this individual are not yet available.

Zaragoza, Michoacán

The Zaragoza site is located in the municipality of La Piedad, on a flat-topped, isolated elevation called Mesa Acuitzio. This site presents a large rocky front that the settlers of Zaragoza took advantage of as a support for their settlement. The monumental area of the site was built on a relatively flat part located more or less halfway up the mesa, around 1,753 meters above sea level, while the living and cultivation area is distributed downwards, both to the east and north, along the slopes through artificial terraces.

Samples M2_Mich_b and M3_Mich_b come from a mixed burial, excavated in 2004 by Anyul Cuellar and Diana Bustos, called Burial 1. The space was used in successive events, that is, not all the bodies were deposited there simultaneously. It is located outside, on the east side, of an architectural unit that we have identified as a possible temascal (low heat sweat lodge), located at the southern end of one of the main group of structures. Among the buildings that make up this complex, a ball court stands out, almost 57 meters long (including the headers) and 14 meters wide.

A total of 14 individuals were located in the funerary context, identified from the number of skulls, i.e., not necessarily complete bodies were found. The remains were in a very poor state of preservation, but at least 5 primary deposits were identified. M2_Mich_b comes from individual 7, located at the western end of the pit, next to the wall that delimits the east side of the temascal. M3_Mich_b corresponds to a second molar of skull 13. Both skulls were found without anatomical connection, although they were part of the group of bodies deposited in the burial space.

M4_Mich_b and M5_Mich_b, also from the Zaragoza/Cerro de los Chichimecas site, but from a different burial, called Burial 2, were also mixed and excavated in 2012 by Alfredo Salas. This burial is located to the south of structure 1, which is part of the same complex as the court, to the east of it. The pit, of 24 $m^2$, presents a total of 8 or 9 individuals, which are not specified yet. The samples correspond to a first molar of M4_Mich_b and a third molar of M5_Mich_b, respectively. Individual M4_Mich_b belongs to an individual placed in extended dorsal decubitus with a south-north axis, slightly deviated from 6 to 8 degrees to the east; it is important to note that this is the only individual with this orientation inside the pit. This individual had an associated offering composed of a cajete (ceramic bowl), a pot and a "Capiral" lid. Individual M5_Mich_b was the most complete skeleton, although in a poor state of preservation; M5_Mich_b was found in an extended dorsal decubitus position, with arms flexed and with an east-west orientation with a 287° deviation with respect to north. Three pots were found associated with this individual, two of them miniature with a composite silhouette; a cajete, a basket-type vessel with zoomorphic handles, a spindle whorl, a tejolote and a tripod cajete, located at the level of the left tibia; also associated with this individual are a fragment of another cajete, a spindle whorl and a tripod cajete, but those located to the left of the skull.

Mitochondrial haplogroups identified

In the Sierra Tarahumara, we found two sub-linages of the mitochondrial haplogroup C (C 100%), as previously reported. In contrast, Central Mexico shows a higher diversity of mitochondrial haplogroups. Cañada de la Virgen was the site with the highest number of sub-haplogroups, distributed in B 50%, A 25%, C 12%, and D 12%. Michoacán presented four different sub-haplogroups (A 40%, D 40%, and C 20%). Sierra Gorda also presented four different sub-haplogroups but corresponding to major lineages A 50%, B 43%, and D 7%, respectively.

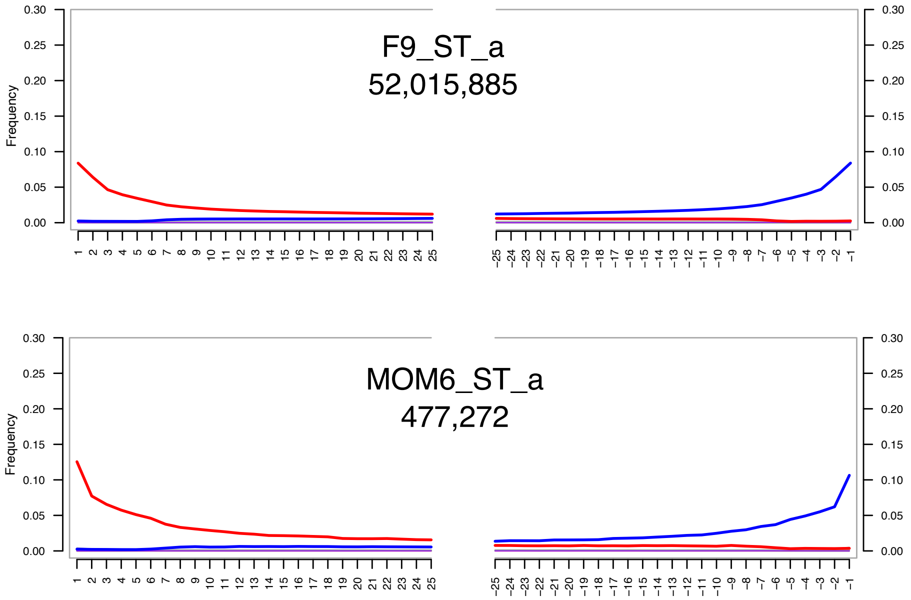


**Fig S1.** Damage patterns of the genomic data from Sierra Tarahumara pre-Hispanic individuals as inferred by mapDamage (*18*). The number of sequences included in the analysis is indicated below the individual's ID. The red color indicates the frequency of cytosine to thymine changes, while the blue color indicates guanine to adenine changes. The x-axis shows the position of the base within the read, in the direction 5' to 3'. The y-axis shows the frequency of such changes.

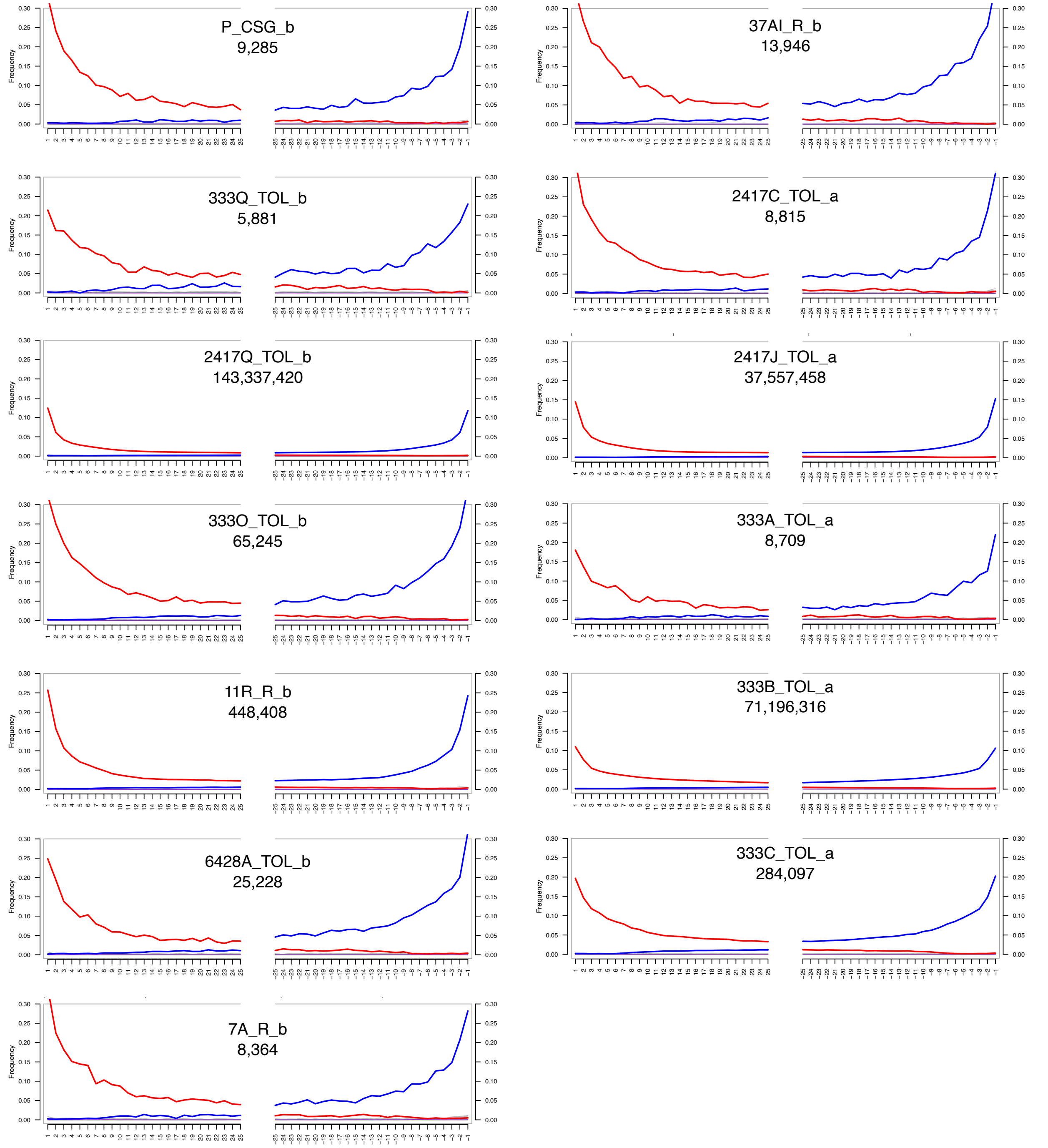


**Fig S2.** Damage patterns of the genomic data from Sierra Gorda pre-Hispanic individuals as inferred by mapDamage (*18*). The number of sequences included in the analysis is indicated below the individual's ID. The red color indicates the frequency of cytosine to thymine changes, while the blue color indicates guanine to adenine changes. The x-axis shows the position of the base within the read, in the direction 5' to 3'. The y-axis shows the frequency of such changes.

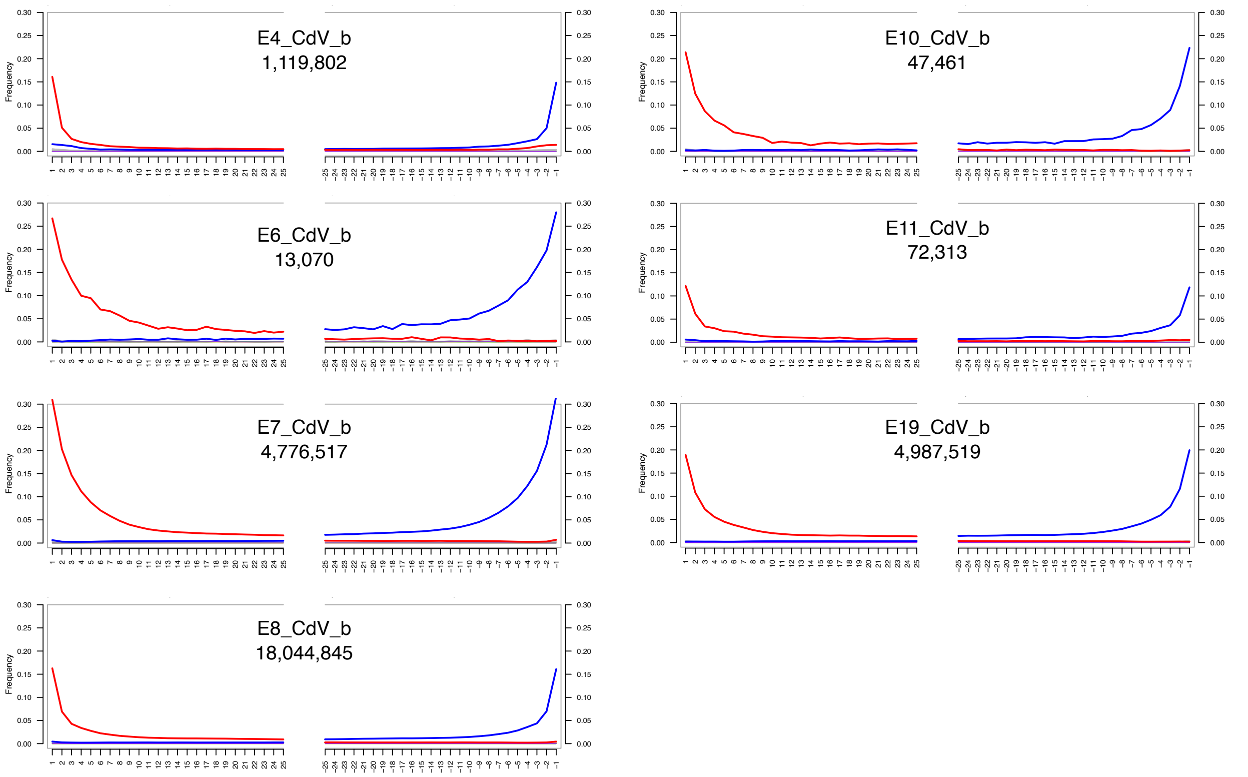


**Fig S3.** Damage patterns of the genomic data from Cañada de la Virgen pre-Hispanic individuals as inferred by mapDamage (*18*). The number of sequences included in the analysis is indicated below the individual's ID. The red color indicates the frequency of cytosine to thymine changes, while the blue color indicates guanine to adenine changes. The x-axis shows the position of the base within the read, in the direction 5' to 3'. The y-axis shows the frequency of such changes.

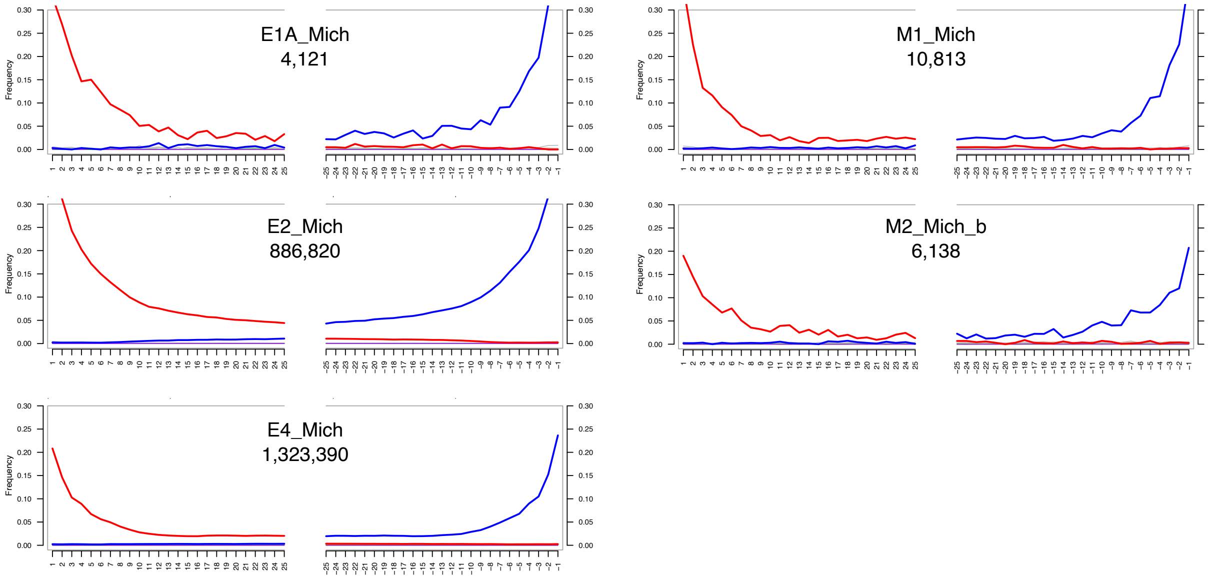


**Fig S4.** Damage patterns of the genomic data from Michoacán pre-Hispanic individuals as inferred by mapDamage (*18*). The number of sequences included in the analysis is indicated below the individual's ID. The red color indicates the frequency of cytosine to thymine changes, while the blue color indicates guanine to adenine changes. The x-axis shows the position of the base within the read, in the direction 5' to 3'. The y-axis shows the frequency of such changes.

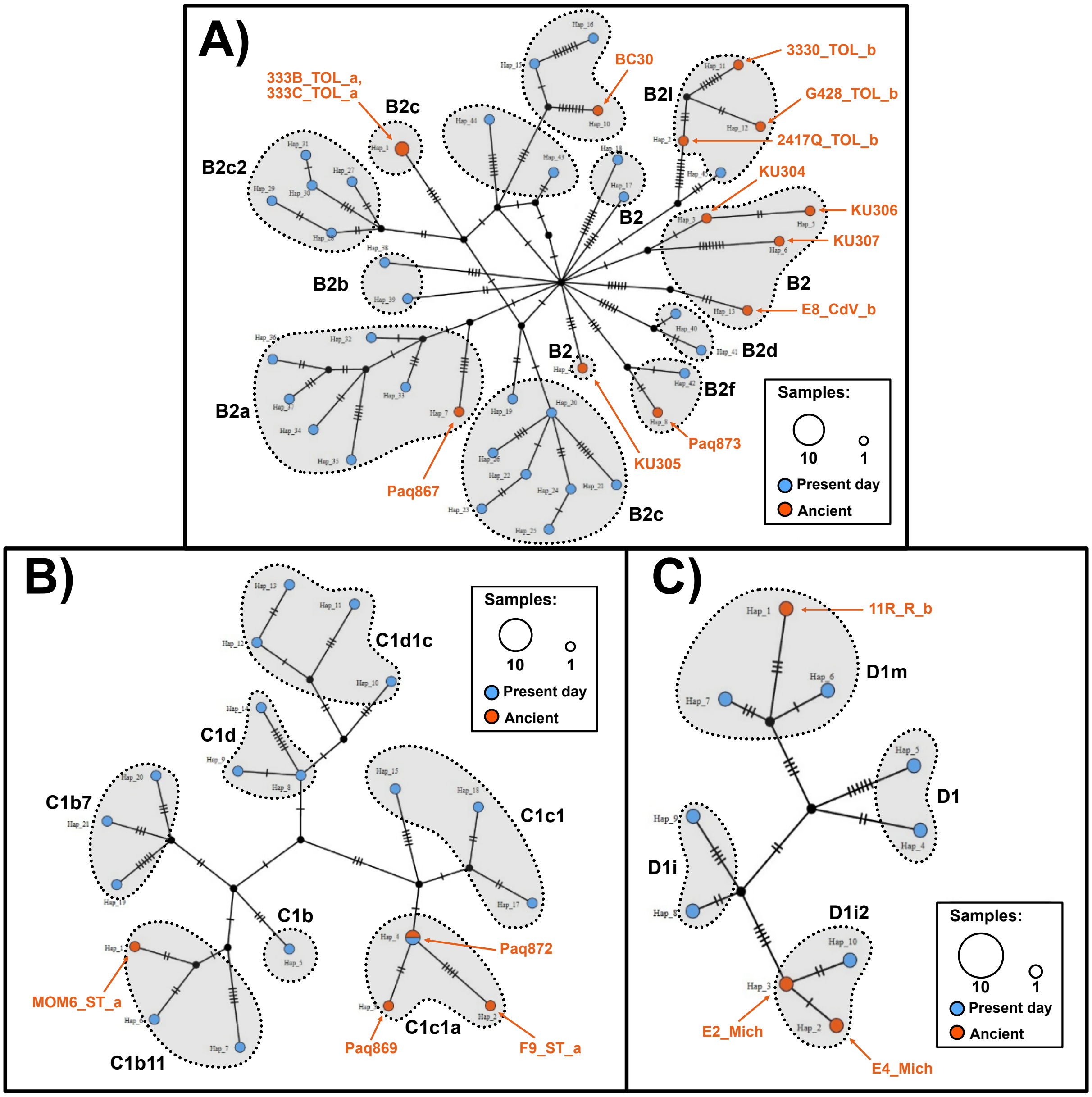


**Fig S5.** Haplotype Networks from the mitochondrial haplogroups of pre-Hispanic individuals from Mexico from this and previous studies. a) Haplogroup B, b) Haplogroup C and c) Haplogroup D. Sub-haplogroups are shaded with gray. Individuals from the same site tend to cluster together.

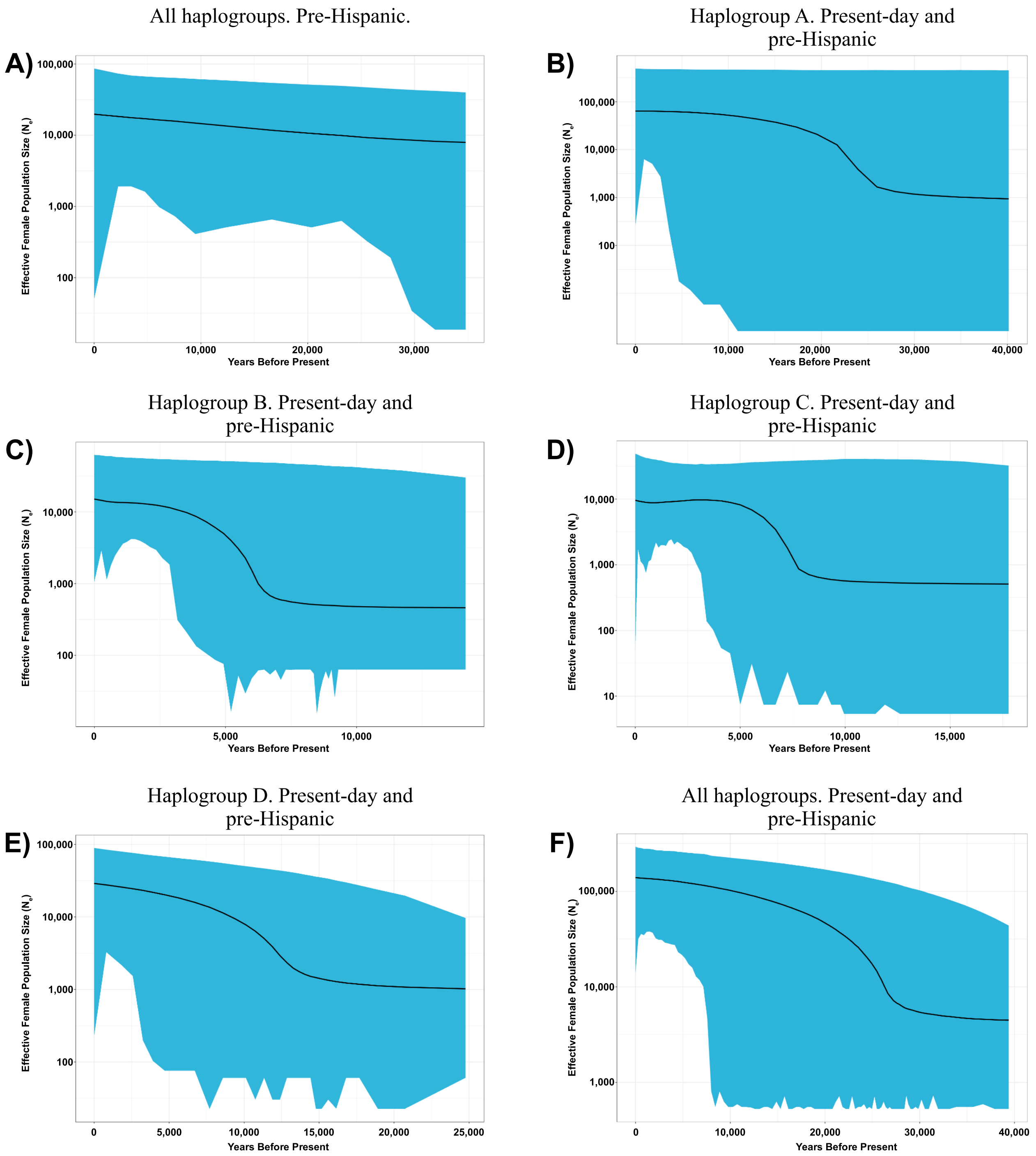


**Fig S6.** Extended Bayesian skyline plot (EBSP) constructed with ancient and present day mitochondrial DNA. The credibility interval of 95% is shown in blue, and the black line represents the median of the mutation rate. A) EBSP of all pre-Hispanic mitochondrial haplogroups (A, B, C, and D). B) EBSP of present-day and pre-Hispanic mitochondrial haplogroup A. C) EBSP of present-day and pre-Hispanic mitochondrial haplogroup B. D) EBSP of present-day and pre-Hispanic mitochondrial haplogroup C. E) EBSP of present-day and pre-Hispanic mitochondrial haplogroup D. F) EBSP of all mitochondrial haplogroups (A,B,C and, D) including present-day and pre-Hispanic mitochondrial sequences.

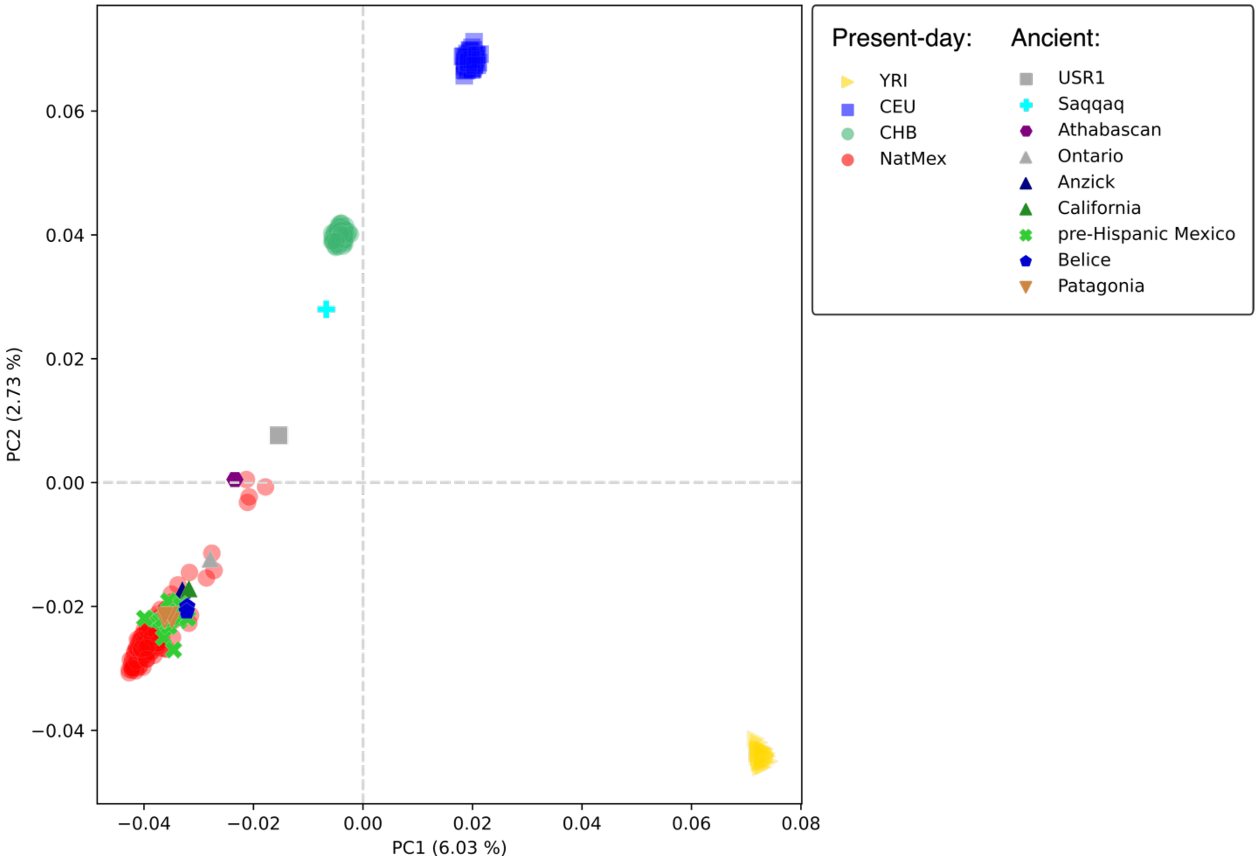


**Fig. S7.** Principal Components Analysis including pre-Hispanic individuals from Mexico, ancient individuals from the Americas (Supplementary Table S1 and S3), Indigenous populations from Mexico (NatMex) from **(*19–21*)**, and continental reference populations from Africa (YRI), Europe (CEU), and Asia (CHB) from 1000 Genomes **(*22*)** . Pre-Hispanic individuals from Mexico cluster closely with NatMex. All ancient individuals were projected into the PC space using the lsqproject function from the eigensoft package.

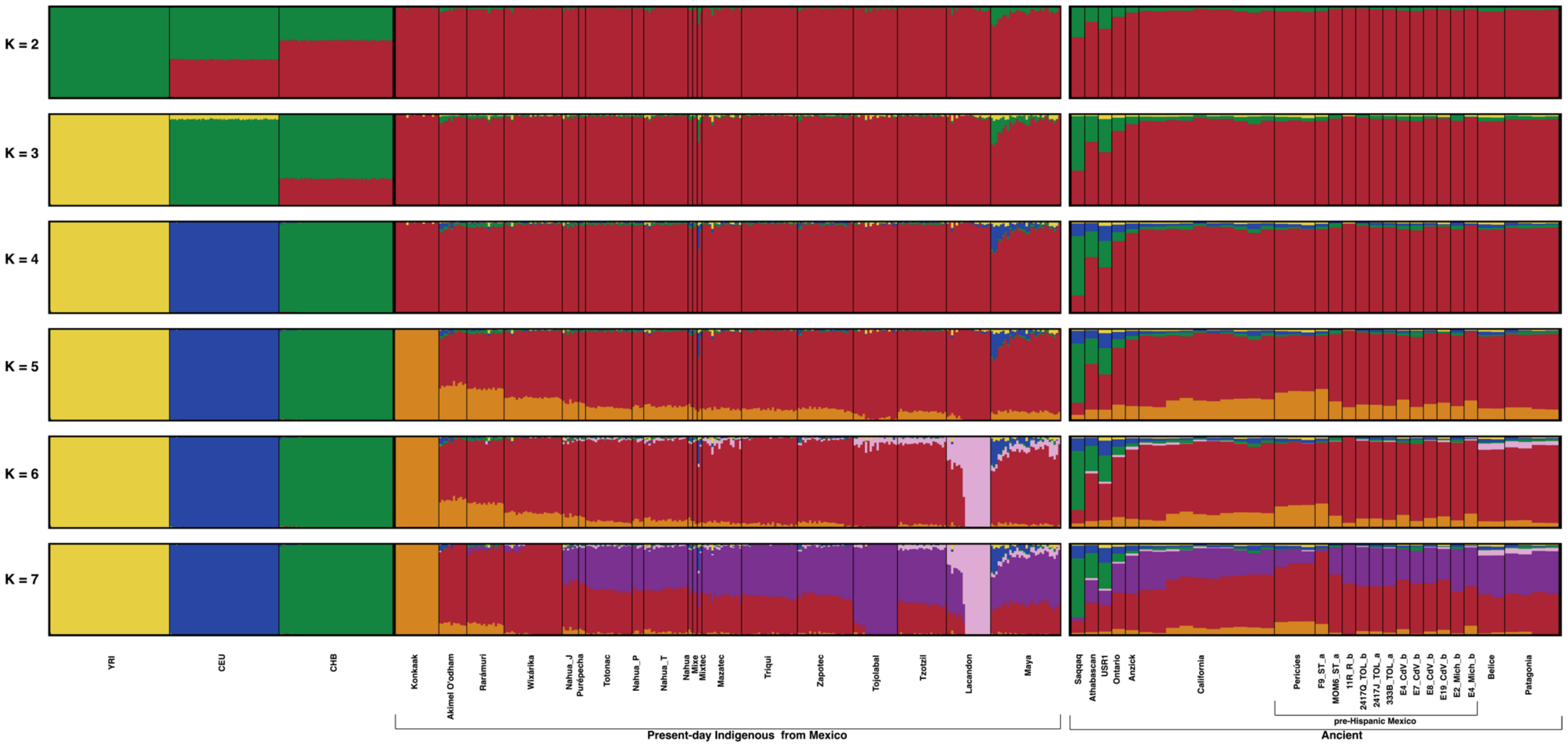


**Fig. S8.** ADMIXTURE analysis with pre-Hispanic individuals from Mexico and America, present-day Indigenous populations from Mexico, and continental references from Africa (YRI), Europe (CEU), and Asia (CHB) from 1000 Genomes ***(22)*** . Ancient individuals from the Americas are on the right, sorted from north to south. K=4 (the K value with the lowest CV error) shows that pre-Hispanic individuals from Mexico have similar ancestral components to present-day Indigenous populations from Mexico.

A)

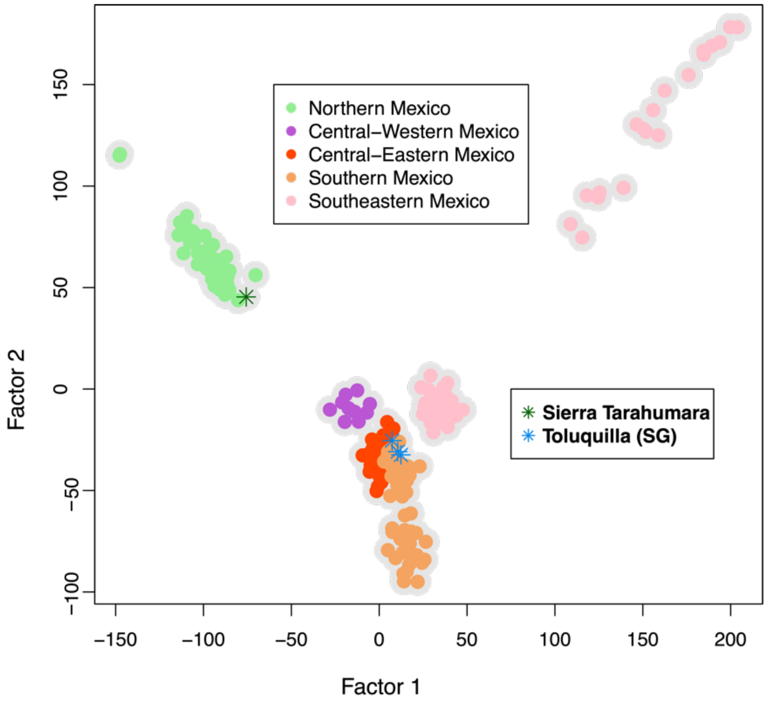


B)

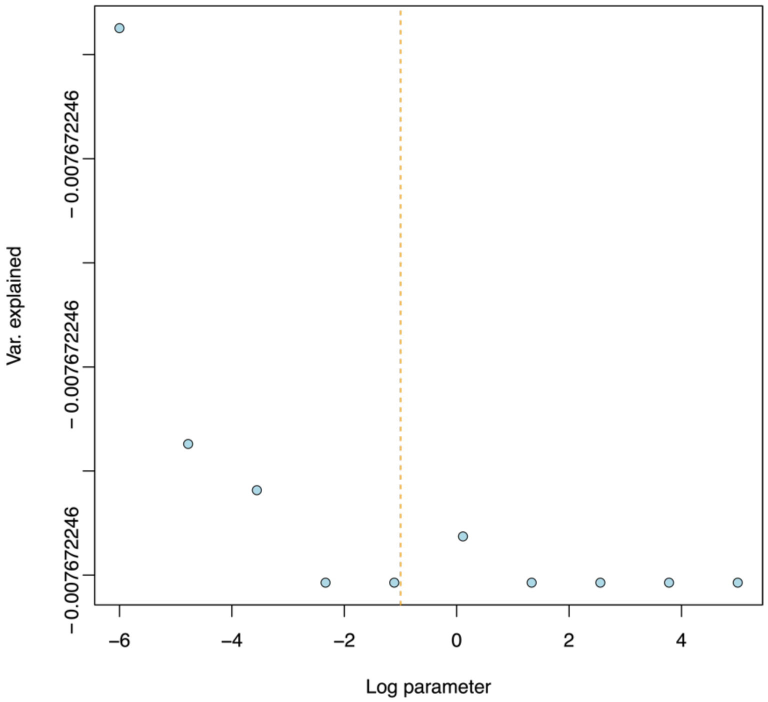


**Fig. S9.** Temporal factor analysis (TFA) of populations from Mexico. A) TFA includes the four pre-Hispanic individuals, F9_ST_a from Sierra Tarahumara and 2417Q_TOL_b, 2417J_TOL_a 333B_TOL_a from Toluquilla. The plot shows the individual F9_ST_b cluster with populations from northern Mexico and Toluquilla clusters close to central-east Mexico populations. B) Percentage of variance explained in the y-axis by different lambda values. The x-axis shows the lambda values in a log scale. A decrease in variance from right to left represents the lambda value used to correct temporality. However, the small range of variances and the absence of a downstep suggest there is no considerable drift explained by time to be corrected by a specific lambda value.

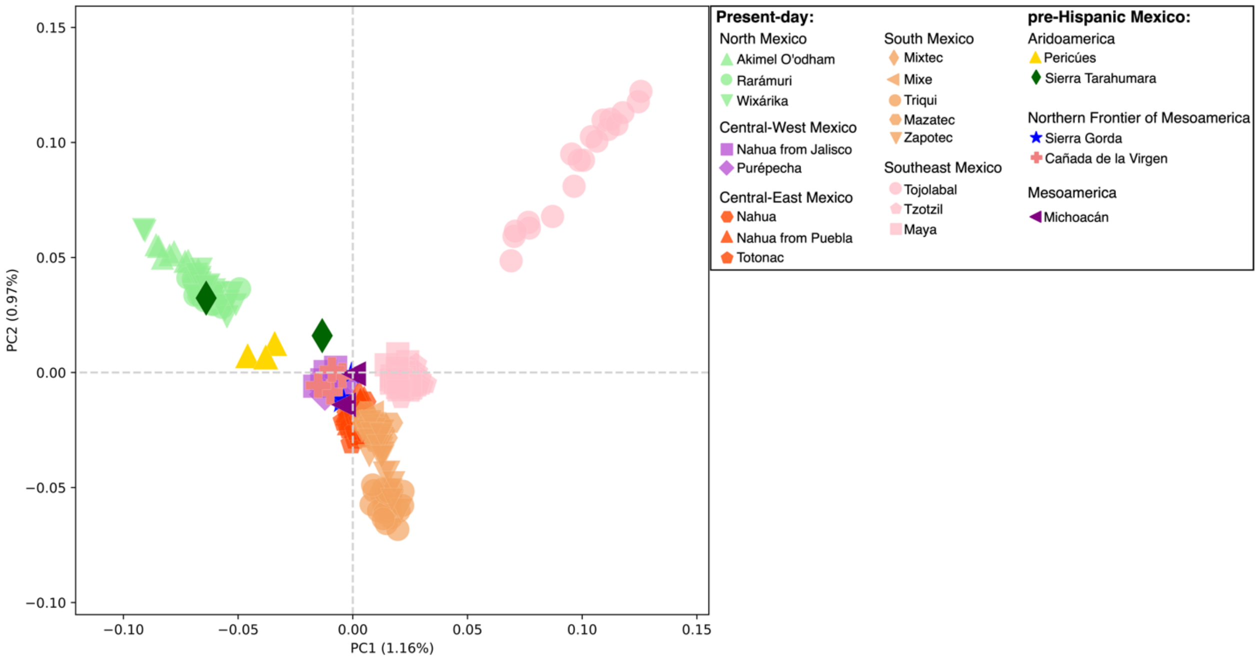


**Fig. S10.** mdPCA of populations from Mexico. The analysis includes the reference panel of present-day Indigenous Populations (see methods) and the genomic data of the pre-Hispanic individuals from Mexico. Individual F9_ST_a from Sierra Tarahumara clusters with present-day populations from northern Mexico, while individuals from CdV and MOM6 cluster close to central-west present-day populations. SG and Michoacán individuals clusters between central-west and central-east present-day populations.

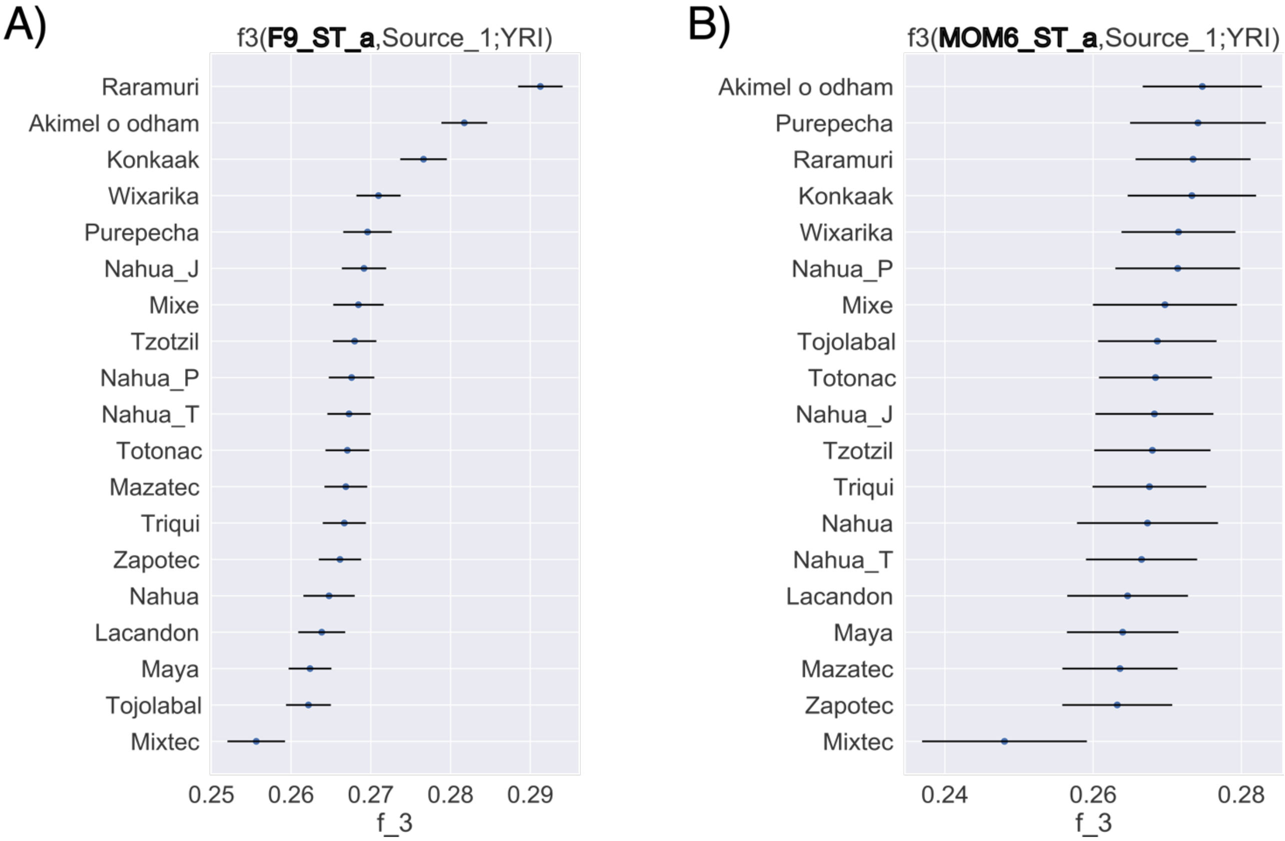


**Fig. S11.** Outgroups f3 statistics for pre-Hispanic individuals from Sierra Tarahumara and other present-day populations of Mexico. A) Mummy F9_ST_a and B) Mummy MOM6_ST_a. Higher values of f3 indicate higher shared genetic drift. Point estimates and one standard error are shown. The individual F9_ST_a have a higher genetic drift shared with present-day Rarámuri population from Northern Mexico. While individual MOM6_ST_a have a similar genetic drift shared with all present-day Indigenous populations from Mexico except with Mixtec.

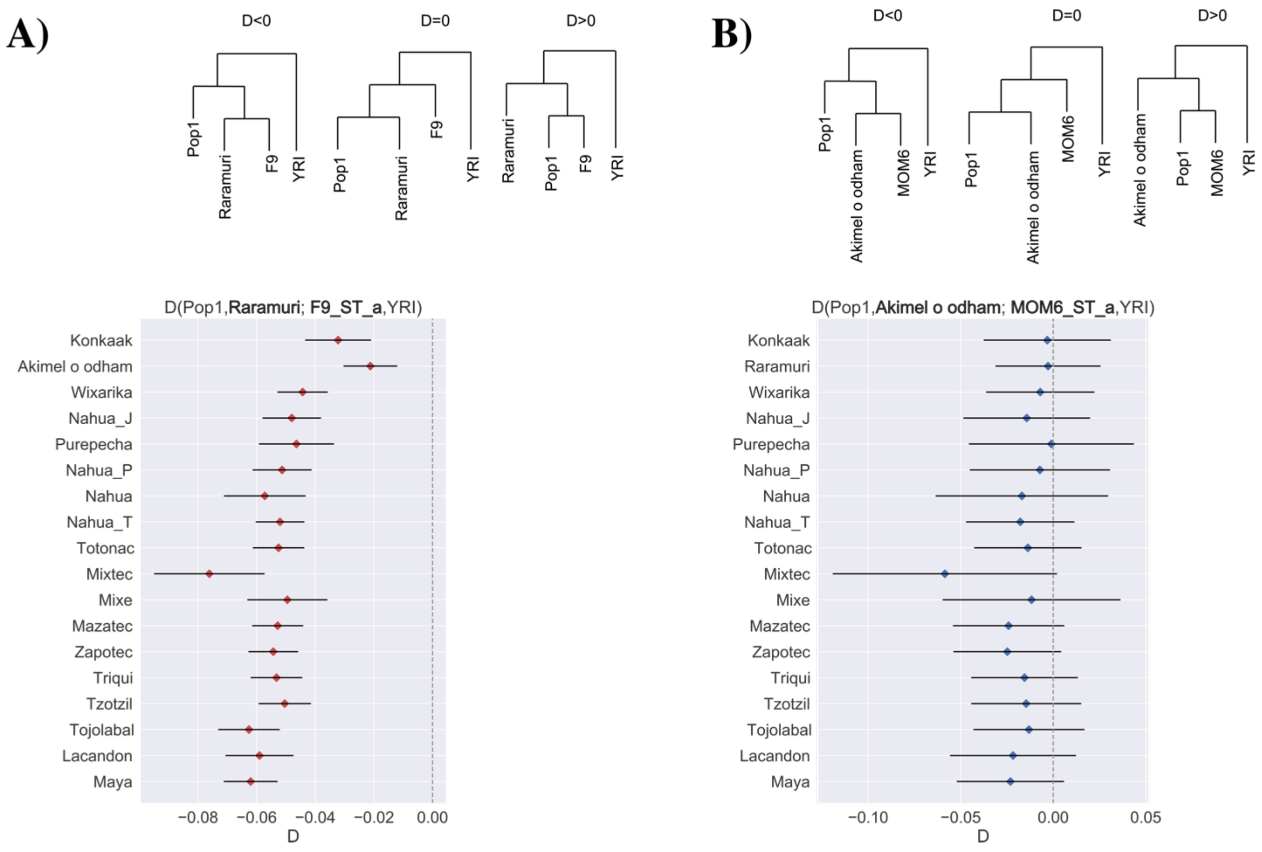


**Fig. S12.** D statistics for pre-Hispanic individuals from Sierra Tarahumara and other present-day populations. A) D statistics in the form D(Pop1, Rarámuri; F9_ST_a, YRI), B) D statistics in the form D(Pop1, Akimel o'odham; MOM6_ST_a, YRI). Expected tree topologies according to D value are drawn on the top of the plot, individual IDs in the trees are indicated with no suffixes. Red dots indicate significant deviations from D=0 (|Z|>3). The individual F9_ST_a shows a significantly higher relationship with present-day Raramuri then with any other present-day population. The individual MOM6_ST_a seems to be equally related to all present-day Indigenous populations from Mexico.

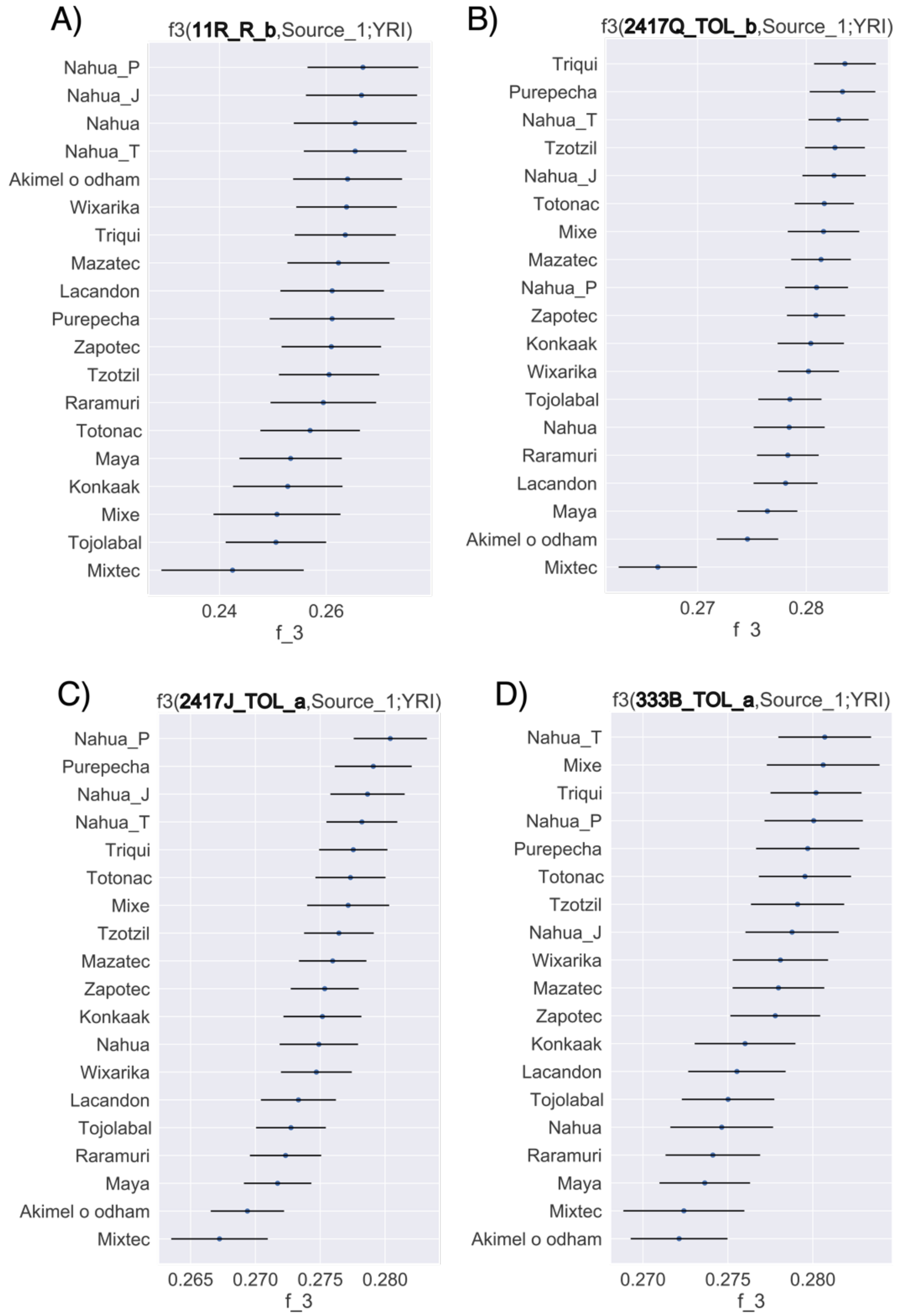


**Fig. S13.** Outgroups f3 statistics for pre-Hispanic individuals from Sierra Gorda and present-day Indigenous populations. A) 11R_R_b, B) 2417Q_TOL_b, C) 2417J_TOL_a ,and D) 333B_TOL_a. Higher values of f3 indicate higher shared genetic drift. Point estimates and one standard error are shown. All individuals show a similar genetic drift shared with all present-day Indigenous populations from Mexico.

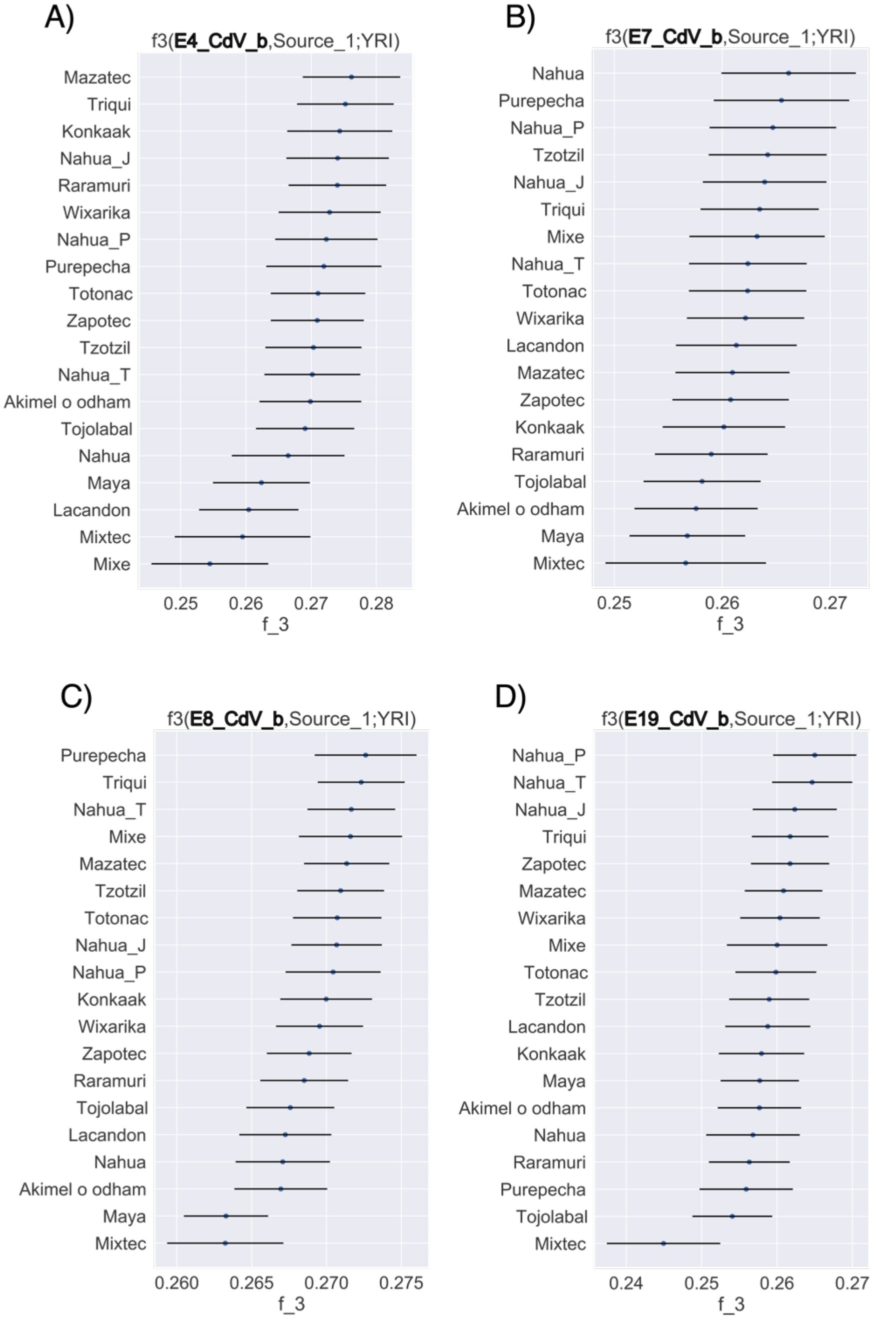


**Fig. S14.** Outgroups f3 statistics for pre-Hispanic individuals from Cañada de la Virgen and present-day Indigenous populations. A) E4_CdV_b, B) E7_CdV_b, C) E8_CdV_b ,and D) E19_CdV_b. Higher values of f3 indicate higher shared genetic drift. Point estimates and one standard error are shown. All individuals show a similar genetic drift shared with all present-day Indigenous populations from Mexico, except with Mixtec.

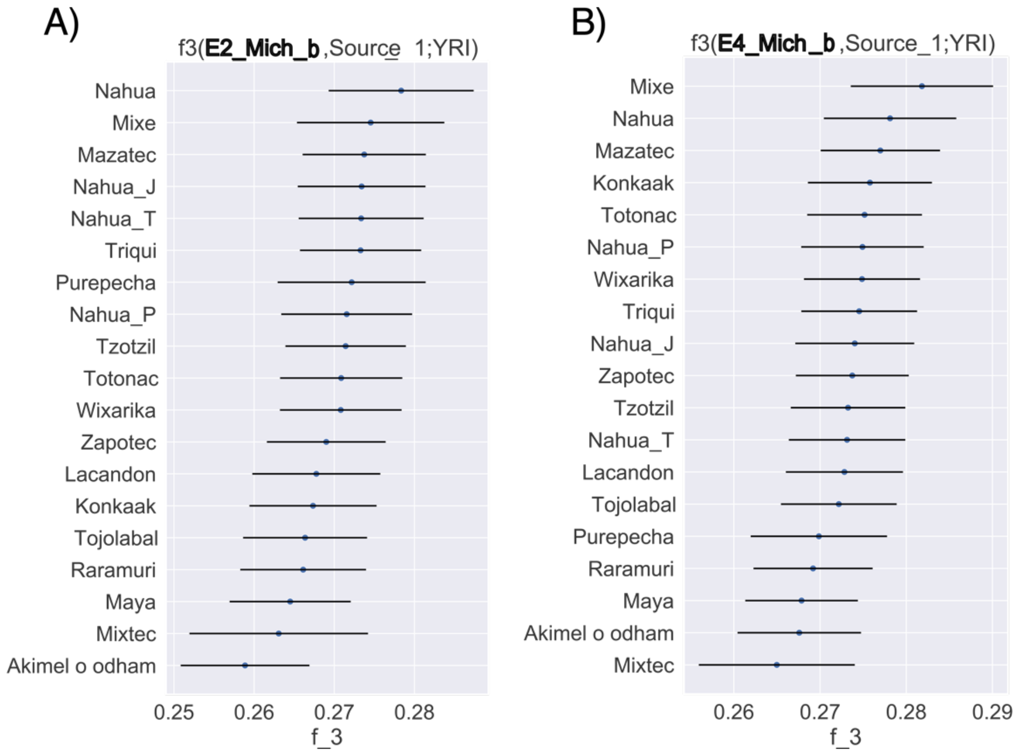


**Fig. S15.** Outgroups f3 statistics for pre-Hispanic individuals from Michoacán and present-day Indigenous populations. A) E2_Mich_b, and B) E4_Mich_b. Higher values of f3 indicate higher shared genetic drift. Point estimates and one standard error are shown. Both individuals show a similar genetic drift shared with all present-day Indigenous populations from Mexico.

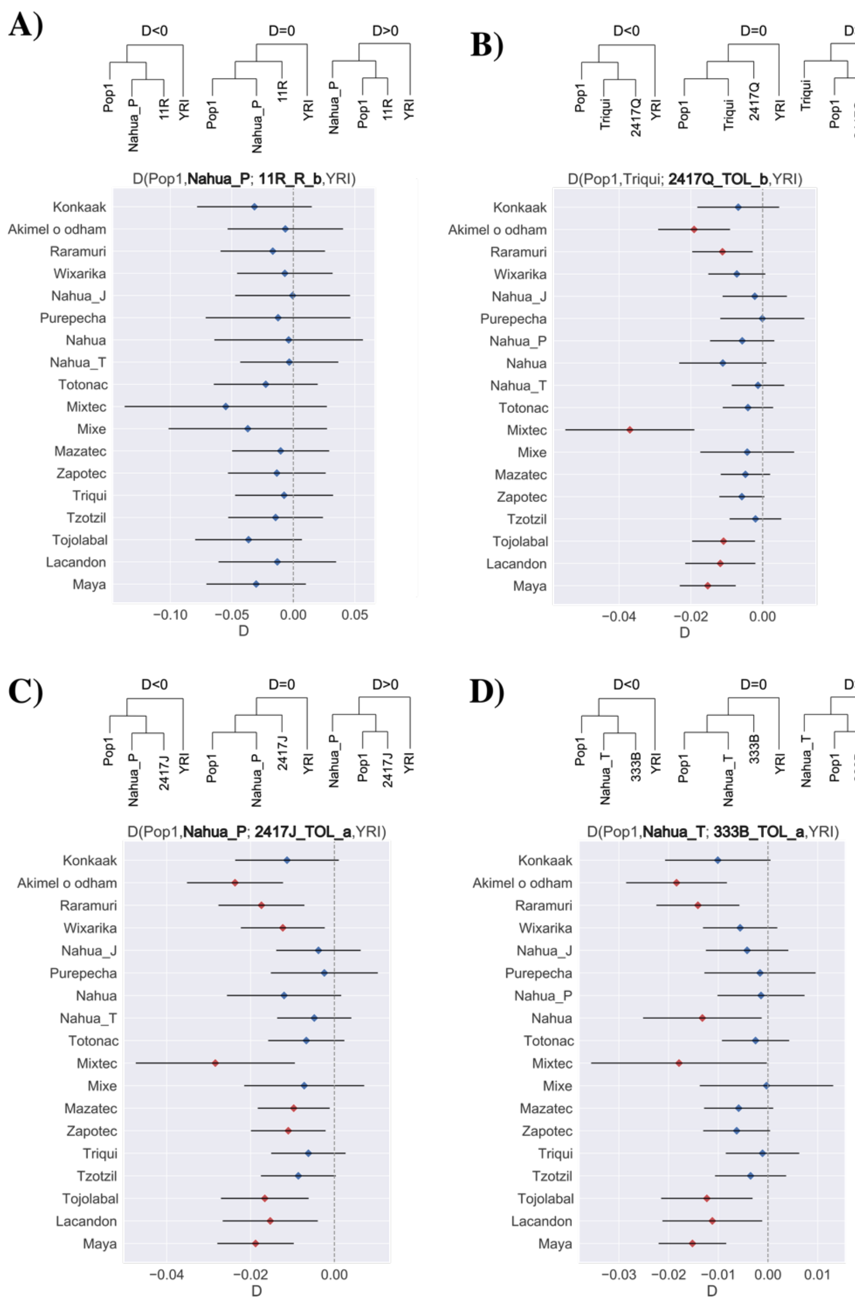


**Fig. S16.** D statistics for pre-Hispanic individuals from Sierra Gorda and present-day Indigenous populations. A) D(Pop1, Nahua_P; 11R_R_b, YRI), B) D(Pop1, Triqui; 2417Q_TOL_b, YRI), C) D(Pop1, Nahua_P; 2417J_TOL_a, YRI) and D) D(Pop1, Nahua_T; 333B_TOL_a, YRI) . Expected tree topologies according to D value are drawn on the top of the plot, individual IDs in the trees are indicated with no suffixes. Red dots indicate significant deviations from D=0 (|Z|>3). Individuals from Sierra Gorda tend to have a significantly higher relationship with the present-day population used as Pop2 when Pop1 is a population from Northern or Southeast Mexico, but not when they are compared with other present-day populations from Central Mexico.

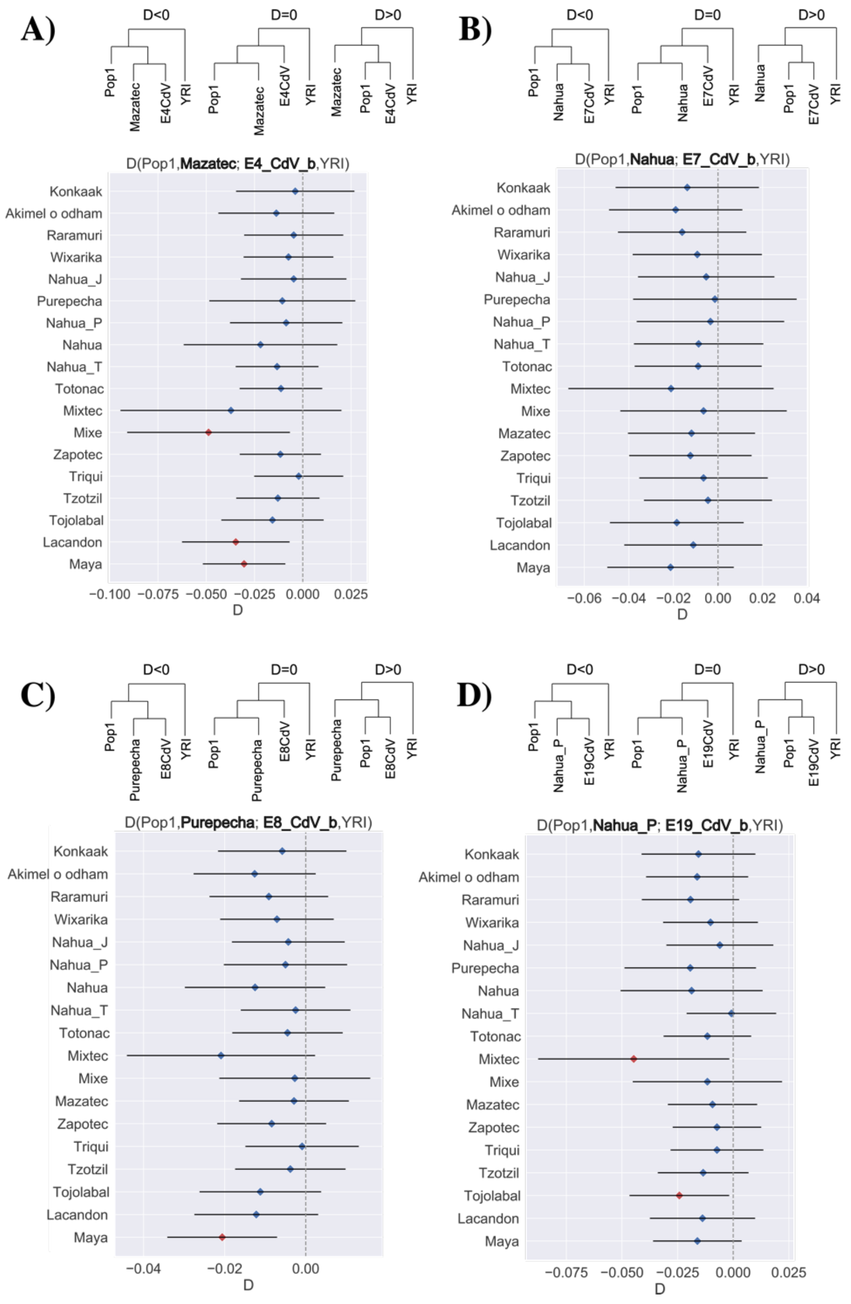


**Fig. S17.** D statistics for pre-Hispanic individuals from Cañada de la Virgen and present-day Indigenous populations. A) D(Pop1, Mazatec; E4_CdV_b, YRI), B) D(Pop1, Nahua; E7_CdV_b, YRI), C) D(Pop1, Purepecha; E8_CdV_b, YRI) and D) D(Pop1, Nahua_P; E19_CdV, YRI) . Expected tree topologies according to D value are drawn on the top of the plot, individual IDs in the trees are indicated with no suffixes. Red dots indicate significant deviations from D=0 (|Z|>3). Individuals from Cañada de la Virgen seem to be equally related to all present-day Indigenous populations except when Pop1 is Mixtec, Mixe or Southeastern populations, where they show a significant higher relationship with Pop2 .

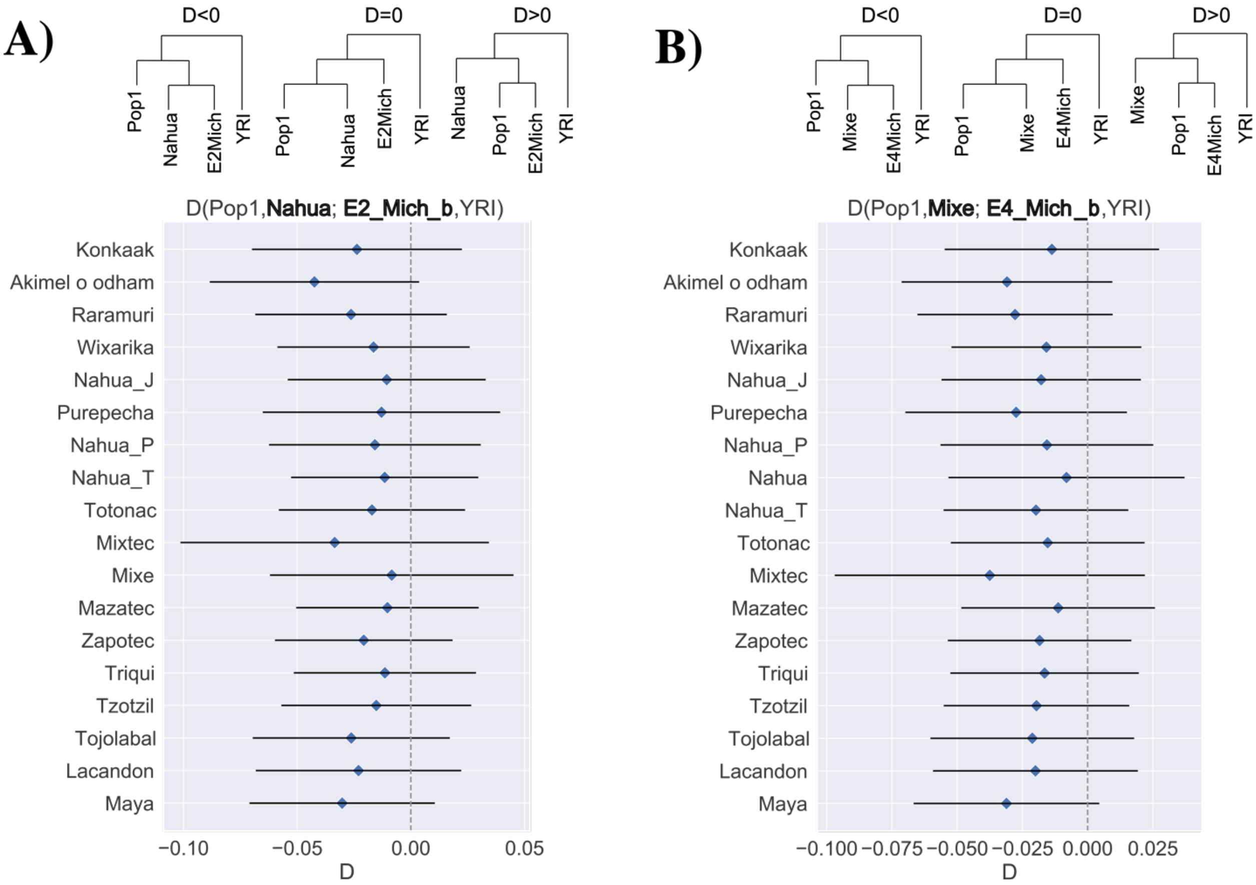


**Fig. S18.** D statistics for pre-Hispanic individuals from Michoacán and present-day Indigenous populations. A) D(Pop1, Nahua; E2_Mich_b, YRI), B) D(Pop1, Mixe; E4_Mich_b, YRI). Expected tree topologies according to D value are drawn on the top of the plot, individual IDs in the trees are indicated with no suffixes. Red dots indicate significant deviations from D=0 (|Z|>3). Individuals from Michoacán seem to be equally related to all present-day Indigenous populations.

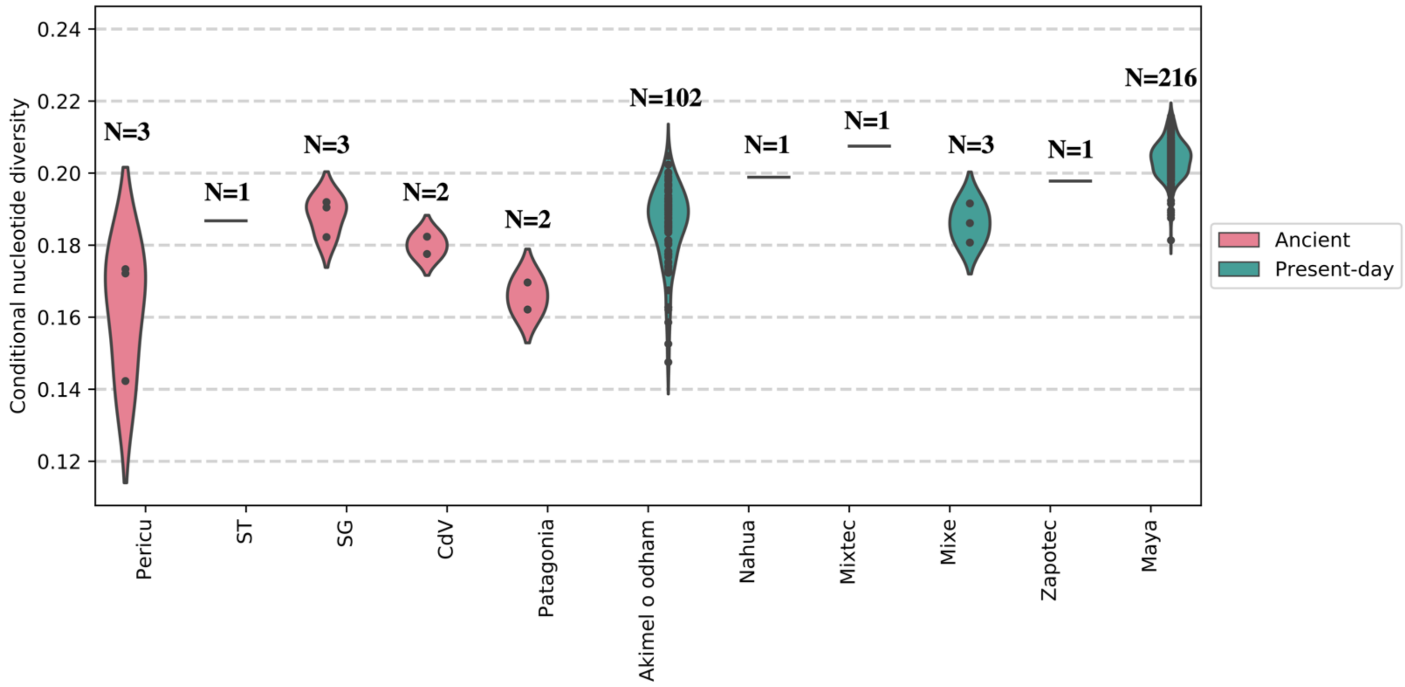


**Fig. S19**. Conditional Nucleotide Diversity. Violin plot indicating the CND values estimated by pairs of individuals from the same archaeological site or population. Ancient pre-Hispanic populations are indicated in pink while present-day populations are in green. Up to each violin plot are indicated the number of comparisons (CND values estimated) for each population or archaeological site.

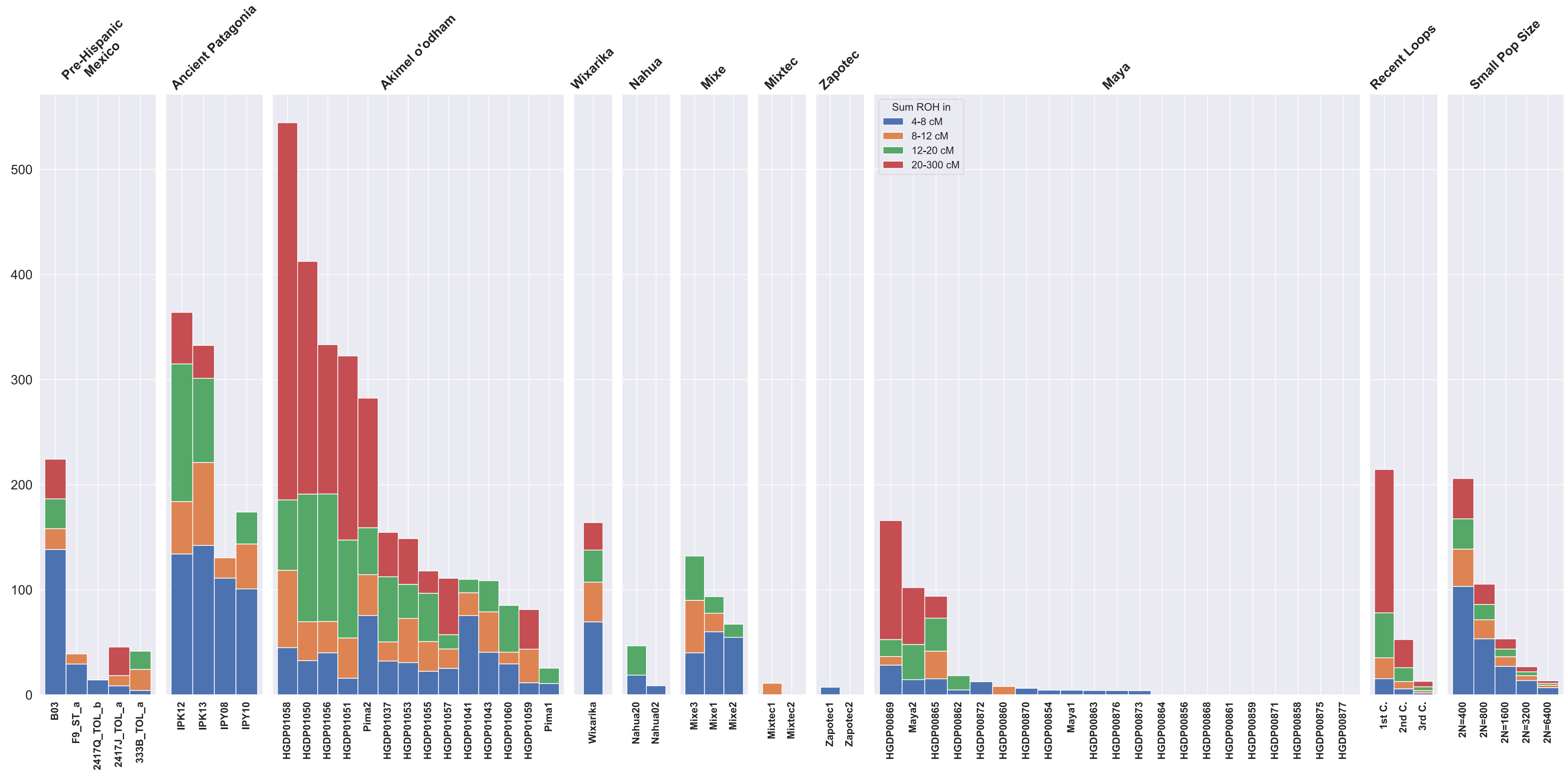


**Fig. S20.** Analysis of runs of homozygosity in pre-Hispanic and present-day Indigenous individuals from Mexico. Four pre-Hispanic individuals from Patagonia were analyzed for comparison. Length of ROH are shown in different colors. Pre-Hispanic individual B03 (Pericú) and the present-day Akimel O'odham population from Northern Mexico tend have the longer ROH from their time. Pre-Hispanic individuals from Patagonia have longer ROH than pre-Hispanic individuals from Mexico. Recent loops is an estimation of the ROH expected when an individual is a descendant of 1st cousins, 2nd cousins and 3rd cousins. Small Pop Size refers to the expected values in individuals from small populations.

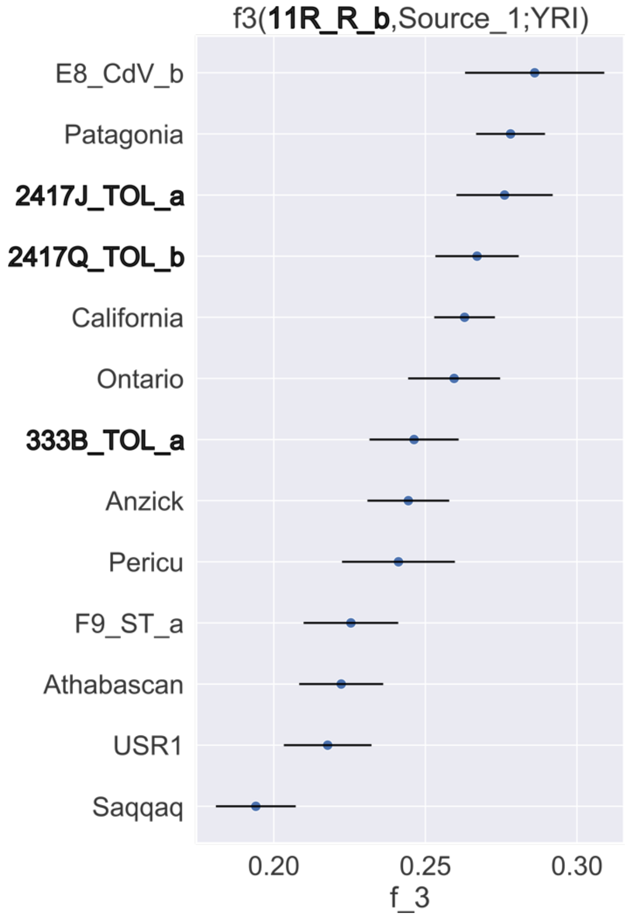


**Fig. S21.** Outgroups f3 statistics for pre-Hispanic individual 11R_R_b from Sierra Gorda and pre-Hispanic individuals. Individuals from Sierra Gorda are shown in bold. Higher values of f3 indicate higher shared genetic drift. Point estimates and one standard error are shown. The individual 11R_R_b seems to have a higher genetic drift shared with the individual E8_CdV_b from Cañada de la Virgen and not with the individuals from Sierra Gorda.

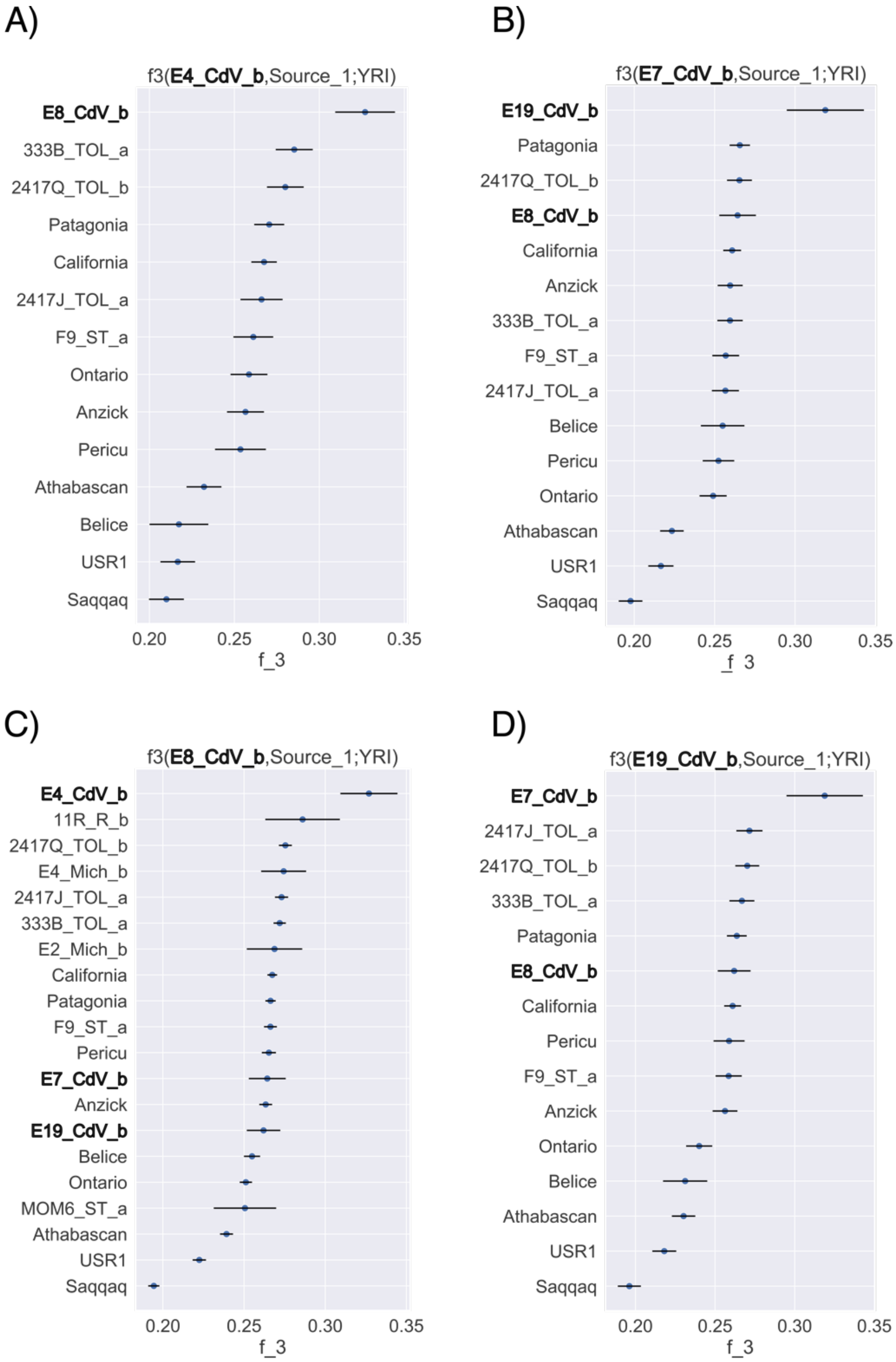


**Fig. S22**. F3 outgroup statistics for pre-Hispanic individuals from Cañada de la Virgen site. A) E4_CdV_b, B) E7_CdV_b, C) E8_CdV_b, and D) E19_CdV_b compared with pre-Hispanic individuals from Mexico and America. Higher values of f3 indicate higher shared genetic drift. Point estimates and one standard error are shown. Individuals E8_CdV_b and E4_CdV_b are more similar to each other than to any other ancient Native American, consistent with them being second-degree relatives as revealed by a relatedness test. The same happens for E7_CdV_b and E19_CdV_b. Intriguingly, f3 values of individuals from Cañada de la Virgen are not consistently the highest with other unrelated individuals from Cañada de la Virgen.

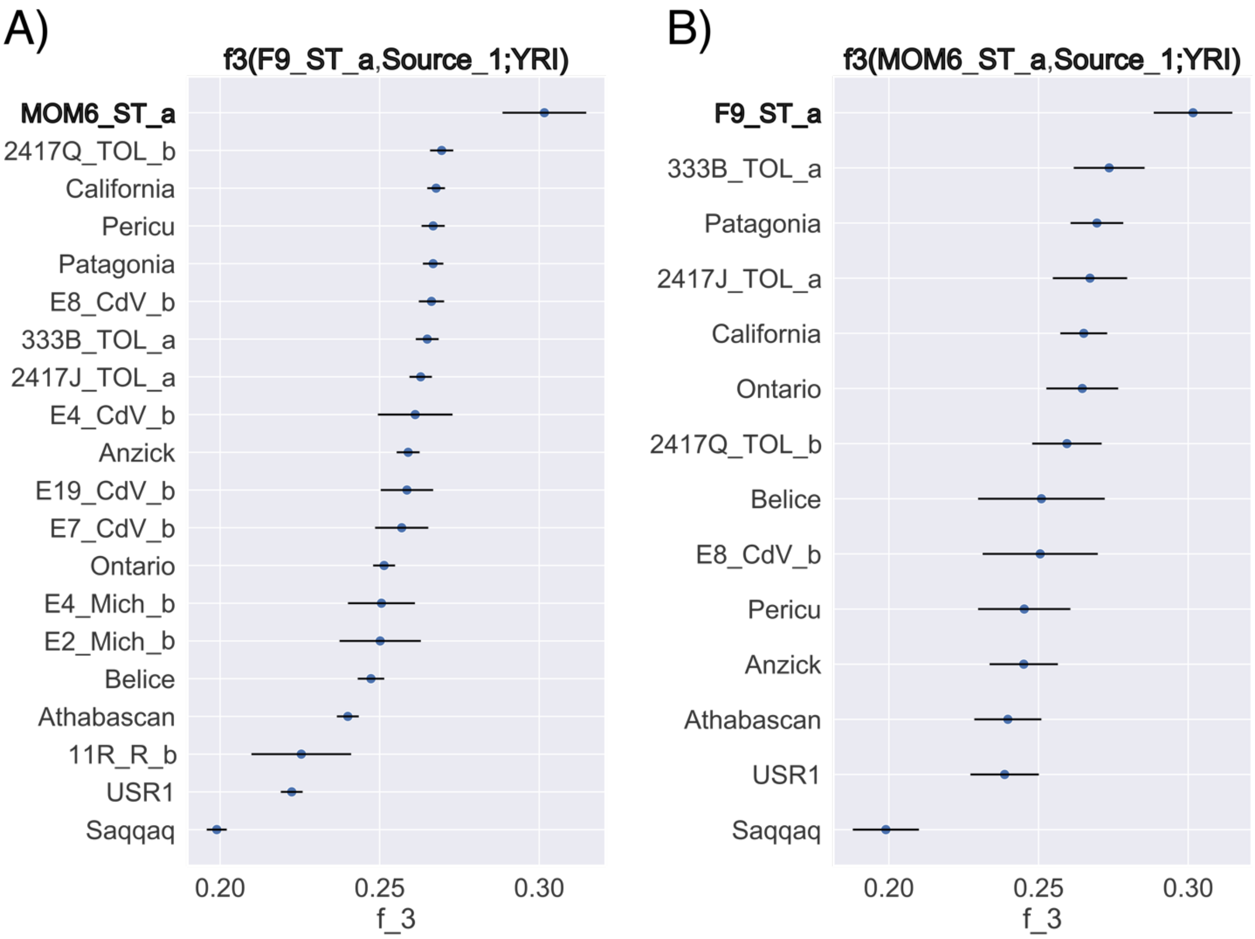


**Fig. S23.** Outgroups f3 statistics for pre-Hispanic individuals from Sierra Tarahumara and other ancient Native Americans. A) Individual F9_ST_a and B) Mummy MOM6_ST_a. Higher values of f3 indicate higher shared genetic drift. In both cases, we observe that F9 and MOM6 are mutually more similar to each other than to any other Ancient Native American. Individuals from Sierra Tarahumara are highlighted in bold.

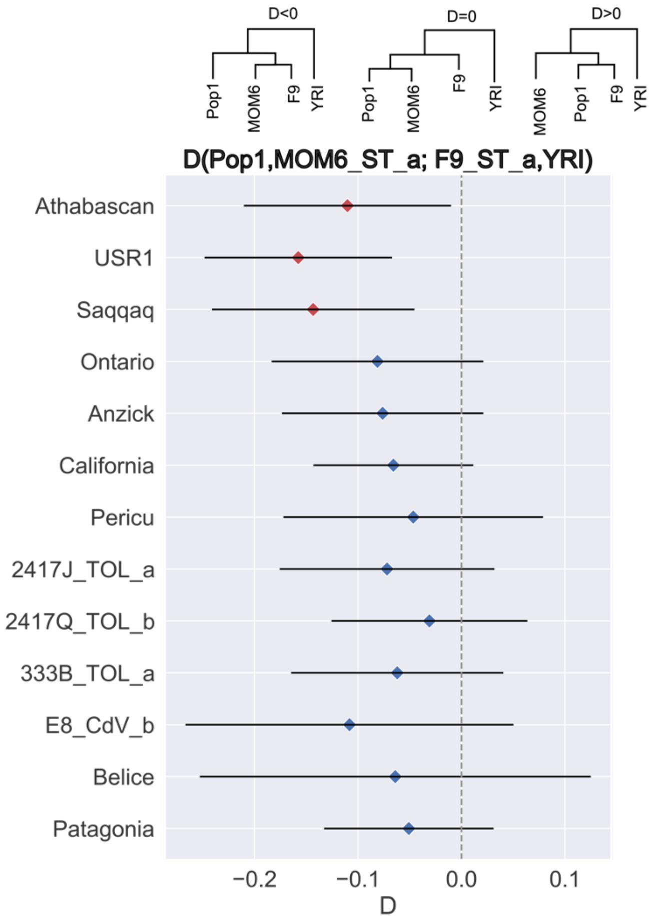


**Fig. S24.** D statistics of pre-Hispanic individuals from North-Mexico in the form D (Pop1, MOM6; F9, YRI). Pop1 is any of the other pre-Hispanic individuals from Mexico or America and shown in the Y axis. Expected tree topologies according to different D values are drawn on the top of the plot. Individual IDs in the trees are indicated with no suffixes. Red dots indicate significant deviations from D=0 (|Z|>3). MOM6_ST_a and F9_ST_a are significantly closer to each other only when they are compared with Athabascan, USR1, and Saqqaq individuals.

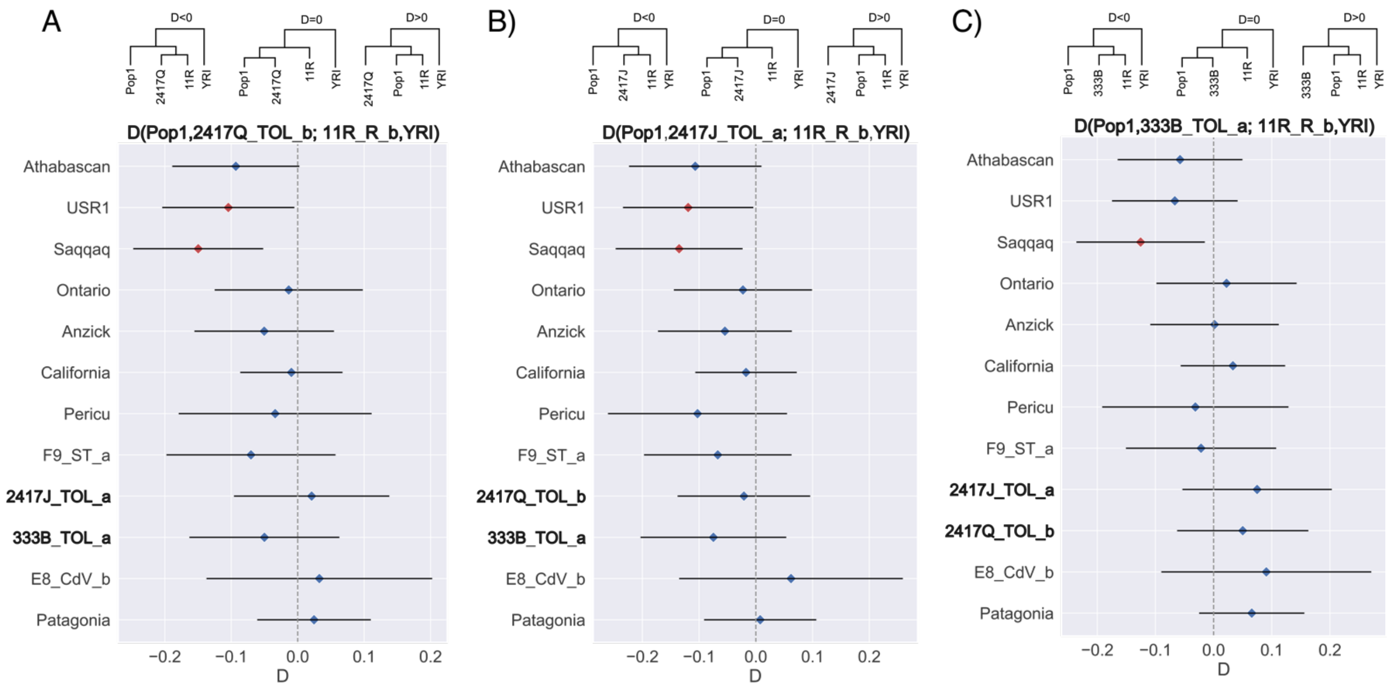


**Fig. S25**. D statistic values between the pre-Hispanic individual 11R_R_b from Central-Mexico, Sierra Gorda in the form D (Pop1,Pop2; Test, YRI). Test is individual 11R_R_b and Pop2 are any individual from Sierra Gorda, Pop1 are any of the other ancient Native Americans from Mexico and elsewhere and are shown in the Y axis. Expected tree topologies according to D value are drawn on the top of the plot, individual IDs in the trees are indicated with no suffixes. Red dots indicate significant deviations from D=0 (|Z|>3). Individuals from Sierra Gorda are highlighted in bold. A) Dstats in the form D (Pop1,2417Q_TOL_b; 11R_R_b, YRI). B) Dstats in the form D (Pop1,2417J_TOL_a; 11R_R_b, YRI). C) Dstats in the form D (Pop1,333B_TOL_a; 11R_R_b, YRI). Error bars in D (Pop1, X_TOL_x; 11R_R_b, YRI) are big and do not show a higher relationship between the individual 11R_R_b from Ranas and the other individuals from the Sierra Gorda (Toluquilla).

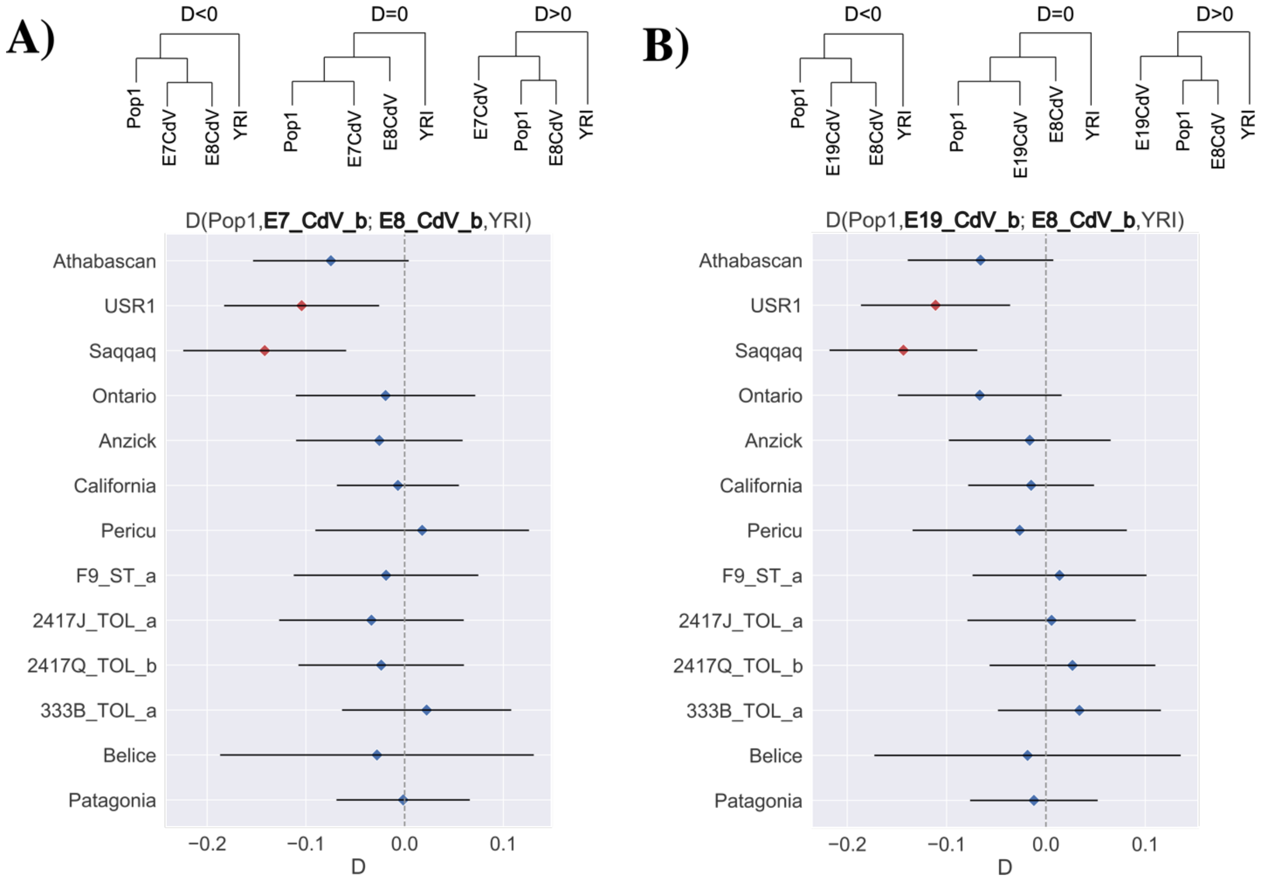


**Fig. S26**. D statistic values between pre-Hispanic individuals from Cañada de la Virgen, in the form D (Pop1,Pop2; Test, YRI). Pop2 and Test are any non-related individual from Cañada de la Virgen, Pop1 are any of the other ancient Native Americans from Mexico and elsewhere and are shown in the Y axis. Expected tree topologies according to D value are drawn on the top of the plot, individual IDs in the trees are indicated with no suffixes. Red dots indicate significant deviations from D=0 (|Z|>3). Individuals from Guanajuato are highlighted in bold. Overlapping sites between E7_CdV_b, E8_CdV_b and E19_CdV_b where <1,000 SNPs, thus plots only include: A) Dstats in the form D (Pop1, E7_CdV_b; E8_CdV_b, YRI) and, B) Dstats in the form D (Pop1, E19_CdV_b; E8_CdV_b, YRI). Individuals from Cañada de la Virgen do not show a significantly higher relationship than with any other ancient individual from the Americas, except when compared with USR1 and Saqqaq.

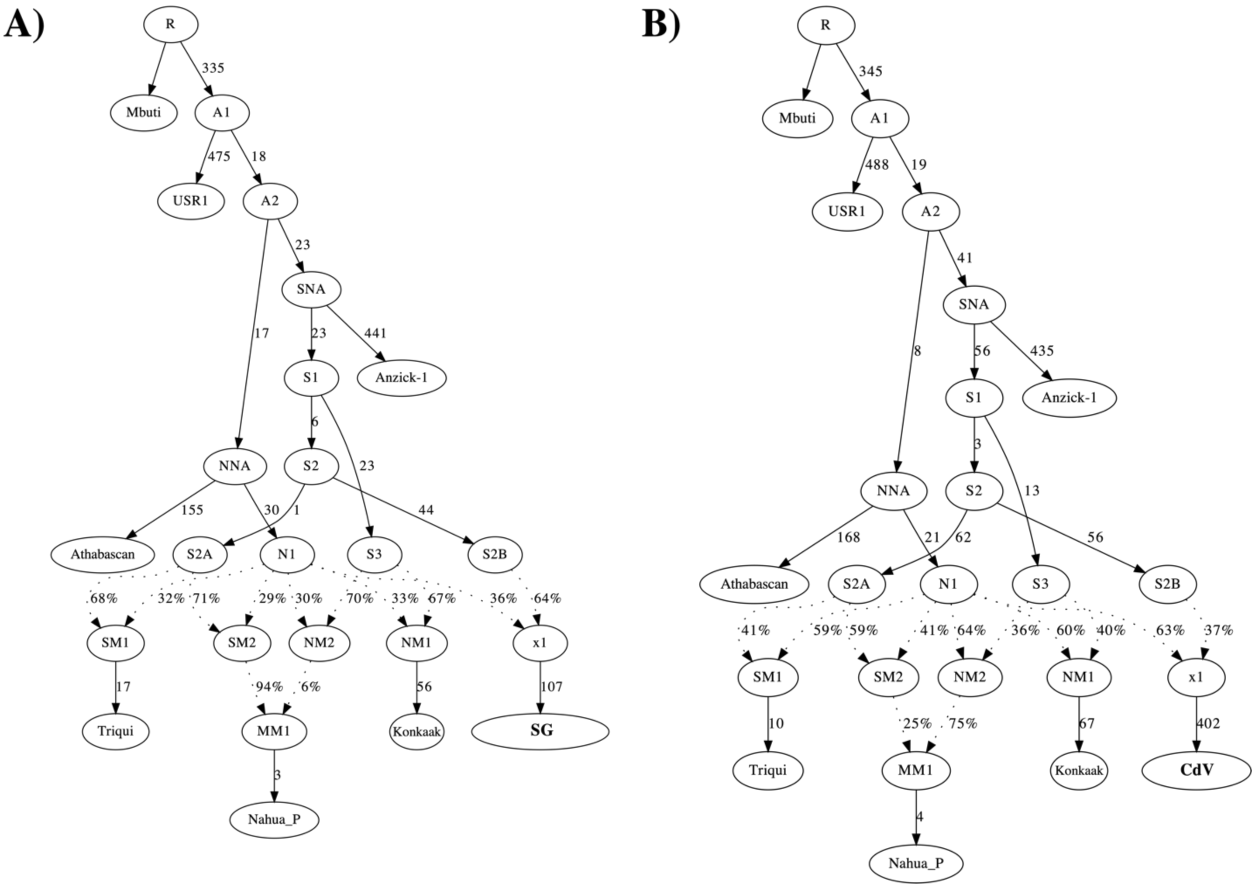


**Fig. S27**. Ancestry proportions for the individuals from Central Mexico, analyzed through qpGraph. Base model includes Anzick as a reference for the SNA and Athabascan as a references for the NNA branch. Konkaak was used a proxy for Northern Indigenous population from Mexico, Triqui as South proxy and Nahua_P as Central Mexico proxy. A) Model for SG individuals, it includes the genetic information of 2417Q_TOL_b, 2417J_TOL_a and, 333B_TOL_a. Tree with |Z-score| of 1.319. B) Model for CdV individuals, it includes the genetic information of E7_CdV_b, E8_CdV_b and E19_CdV_b. Individuals from the SG and CdV shows very different proportions of S2B and N1 ancestries. Tree with |Z-score| of 2.821. All f statistics are within 1.75 standard error.

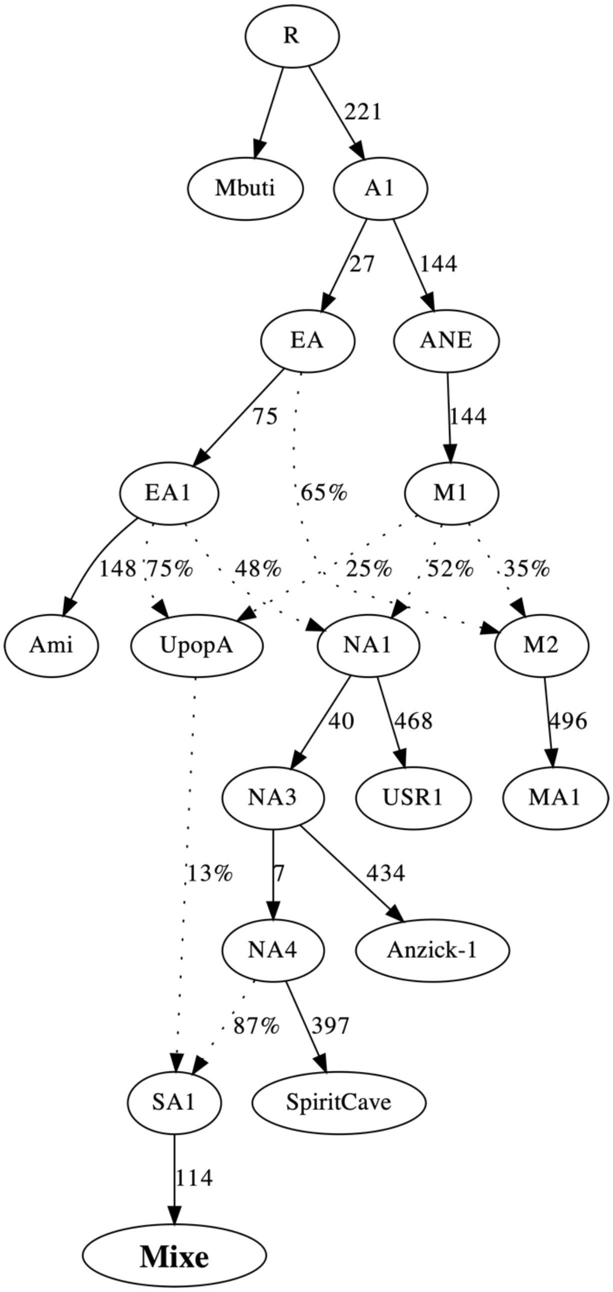


**Fig. S28**. Presence of ghost population ancestry in present-day Mixe population, analyzed through qpGraph. Base model includes Mbuti as an outgroup, Ami as the East Asian, MA1 as the ancient North Eurasian, USR1 as the ancient Beringian, Anzick-1 and Spirit Cave as the NNA. Model shows a 13% of the ghost population UpopA in the present-day Mixe. Tree with |Z-score| of 2.033. All statistics are within 1.75 standard error.
